# The fungal gene cluster for biosynthesis of the antibacterial agent viriditoxin

**DOI:** 10.1101/601716

**Authors:** Andrew S. Urquhart, Jinyu Hu, Yit-Heng Chooi, Alexander Idnurm

**Affiliations:** School of BioSciences, University of Melbourne, Australia; School of Molecular Sciences, University of Western Australia, Australia

**Keywords:** Atropisomer, Eurotiales, gene cluster, laccase, polyketide synthase

## Abstract

**Background:** Viriditoxin is one of the ‘classical’ secondary metabolites produced by fungi and that has antibacterial and other activities; however, the mechanism of its biosynthesis has remained unknown.

**Results:** Here, a gene cluster responsible for its synthesis was identified, using bioinformatic approaches from two species that produce viriditoxin and then through gene disruption and metabolite profiling. All eight genes in the cluster in *Paecilomyces variotii* were mutated, revealing their roles in the synthesis of this molecule and establishing its biosynthetic pathway which includes an interesting Baeyer-Villiger monooxygenase catalyzed reaction. Additionally, a candidate catalytically-inactive hydrolase was identified as being required for the stereoselective biosynthesis of (*M*)-viriditoxin. The localization of two proteins were assessed by fusing these proteins to green fluorescent protein, revealing that at least two intracellular structures are involved in the compartmentalization of the synthesis steps of this metabolite.

**Conclusions:** The full pathway for synthesis of viriditoxin was established by a combination of genomics, bioinformatics, gene disruption and chemical analysis processes. Hence, this work reveals the basis for the synthesis of an understudied class of fungal secondary metabolites and provides a new model species for understanding the synthesis of biaryl compounds with a chiral axis.

## Background

Fungi produce a diverse array of polyketide-derived biaryl compounds with biological activities that are of interest as pharmaceutical lead molecules or through modulating fungal interactions with other species in the environment. Viriditoxin (Compound 1, Figure 1) is a naphtho-α-pyrone produced by the Eurotiales fungi *Aspergillus viridinutans* and *Paecilomyces variotii*, and in limited amounts by *Aspergillus brevipes* [1–3]. Related biaryl molecules have also been identified from other fungi including *Fusarium* spp. (aurofusarin [4]), *Penicillium* sp. (rugulotrosin A and B [5]) and *Parastagonospora nodorum* (elsinochrome A [6]) (Figure 1).

**Fig. 1.**
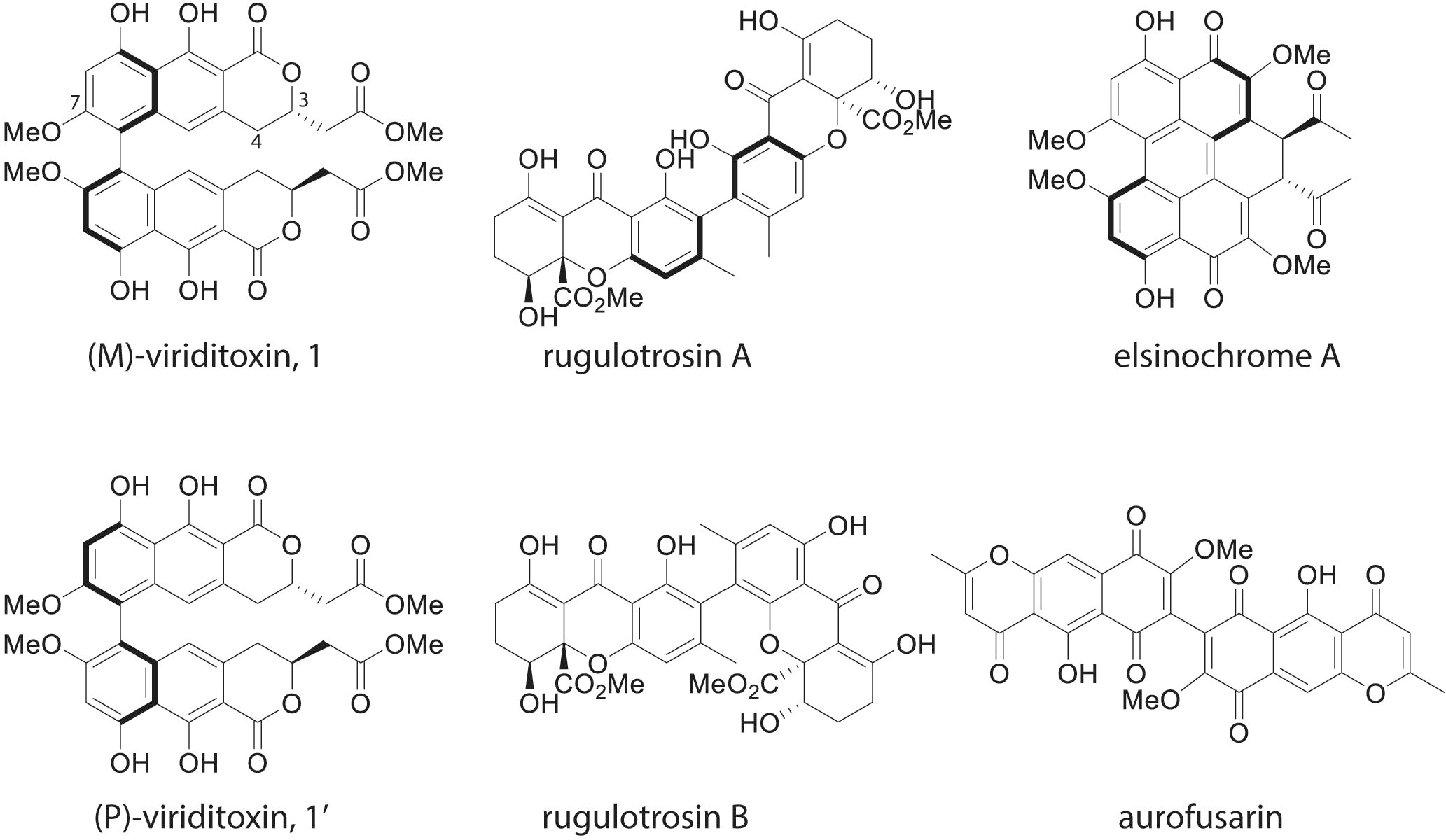
Structures of (*M*)-viriditoxin and (*P*)-viriditoxin (**1** and **1′**), as well as related metabolites produced by other fungi. The “*M*” and “*P*” refer to the atropisomer confirmations. The helical configuration is illustrated by the bold lines. **1** is the major form produced by *A. viridinutans* and *P. variotii.* Carbon numbers C3, C4 and C7 are indicated.

Viriditoxin **1** exhibits interesting biological activities. In particular, viriditoxin was identified as a potent inhibitor of bacterial FtsZ, a protein that is required for bacterial cell division [7]. Subsequent work demonstrated that this was due to decreased transmembrane potential and perturbing membrane permeability, which prevents membrane proteins including FtsZ from effectively binding to the membrane [8]. Viriditoxin is also active against cancer cells lines and is a proposed starting molecule for possible therapeutic applications [9, 10]. It has also been examined for a potential role in protection of sheep against blowflies [11]. In natural settings viriditoxin likely plays a role in competition against other microbes. For example, a strain of *P. variotii* was isolated as an endophyte from mangroves, and its antagonistic activities against bacteria attributed to production of viriditoxin [12].

From the perspective of biochemical synthesis, viriditoxin is of interest as it is a biaryl compound with axial chirality, in which rotation about the biaryl axis is hindered, resulting in two different stable forms which are known as atropisomers. The two atropisomers are referred to as *M*-viriditoxin **1** and *P*-viriditoxin **1′** with the “*M*” stereoisomer **1** being the major natural product found in *P. variotii* and *A. viridinutans* ([13], Figure 1). The formation of such compounds requires the coupling of two subunits. How this coupling occurs in fungi is only beginning to be understood. It is known that both P450 monooxygenases and laccases/multicopper oxidases are able to catalyze this coupling. While P450 monooxygenases have been shown to be capable of stereoselective coupling [14], none of the fungal laccase/multicopper oxidases enzymes which have been implicated in biaryl coupling have demonstrated stereospecificity [4, 15, 16]. In plants, laccases are able to catalyze stereoselective coupling in combination with dirigent proteins, however, no such proteins have been reported in fungi [17]. Despite the growing appreciation of the importance of atropisomers in the pharmaceutical industry, it is still not understood how laccase-catalyzed reactions can result in stereoselective biosynthesis of biaryl compounds in fungi [18, 19].

The genomes of two isolates of *P. variotii* were recently sequenced [20], opening opportunities for identifying the basis for the synthesis of viriditoxin and other metabolites that this fungus may produce. In this study, we identify a gene cluster for the synthesis of viriditoxin and define the genetic components for its synthesis. As part of this work we generated a draft genome of the original isolate that produces viriditoxin, *A. viridinutans*, and find a similar gene cluster in the related Eurotiales species, *P. variotii*. Gene manipulation tools are not yet available for *A. viridinutans*. Hence, the gene cluster was functionally characterized in *P. variotii*, a newly-emerging model fungal species where methods for gene manipulation were recently developed [20].

## Results and Discussion

### Identification of a putative viriditoxin biosynthesis cluster by comparative genomics between *P. variotii* and *A. viridinutans*

We recently sequenced the genome of *P. variotii* CBS 101075 which is known to produce viriditoxin [2] and a second *P. variotii* strain, CBS 144490 [20]. For comparison, the *A. viridinutans* strain FRR 0576 genome was sequenced using Illumina HiSeq2500 paired-end reads. 14,757,842 reads were generated and assembled into 285 contigs, totaling approximately 30 Mb (Table S1). The assembly and raw reads are available from GenBank under BioProject PRJNA513223. A curious commonality between *A. viridinutans* and *P. variotii* is that both genomes have a bimodal GC content, which in *P. variotii* has been attributed to active repeat inducted point (RIP) mutation [20].

The structure of viriditoxin is similar to a number of other fungal polyketide secondary metabolites, such as aurofusarin and the α-pyrone elsinochrome C, whose synthesis requires a polyketide synthetase (PKS) enzyme. Hence, candidate genes encoding PKSs were sought using BLAST with the protein sequences of the PKS enzymes PKS12 (which is required for aurofusarin biosynthesis [21], and ElcA (which is required for elsinochrome C biosynthesis [22]). Two candidate PKS enzymes were identified in the CBS 101075 genome: protein IDs 480069 (named VdtA) and 456077 (named PvpP), available through MycoCosm [20, 23].

BLAST comparisons of the *A. viridinutans* PKS to these two *P. variotii* PKS genes revealed a putative candidate cluster for viriditoxin (hereafter referred to as the *vdt* cluster). This cluster consists of nine genes in *P. variotii* and eight in *A. viridinutans*, with the difference being due to the presence of a second putative PKS-encoding gene (*vdtX*) in *P. variotii* (Figure 2A, Table S2).

While we were preparing this manuscript, a parallel genomic analysis was conducted by Fürtges et al., who proposed the same biosynthetic cluster in *A. viridinutans* [15]. However, the role of the genes in the cluster was not experimentally validated beyond demonstration of enzymatic activity of the recombinantly-expressed laccase enzyme on a non-native substrate. Thus, the gene cluster was not conclusively linked to viriditoxin biosynthesis.

### Transcription factor VdtR regulates expression of the *vdt* gene cluster

Fungal secondary metabolite clusters may contain a transcription factor responsible for regulation of the cluster [24]. VdtR has the characteristics of a six-cysteine (C6) zinc cluster type of transcription factor, with similarity to AflR of the aflatoxin biosynthesis cluster in *Aspergillus* species. The *vdtR* gene was deleted, and the expression of the genes in the gene cluster compared between the wild type and *vdtR* deletion strains. This revealed reduced transcript levels of genes in the *vdt* cluster (Figure 2B), confirming the identity of VdtR as a regulator of the genes in the cluster. This approach was also used to help define the boundaries of the cluster: one of the flanking genes (protein ID423248) was unaffected while expression of the other flanking gene (protein ID480071) could not be detected by qPCR (data not shown).

**Fig. 2.**
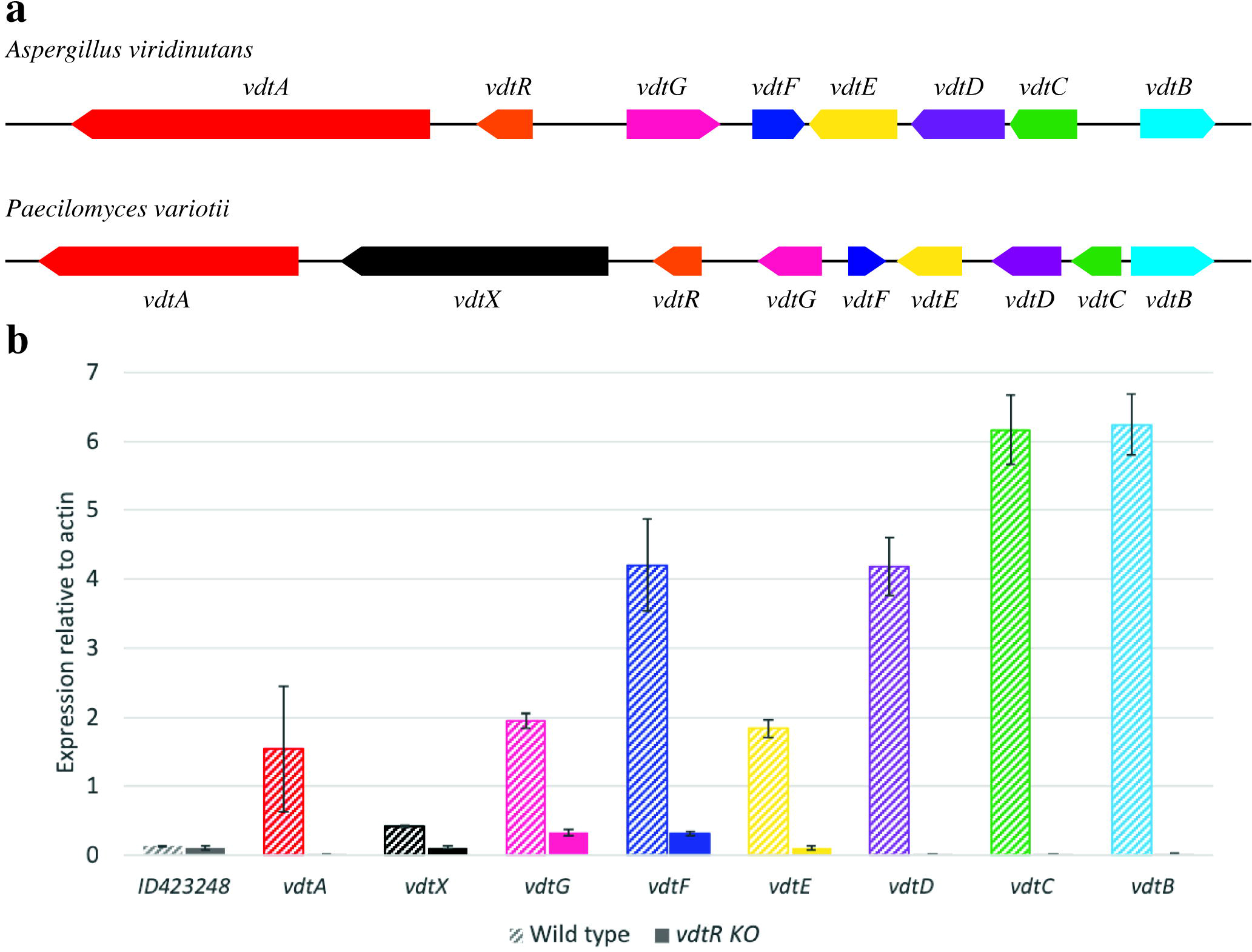
Putative *vdt* cluster encoding viriditoxin **1** biosynthesis. **a.** Comparison between the *A. viridinutans* and *P. variotii* clusters. **b.** Comparison of the transcript levels of *vdt* genes between wild type and *vdtR* deletion mutant measured by qPCR.

### The polyketide synthase VdtA is required for the first step of viriditoxin biosynthesis

*P. variotii* encodes two candidate PKSs that could be involved in the synthesis of heptaketides similar to YWA1, a precursor of both aurofusarin and pigments – ID480069 (VdtA) and ID456077 (PvpP). Both PKS genes were replaced with a hygromycin resistance marker through homologous recombination. The deletion of *vdtA*, in both *P. variotii* strains CBS 101075 and CBS 144490, or the associated transcription factor *vdtR* in CBS144490, abolished the production of **1** and related derivatives **2** and **3** in the culture filtrates of the strains as detected by LC-DAD-MS, confirming its role early in the biosynthesis pathway (Figure 3). Structures of these three compounds were confirmed by 1D and 2D NMR analysis. Electronic circular dichroism (ECM) spectroscopy was used to determine the helical confirmation of the compounds by comparing to a commercial standard of **1** (Tables S3, S5, S6, Figures S1-S8, S15-S26)

**Fig. 3.**
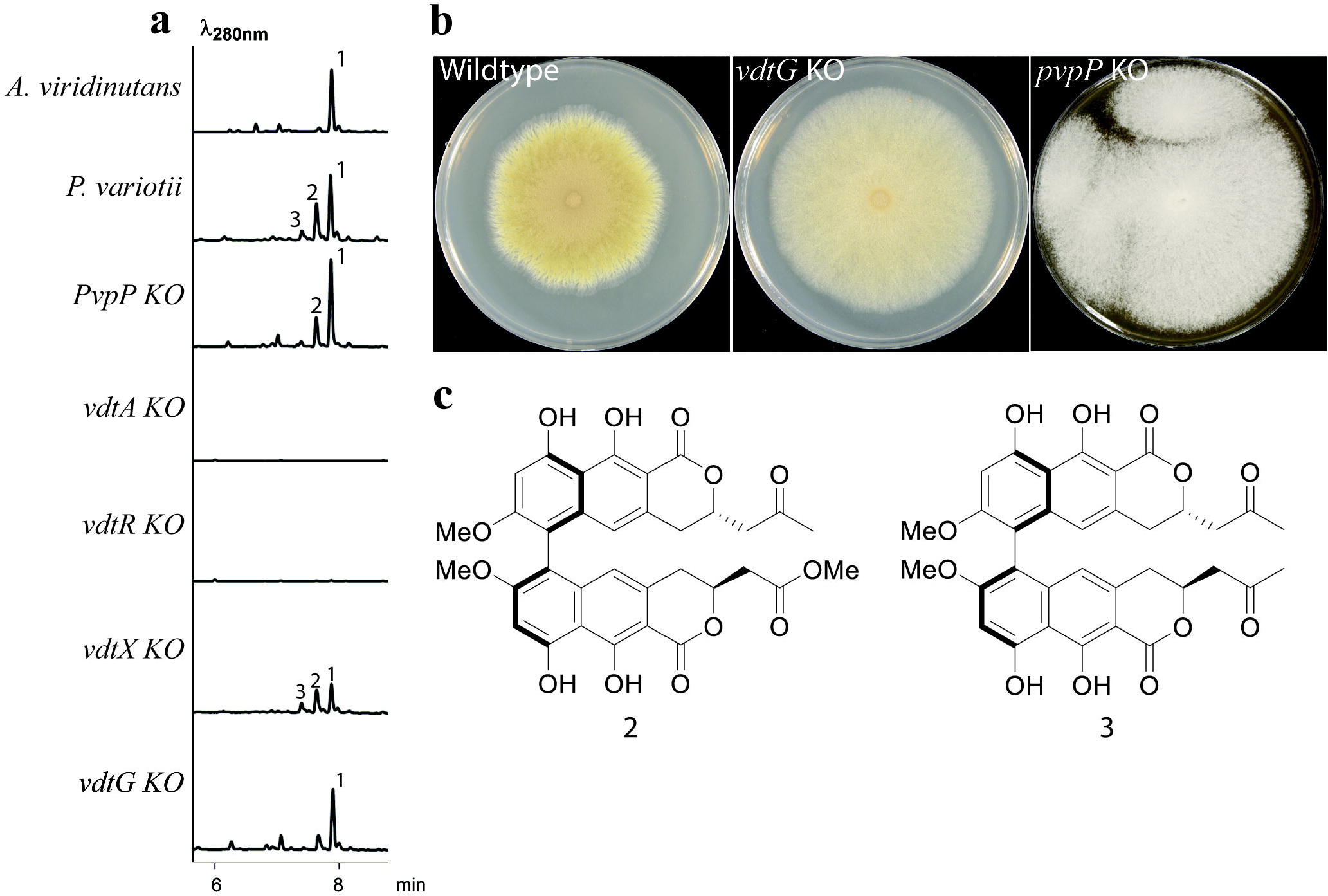
**a.** LC-DAD-MS analysis of *A. viridinutans* and *P. variotii* CBS 144490 and *P. variotii* mutants (*pvpP, vdtA, vdtR, vdtX and vdtG*). **b.** Deletion of *vdtG* resulted in a sporulation defects and deletion of *pvpP* resulted in a loss of conidial pigmentation. **c.** Structures of the additional viriditoxin derivatives (**2** and **3**) produced by the wild type *P. variotii* strain CBS 144490.

The deletion of the *pvpP* gene in strain CBS 101075 altered the pigmentation of the strains but did not alter viriditoxin synthesis (Figure 3). Comparison to the biosynthesis pathway for the pigment DHN melanin in *Aspergillus fumigatus* shows that this protein is homologous to PksP that is responsible for the synthesis of the yellow molecule YWA1 [25]. BLAST searches with the DHN melanin pathway genes in *A. fumigatus* [26, 27] revealed their absence from the *P. variotii* genome.

The *vdt* gene cluster in *P. variotii* encodes a second putative PKS that is absent from the *A. viridinutans* genome. Hence, one hypothesis is that this gene is not required for viriditoxin production. Disruption of this gene (*vdtX*) via homologous recombination was unsuccessful, so we employed an alternative approach using repeat induced point mutation, which is an active process in *P. variotii* during heterothallic sexual crosses [20]. RIP is a fungal-specific mechanism in which duplicated DNA is recognized and specifically mutated as part of the sexual cycle, and has been used to make targeted mutations in genes in *Neurospora crassa* [28]. To achieve this, a second partial copy of *vdtX* was introduced into *P. variotii* CBS 101075, and that strain crossed to the wild type strain of opposite mating type (CBS 101075) such that RIP can mutate the duplicated DNA, and the identification of progeny carrying mutations. A progeny was obtained in which the native copy of the gene had been heavily mutated by RIP, introducing premature stop codons (Supplementary Figure 1). The ability to generate mutants efficiently through RIP adds another technique to the repertoire available in *P. variotii*. The *vdtX* RIP mutant continued to produce viriditoxin, indicating that VdtX is not required viriditoxin biosynthesis (Figure 3). The variation between the RIP mutant and wild type might be due to the variability expected in the progeny of a sexual cross. As such, this gene was not considered further in regard to viriditoxin biosynthesis but might possibly play a role in the production of additional metabolites not extracted/detected by the techniques employed in this study.

### The PKS VdtA is localized within a specialized structure

The localization of PKS enzymes in fungal secondary metabolite synthesis is largely unknown. The PksP (Alb1) PKS of *Aspergillus* species involved in the production of melanin is localized to the endosome [29], and it represents the only fungal PKS with a demonstrated subcellular localization. A strain expressing a VdtA-GFP fusion protein displayed green fluorescence tightly confined to small circular structures within the cells (Figure 4A). Expression of GFP alone using PLAU17 [30] localizes to the cytosol (data not shown). The VdtA-GFP strains continued to produce viriditoxin, indicating that the addition of GFP to its carboxyl-terminus end has not destroyed the activity of the enzyme (data not shown) and that the structures are not an artefact from gene manipulation. These small spherical structures are reminiscent of peroxisomes. Peroxisomes have been implicated in the production of other fungal secondary metabolites [31]. In particular, the fluorescent aflatoxin precursor norsolorinic acid, produced by PksA, is localized to the peroxisome [32]. However, the PKS enzyme itself has not been localized. Given sequence similarity between the aflatoxin-producing PksA and *P. variotii* VdtA, we explored the possibility that the unusual localization pattern observed for VdtA-GFP represented peroxisome localization by dual-labelling with an mCherry-SKL peroxisomal construct. The addition of the three amino acid motif SKL to the carboxy terminal is widely used to localize fluorescent proteins to the peroxisome [33]. However, VdtA-GFP and mCherry-SKL showed clearly distinct patterns of localization. Based on previous analysis of nuclei and mitochondria using tagged proteins we can exclude those organelles [20]. Thus, these structures where VdtA is targeted remain unknown.

**Fig. 4.**
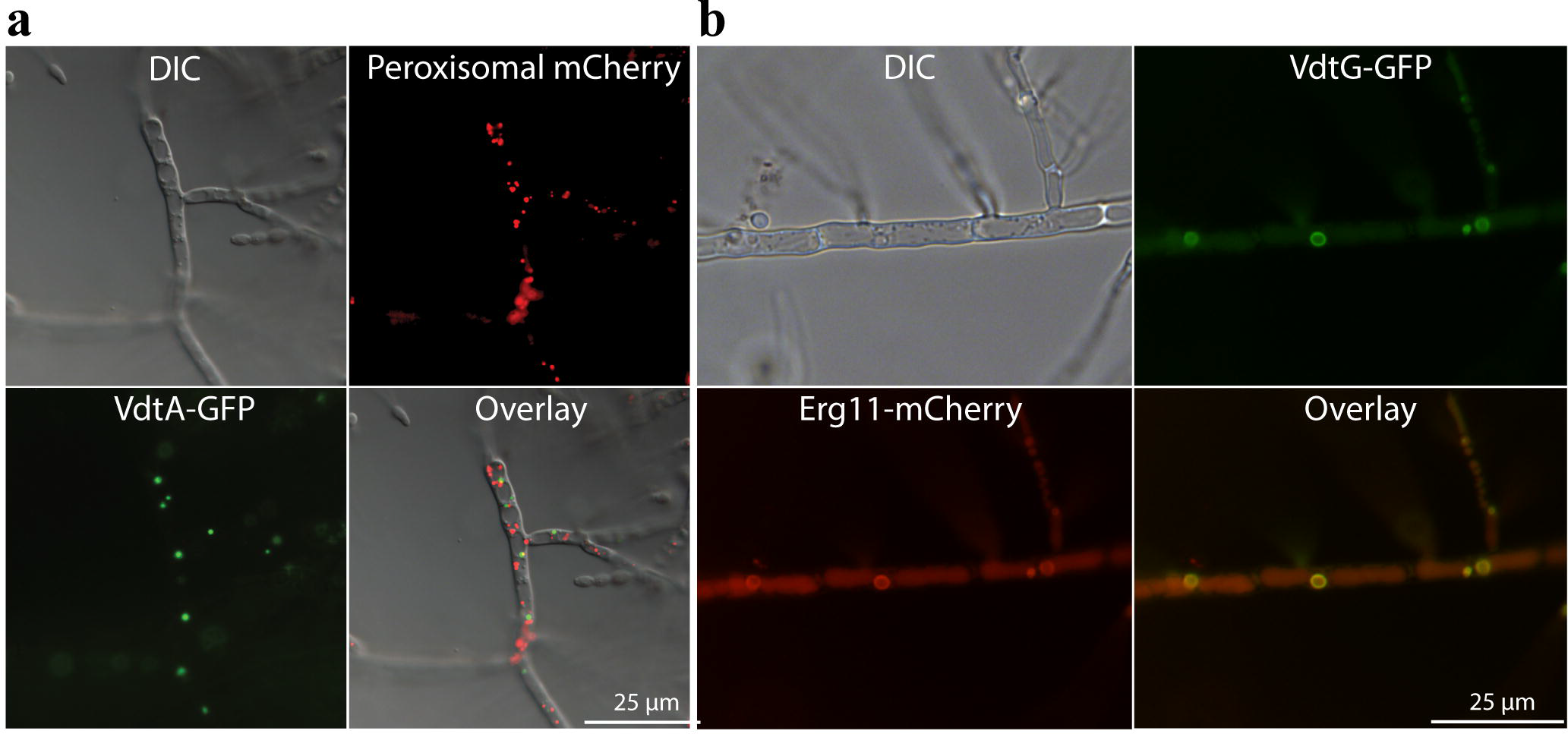
Viriditoxin is synthesized in different organelles. **a.** VdtA-GFP fusion protein localizes to small spherical structures within hyphae, which are distinct from the peroxisomes (labelled with mCherry-SKL). **b.** VdtG-GFP localized around a small circular structure and co-localizes with the endoplasmic reticulum marker Erg11-mCherry.

### The transporter VdtG is localized on an internal membrane and is not essential for viriditoxin production

Deletion of the gene encoding a putative transporter VdtG had no effect on the production of viriditoxin (Figure 3A). However, the gene deletion strains have a sporulation defect (Figure 3B). Production of secondary metabolites is often developmentally timed with the beginning of spore formation [34], so we hypothesize that this phenotype is likely to result from toxicity effects beginning at this stage. Such morphological differences have not been observed in previously disrupted transporters associated with secondary metabolite gene clusters such as the genes *aflT* in *A. parasiticus* [35] or *sirA* in *Leptosphaeria maculans* [36], although in the case of SirA, sensitivity to sirodesmin was increased in the *sirA* mutants.

To further probe the role of this putative transporter, DNA encoding GFP was fused to the native *vdtG* gene of *P. variotii*. Strains expressing this fusion protein did not show the sporulation defects present in the *vdtG* KO strains, indicating that the fusion protein is functional (data not shown). Fluorescence microscopy revealed that VdtG is localized around an internal membrane structure and the septa (Figure 4B). Taken together these results lead us to hypothesize that VdtG is required to compartmentalize the synthesis of viriditoxin or its intermediates within the cell to limit auto-toxic effects.

The VdtG-GFP structure resembles the ‘toxisomes’ of *F. graminearum* [37] and ‘aflatoxisomes’ of *Aspergillus* species [38]. Toxisomes are proliferations of the smooth endoplasmic reticulum [39]. For this reason, we examined the co-localization of VdtG-GFP with a known endoplasmic reticulum protein, Erg11 (Cyp51) required for sterol synthesis, as an mCherry fusion [40]. This showed clear co-localization of the two proteins suggesting that the VdtG-GFP structure is part of the endoplasmic reticulum or an endoplasmic reticulum derived structure (Figure 4B). The localization of VdtG to these structures provides evidence that this structure might be homologous to toxisomes and aflatoxisomes. There is no evidence that the VdtG and VdtA structures co-localize, given that the VdtA structures are more numerous and of different size.

### VdtC, VdtF and VdtE are involved in tailoring of the polyketide precursor produced by VdtA to the monomer semi-viriditoxin

We then generated knockout mutants for the genes encoding putative tailoring enzymes VdtC, VdtF and VdtE. LC-DAD-MS analysis of the culture extracts revealed major changes in the metabolite profiles (Figure 5). The compounds were purified from the individual mutant and subjected to NMR analysis for elucidation of the structures of **3**, **4**, **5**, **7**, **8** and **9** (Table S6-S11, Figures S21-S56) whereas **6** is proposed based on mass (Figure S1). ECM spectroscopy was used to determine the helical confirmation of the compounds **3, 4, 5, 7, 8** and **9** (Table S6-S11). The structure of these compounds provides insights into the function of each of these enzymes.

**Fig. 5.**
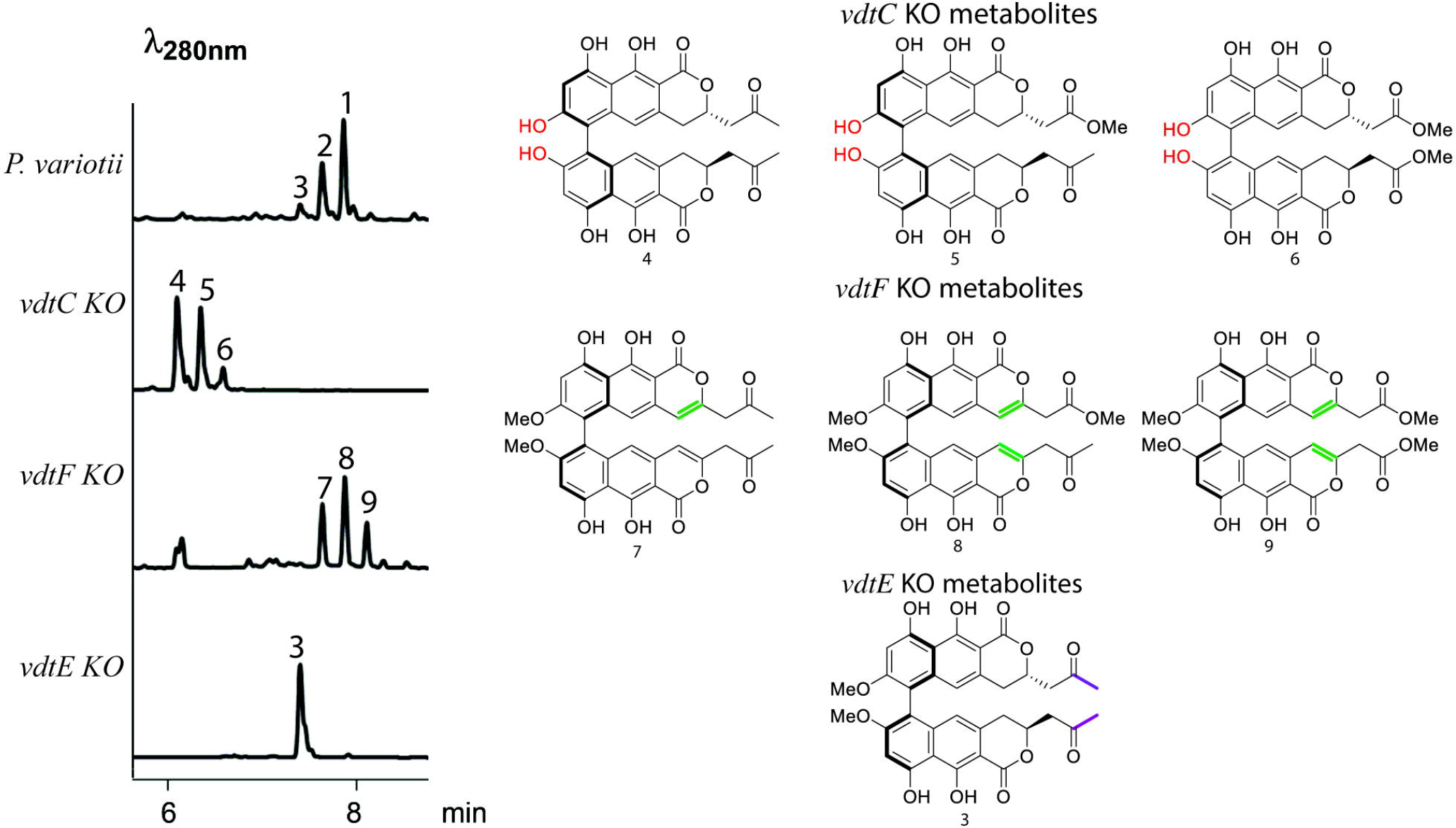
VdtC, VdtF and VdtE are required to tailor the hypothetical alpha pyrone monomer produced by VdtA into semi-viriditoxin. *vdtC*, *vdtF* and *vdtE* deletion strains all failed to make viriditoxin, instead producing a set of viriditoxin-related metabolites. Comparing the structure of the compounds produced by each deletion strain to **1** reveals the role of each tailoring enzyme (differences colored).

VdtC shows 44% similarity to the *O*-methyltranstrase AurJ in *F. graminearum* involved into the biosynthesis of aurofusarin [4]. As expected, deletion of *vdtC* results in the production of compounds **4**, **5** and **6** which differ from viriditoxin in that the hydroxyl group has not been methylated. Interestingly, circular dichroism data showed that the two metabolites purified in sufficient quantity for ECM analysis, **4** and **5** are both in the P helical form, suggesting that the *O*-methyl group is important for the stereoselective coupling of the monomers.

BLAST of the VdtF protein sequence revealed a number of close fungal homologs, but also shows 29% identity to the 3-oxoacyl-[acyl-carrier-protein] reductase FabG of bacterium *Eschericha coli* and thus may be able to catalyze a reduction reaction [41]. Deletion of *vdtF* results in the production of compounds **7**, **8** and **9**, which differ from **1** in that the C3-C4 double bond has not been reduced (Figure 5). We thus conclude that VdtF is responsible for the reduction of this bond.

BLAST against the Swiss-Prot database show that VdtE displays homology to a number of previously characterized Baeyer-Villiger monooxygenases (BVMOs). BVMOs are a group of enzymes responsible for the conversion of ketones into esters [42]. Consistent with the hypothesis that VdtE is a Baeyer-Villiger monooxygenase, deletion of *vdtE* resulted in the accumulation of compound **3** in which the ketone group has not been converted into the ester found in **1** (Figure 5). The activity of VdtE is similar to MoxY of the aflatoxin biosynthesis cluster found in *Aspergillus* spp. [43]. MoxY converts the ketone hydroxyversicolorone (HVN) into the acetate ester versiconal hemiacetal acetate (VHA). This is a rather unique reaction, so efforts towards purification of recombinant VdtE are ongoing to characterize further this unusual enzyme.

### The laccase VdtB is involved in dimerization while a non-catalytic hydrolase-like protein VdtD affects the stereochemistry outcome of the coupling

VdtB shows homology to laccase enzymes previously implicated in oxidative coupling, including GIP1 that is suggested to play a role in the dimerization of two rubrofusarin molecules to form aurofusarin in *F. graminearum* [4] and MCE that dimerizes monapinone A into dinapinone A in *Talaromyces pinophilus* [16]. Thus, we thus hypothesized that VdtB would be required for forming the biaryl bond between two semi-viriditoxin molecules to produce viriditoxin. Consistent with this hypothesis, disruption of the *vdtB* gene led to the accumulation of four major compounds, in which their *m/z* as detected by LC-MS corresponded to the molecular weight expected for monomers. The structure of three of these compounds **10**, **11** and **12** is predicted based on mass (S1). This confirms that the laccase VdtB is responsible for the dimerization step.

The production of semi-viriditoxin **12** is interesting because this shows that the tailoring steps catalyzed by VdtC, VdtF and VdtE most likely to take place prior to dimerization. However, the possibility of these tailoring enzymes acting on dimers cannot be excluded. Furthermore, the fact that the *vdtC*, *vdtF* and *vdtE* deletion mutants produce dimerized compounds shows that VdtB is not strictly specific to semi-viriditoxin in vivo. Our findings concur with the results of a recent study which demonstrated that the VdtB homolog of *A. viridinutans* is able to dimerize the non-native substrate (R)-semi-vioxanthin, using heterologously-produced cell-free extracts from *Aspergillus niger* [15]. However, the heterologously-produced VdtB was shown to lack the appropriate stereospecificity in an in vitro assay [15].

Bioinformatics analysis of the final protein in the cluster, VdtD, shows amino acid sequence similarity to serine hydrolases. This is a functionally diverse group that includes catalytically-active hydrolases and other proteins that have evolved diverse functions independent of catalytic activity. Two such proteins that have lost catalytic function and taken on new roles are neuroligins, which are important components of the post-synaptic membrane [44], and thyroglobulin, a precursor of thyroid hormones [45]. The catalytically active serine hydrolases share a nucleophilic serine residue contained within the conserved G-X-S-X-G motif (where X is any amino acid) at their active site [46]. The non-catalytic variants lack this active site serine [47].

To explore if VdtD is likely to be an active esterase we compared the sequence of VdtD to the top 100 BLAST results in the Swiss-Prot database. These mostly include catalytic enzymes, however there are also some neuroligins and thyroglobulins. A sequence alignment of a subset of these matches showed that, as expected, thyroglobin and neuroligins lacked the serine residue whereas the serine was unmutated in catalytic enzymes. Importantly, VdtD lacks this serine, having instead an aspartate in that position. This suggests that VdtD is probably not an active hydrolase, and as with its mammalian homologues neuroligin and thyroglobulin has taken on new roles.

Intriguingly, disruption of *vdtD* alters the ratio of the two atropisomers of viriditoxin **1** versus **1′** (Figure 6). While the wild-type strain strongly favors the production of the *M* form **1**, the *vdtD* mutant produced favored the P form **1′** as revealed by circular dichroism (Figure 6C). NMR data for **1′** is provided is Table S4 and Figures S9-S14. Despite the growing appreciation of the importance of atropisomeric compounds [19], how organisms selectively synthesize particular atropisomeric forms remains unknown. Given that the laccase VdtB alone has been shown to favor the unexpected *P*-configured stereoisomer in in a heterologous system [15] we suggest VdtD is likely to be required to control the stereoselectivity of the laccase-catalyzed reaction, possibly playing an analogous role to the dirigent proteins of plants [17].

**Fig. 6.**
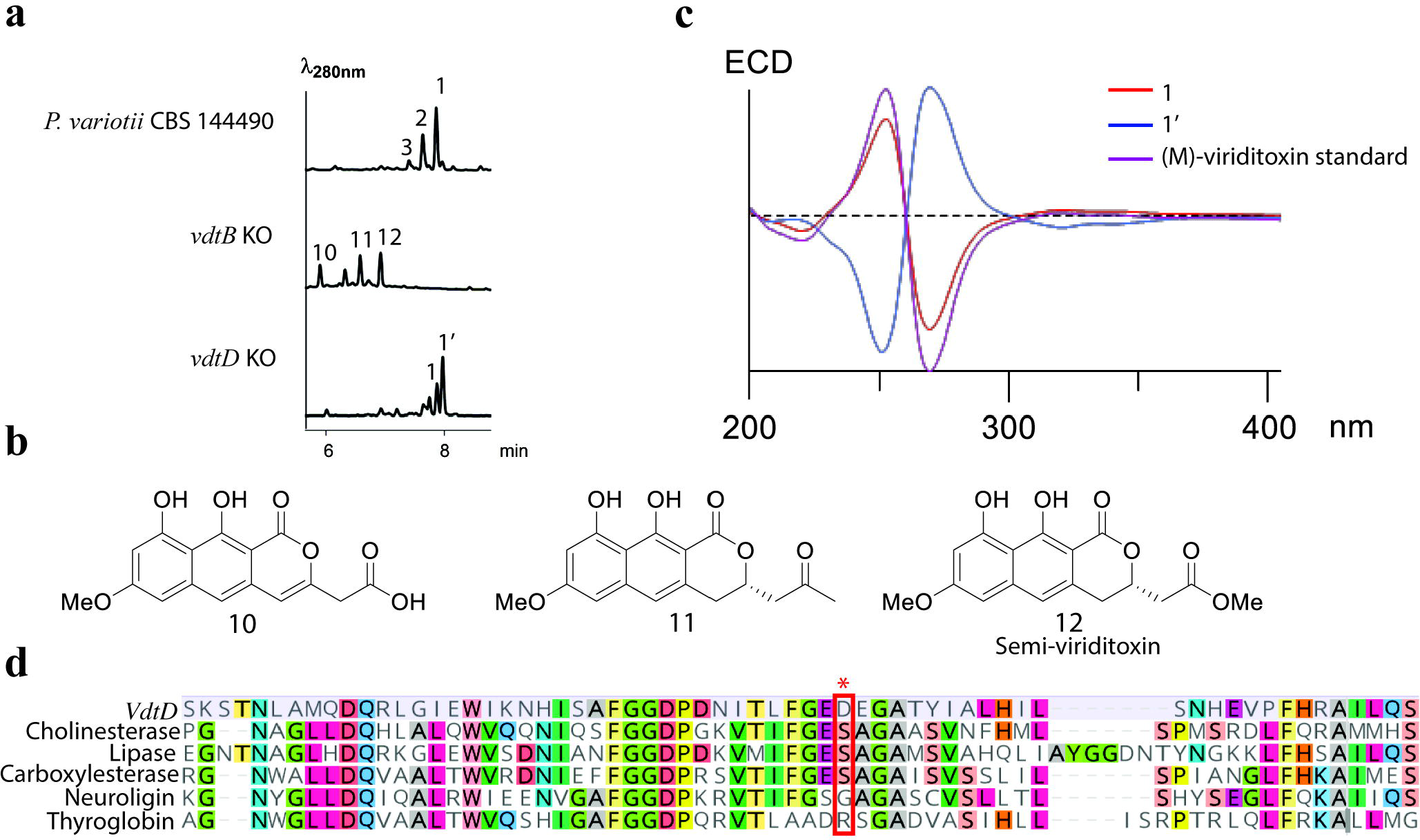
**a**. LC-DAD-MS analysis of the *vdtB* and *vdtD* deletion strains. **b.** The predicted structures of monomers produced by the *vdtB* deletion strain based on mass. **c.** Comparison of the electronic circular dichroism data of 1 vs 1′ showing that the spectra are mirrors of each other, indicative of two atropisomeric forms. **d.** Multiple sequence alignment of VdtD and a lipase from *Geotrichum candidum* (UniProtKB/Swiss-Prot: P22394.2), a choline esterase from *Branchiostoma lanceolatum* (UniProtKB/Swiss-Prot: Q95000.1), a carboxylesterase from *Felis catus* (UniProtKB/Swiss-Prot: Q8I034.1), and a neuroligin (UniProtKB/Swiss-Prot: Q8N0W4.1) and a thyroglobin (UniProtKB/Swiss-Prot: O08710.3) from *Homo sapiens*. Red box encloses the active site serine in the G-X-S-X-G motif of active hydrolases.

The stereochemistry of the metabolites produced by the tailoring-enzyme knockout (Figure 5) provides an initial insight into the substrate requirements of the hypothesized VdtD/VdtB coupling system. The compounds produced by the *vdtF* and *vdtE* knockouts, lacking C3-C4 bond reduction and the methyl ester respectively, are predominantly coupled in the *M* helical conformation as **1**. This suggests that these chemical groups are not integral for interaction with coupling enzymes. However, while not the dominant product as confirmed by ECM (Table S6), some of the corresponding *P* stereoisomer appears to form a shoulder peak in **3** (Figure 3), so we cannot completely exclude the possibility that the methyl ester makes a contribution to the interaction. On the other hand, the metabolites produced by the *vdtC* mutant that lack methylation of the oxygen attached to C7 have been coupled in the unexpected *P* configuration. We hypothesize that interaction of this methyl group with either VdtB or VdtD is likely to be crucial for a controlled stereochemical outcome. Further biochemical studies, which are currently in progress, will seek to establish the mechanism by which VdtD controls the stereospecificity of VdtB.

## Conclusion

The gene cluster responsible for the production of viriditoxin has been identified in the genome of two *Eurotiales* species: *P. variotii* and *A. viridinutans*. Knock-out mutations of all the enzymes required for the production of viriditoxin has allowed us to elucidate the biosynthetic pathway of this helical chiral biaryl compound (Figure 7). Second, we have uncovered an intriguing cellular localization pattern for the PKS enzyme responsible for the first step in the pathway. Third, we have demonstrated that a likely catalytically inactive hydrolase enzyme is required for controlling atropisomeric conformations through an as yet undefined mechanism. These are exciting directions whose resolution are being pursued in complementary biochemical analyses.

**Fig. 7.**
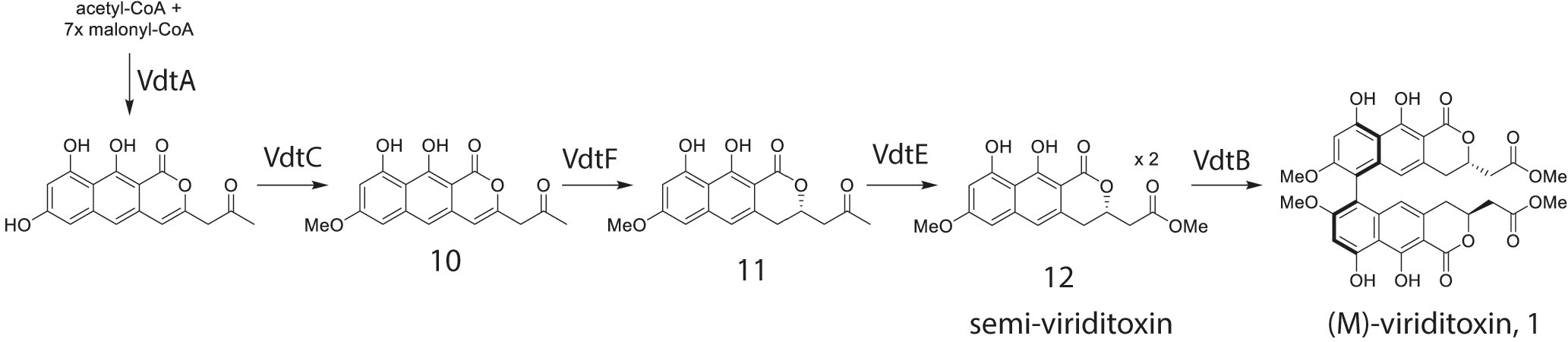
A proposed pathway for the biosynthesis of viriditoxin based on metabolites observed in gene deletion strains. Note that the order in which the enzymes VdtC, VdtF and VdtE act has not been determined.

## Materials and methods

### Strains, DNA extraction, and DNA sequencing of *A. viridinutans*

Two *P. variotii* strains used in this study were CBS101075, which is the type strain of the teleomorph of *P. variotii* (*Byssochlamys spectabilis*), and CBS144490, which we isolated previously [20]. *A. viridinutans* ex-type strain FRR 0576 was used. Genomic DNA was extracted from 4-day old liquid cultures grown in potato dextrose broth (PDB) using an established method [48], followed by treatment with RNAseA. *A. viridinutans* was sequenced on an Illumina HiSeq2500 instrument using 125 bp paired-end reads. The reads were assembled using Velvet [49]. Regions homologous to putative clusters in the *P. variotii* genome were identified by BLASTx searches against the *A. viridinutans* assembly.

### Expression analysis of the cluster via qPCR

RNA was extracted from 3-day old cultures grown in PDB using Trizol reagent (Invitrogen) following the manufacturer’s directions. cDNA was synthesized using AMV reverse transcriptase (New England Biolabs) following the standard protocol provided by the manufacturer. qPCR reactions were conducted using KAPA SYBR FAST qPCR Master Mix. The primer pairs used for each gene are given in Table S12.

### Generation of gene knockout strains

Gene knock outs were generated by homologous recombination for each gene in the cluster, with the exception of *vdtX*. These constructs were generated by first assembling two fragments corresponding to the upstream and downstream gene regions, into plasmid pPZP-201BK digested with EcoRI and HindIII, using Gibson Assembly with the addition of a restriction site between the two fragments. The HYG cassette on plasmid pMAI6 was amplified by PCR and introduced into this restriction site using a second Gibson reaction. Primers used to amplify the HYG cassette with appropriate homology to introduce it into each construct are given in Table S12.

The constructs were introduced into *P. variotii* strain CBS 144490 or CBS101075 via *Agrobacterium tumefaciens* mediated transformation (*At*MT) as described previously [20]. Successful gene replacement events were identified via PCR using primers given in Table S12.

### Disruption of *vdtX* via repeat induced point mutation

A different approach was taken to disrupt *vdtX* because no gene replacements were identified from the homologous recombination approach. The process of repeat induced point (RIP) mutation during a sexual cross was employed. A partial *vdtX* gene fragment was amplified using primers RIPF1 and RIPR2 and cloned into pPZPHyg*Hin*dX [50] linearized with XbaI-EcoRV using Gibson assembly. The construct was then transformed into strain CBS 144490. The resultant transformants were crossed to strain CBS 101075 as described previously [51]. The targeted region in *vdtX* of the resultant transformants was amplified and sequenced with primers RIPseqF and RIPseqR to identify those that had been heavily mutated by RIP.

### Metabolic profile analysis, compound isolation and characterization

Czapek Yeast Extract Agar (CYA, 30 g Sucrose, 5g Yeast Extract, 1g K_2_HPO_4_, 0.3 g NaNO_3_, 0.05 g KCl, 0.05 g MgSO_4_, 0.01 g ZnSO_4_, 0.01 g FeSO_4_, 0.5 mg CuSO_4_, 15 g Agar in 1 L) was used for culturing of *P. variotii* strains and *A. viridinutans*, and accumulation of secondary metabolites. Agar cultures were chopped into small pieces and extracted with methanol under sonication condition. Crude extracts were filtered, dried and redissolved in methanol for LC-DAD-MS analysis.

LC-DAD-MS analysis was performed with Agilent 1260 liquid chromatography (LC) system coupled to a diode array detector (DAD) and an Agilent 6130 quadrupole mass spectrum (MS) with an ESI source. For analytical purpose, a Kinetex C18 column (2.6 μm, 2.1 mm i.d. × 100 mm; Phenomenex) was used. The mobile phase gradient of eluent B (acetonitrile with 0.1% formic acid) started at 5% and gradually increased to 95% over 10 mins at a flow rate of 0.75 ml/min.

For compound isolation, *P. variotii* CYA culture of either wild-type strain or specific mutant was extracted twice with ethyl acetate/methanol (90:10). Crude extracts were dried *in vacuo* and then fractionated on a Reveleris flash chromatography system (Grace) using a dichloromethane/methanol gradient on a Reveleris HP silica flash cartridge. Fractions containing the target compound were combined for further purification using a semi-prep HPLC with a C18 column (Agilent, 5 μm, 21.2 × 150 mm).

For structural characterization, nuclear magnetic resonance (NMR) spectra were collected for purified metabolites on a Bruker Avance IIIHD 500MHz/600MHz NMR spectrometer. Chloroform-*d*, DMSO-*d*_6_, and acetonitrile-*d*_3_ were used as solvents.

Electronic circular dichroism (ECD) spectra were recorded on a JASCO J-810 spectropolarimeter, with acetonitrile as solvent. The axial chirality of dimeric compounds isolated was determined by comparing with both the ECD spectrum of (*M*)-viriditoxin standard purchased from Sapphire Bioscience (Redfern, Australia) and published data [13].

### Generation and examination of fluorescently-tagged proteins

The open reading frame of GFP was tagged immediately downstream of *vdtG* and *vdtA* via homologous recombination.

In the case of *vdtG*, a fragment corresponding to the end of the gene (omitting the stop codon) was amplified using primers 24GFP5F and 24GFP5R and GFP was amplified using primers 24GFPGFPF and 24GFPGFPR and cloned into plasmid pPZP-201BK linearized with EcoRI and HindIII using Gibson assembly. The HYG construct and 3′ flank of the gene was amplified with primers pairs 24GFPHYGF-24GFPHYGR and 24GFP3F-24GFP3R off plasmids pMAI6 and PLAU17, respectively [30] and cloned into the EcoRI site of the previously generated plasmid to produce the final construct. This construct was transformed into CBS 144490 using *At*MT [20].

In the case of *vdtA*, a fragment corresponding to the end of the gene (omitting the stop codon) was amplified using primers 69GFP5F and 69GFP5R and the 3′ flank of the gene was amplified with primers 69GFP3F and 69GFP3R and cloned into plasmid pPZP-201BK linearized with EcoRI and HindIII using Gibson assembly. The combined GFP-HYG section of the construct previously generated to tag VdtG was amplified using 69GFPHYGF and 69GFPHYGR and cloned into the BamHI site of the previous construct. This construct was introduced into CBS 144490 by *At*MT as described previously [20].

Constructs were generated for the purpose of resolving the subcellular localization of proteins within the cell, namely Erg11-mCherry to tag the endoplasmic reticulum and mCherry-SKL to tag the peroxisomes. Erg11 was amplified using primers Erg11F and Erg11R and mCherry was amplified using primers McherryErg11F and McherryErg11R off genomic DNA and plasmid mCherry-dspA [20], respectively, and cloned into the BglII site of plasmid PLAU53 which has been designed to express proteins under the expression of the actin promoter from *Leptosphaeria maculans* and confers resistance to G418 [30]. The mCherry isoform with the addition of sequence coding for the three amino acid sequence SKL was amplified using primers AP57 and AP58, and similarly cloned into PLAU53. The mCherry-SKL construct was transformed into the VdtA-GFP strain and the Erg11-mCherry construct was transformed into the VdtG-GFP strain.

Isolates were cultured PDB for 3 days, and then examined using a Leica DM6000 fluorescence microscope.

## Authors’ contributions

ASU, JH, Y-HC and AI designed the experiments. ASU performed the sequencing, sequence analysis, and genetic manipulations of *P. variotii*. JH performed the chemical purifications and metabolite characterizations. ASU, JH, Y-HC and AI analyzed and interpreted the data and results. ASU was the major contributor in writing the manuscript. JH, Y-HC and AI provided critical feedback and helped in writing the manuscript. All authors read and approved the final manuscript.

## Acknowledgements

We thank Mark Wilson (CSIRO, Australia) for providing the *A. viridinutans* strain and Allison van de Meene (University of Melbourne) for assistance with microscopy.

## Availability of data and materials

The *A. viridinutans* genome sequencing raw reads and assembly are available from GenBank. NMR data are presented in the supplemental information. Strains are available from AI upon request.

## Consent for publication

Not applicable.

## Ethics approval and consent to participate

Not applicable.

