## Supplemental data for "The fungal gene cluster for biosynthesis of the antibacterial agent viriditoxin"

**Table S1. Genome sequencing statistics for the assembly of *A. viridinutans* strain FRR 0576.**

| | |
|---|---|
| Number of contigs | 285 |
| Median length (bp) | 5,707 |
| Mean length (bp) | 104,615 |
| Max length (bp) | 1,359,167 |
| N50 length (bp) | 425,947 |
| Number of contigs >N50 | 21 |
| Length Sum (bp) | 29,815,307 |

**Table *S*2. Protein IDs of the Vdt cluster genes in *P. variotii* strains CBS 101075 and CBS 144490.**

| Protein | Protein ID<br>CBS 101075 | Protein ID<br>CBS 144490 | Putative function |
|---|---|---|---|
| VdtA | 480069 | 260870 | Polyketide synthase |
| VdtB | 480050 | 260889 | Laccase |
| VdtC | 488617 | 127223 | *O*-methyltranstrase |
| VdtD | 510289 | 190631 | Hydrolase-like |
| VdtE | 480056 | 190779 | BVMO |
| VdtF | 480057 | 260883 | Reductase |
| VdtG | 488624 | 275282 | MFS transporter |
| VdtR | 105452 | 288289 | Transcription factor |
| VdtX | 515060 | 190767 | PKS-like |

#### Table S3. Structural information of 1 (chloroform-*d*).

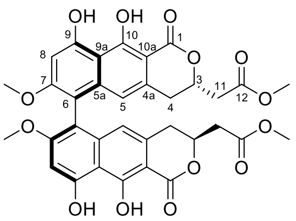

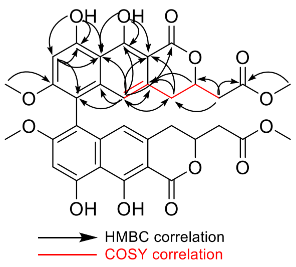


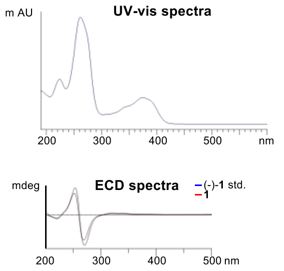


| Carbon No. | $^{13}C$ NMR | $^{1}H$ NMR (ppm, multi, *J*) | gCOSY | HMBC |
|---|---|---|---|---|
| 1 | 171.0 | - | - | - |
| 2 | - | - | - | - |
| 3 | 75.9 | 4.96 (1H, m) | 4, 11 | 4a |
| 4 | 33.0 | 2.81 (2H, m) | 3, 5 | 4a, 5, 10a |
| 4a | 132.1 | - | | - |
| 5 | 114.2 | 6.23 (1H, s) | 4 | 1, 3, 4a, 5a, 6, 9a, 10, 10a |
| 5a | 139.3 | - | - | - |
| 6 | 110.1 | - | - | - |
| 7 | 161.2 | - | - | - |
| 7-OMe | 56.3 | 3.77 (3H, s) | - | 7 |
| 8 | 98.4 | 6.79 (1H, s) | - | 6, 9, 9a |
| 9 | 159.3 | - | - | - |
| 9-OH | - | 9.69 (1H, s) | - | 8, 9, 9a |
| 9a | 108.1 | - | - | - |
| 10 | 163.5 | - | - | - |
| 10-OH | - | 13.75 (1H, s) | - | 9a, 10, 10a |
| 10a | 98.8 | - | - | - |
| 11 | 39.5 | 2.65 (1H, dd, *J*=7.74, 19.5)<br>2.88 (1H, dd, *J*= 8.16, 19.4) | 4 | 3, 4, 12 |
| 12 | 169.8 | - | - | - |
| 12-OMe | 52.2 | 3.70 (3H, s) | - | 12 |

#### Table S4. Structural information of 1′ (chloroform-*d*).

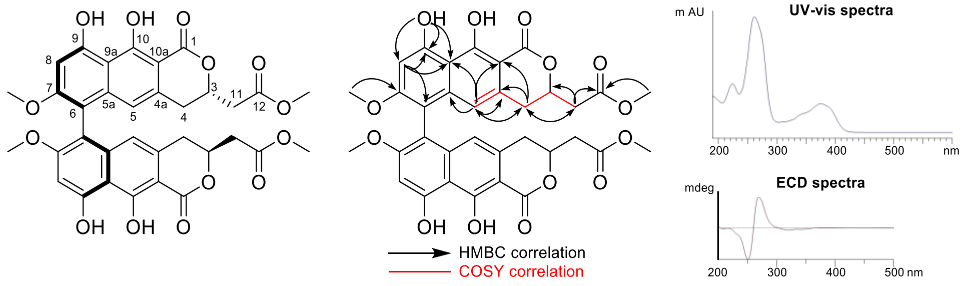


| Carbon No. | $^{13}$C NMR | $^{1}$H NMR (ppm, multi, *J*) | gCOSY | HMBC |
|---|---|---|---|---|
| 1 | 171.1 | - | - | - |
| 2 | - | - | - | - |
| 3 | 75.9 | 4.96 (1H, m) | 4, 11 | - |
| 4 | 32.9 | 2.87 (2H, m) | 3, 5 | 4a, 5, 10a, 11 |
| 4a | 132.1 | - | | - |
| 5 | 114.2 | 6.24 (1H, s) | 4 | 4, 4a, 5a, 9a, 10a |
| 5a | 139.4 | - | - | - |
| 6 | 110.2 | - | - | - |
| 7 | 161.2 | - | - | - |
| 7-OMe | 56.3 | 3.75 (3H, s) | - | 7 |
| 8 | 98.3 | 6.79 (1H, s) | - | 6, 9, 9a |
| 9 | 159.3 | - | - | - |
| 9-OH | - | 9.77 (1H, s) | - | 8, 9, 9a |
| 9a | 108.2 | - | - | - |
| 10 | 163.6 | - | - | - |
| 10-OH | - | 13.77 (1H, s) | - | - |
| 10a | 98.9 | - | - | - |
| 11 | 39.5 | 2.68 (1H, dd, *J*=6.55, 16.3)<br>2.92 (1H, dd, *J*= 6.70, 16.3) | 4 | 3, 4, 12 |
| 12 | 169.9 | - | - | - |
| 12-OMe | 52.3 | 3.70 (3H, s) | - | 12 |

### Table S5. Structural information of 2 (acetonitrile-$d_3$).

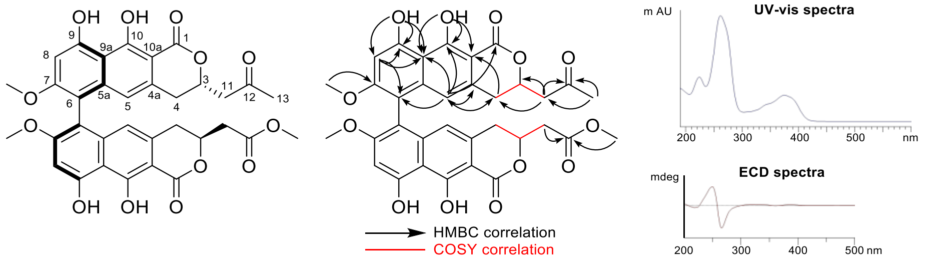


| Carbon No. | $^{13}$C NMR | $^{1}$H NMR (ppm, multi, $J$) | gCOSY | HMBC |
|---|---|---|---|---|
| 1 | 172.2 | - | - | - |
| 1′ | 172.0 | - | - | |
| 2 (2′) | - | - | - | - |
| 3 | 76.8 | 4.96 (2H, m) | 4, 11 | - |
| 3′ | 77.3 | 4.96 (2H, m) | 4′, 11′ | - |
| 4 | 32.8 | 2.68[a] | 3 | 5, 10a |
| 4′ | 33.0 | 2.68[a] | 3′ | 5′, 10a′ |
| 4a | 134.2 | - | - | - |
| 4a′ | 134.0 | - | - | - |
| 5 | 114.8 | 6.28 (2H, s) | - | 1, 4, 4a, 6, 9a, 10 |
| 5′ | 114.7 | 6.28 (2H, s) | - | 1′, 4′, 4a′, 6′, 9a′, 10′ |
| 5a (5a′) | 140.1 | - | - | - |
| 6 (6′) | 110.9 | - | - | - |
| 7 (7′) | 161.8 | - | - | - |
| 7 (7′)-OMe | 56.8 | 3.71 (6H, s) | - | 7 (7′) |
| 8 (8′) | 99.0 | 6.83 (1H, s) | - | 6 (6′), 9 (9′), 9a (9a′) |
| 9 (9′) | 159.9 | - | - | - |
| 9 (9′)-OH | - | 9.68 (2H, s) | - | 8 (8′), 9 (9′), 9a (9a′) |
| 9a (9a′) | 108.5 | - | - | - |
| 10 | 163.7 | - | - | - |
| 10′ | 163.8 | - | - | - |
| 10 (10′)-OH | - | 13.8 (1H, s) | - | 9a (9a′), 10 (10′), 10a (10a′) |
| 10a | 100.0 | - | - | - |
| 10a′ | 99.9 | - | - | - |
| 11 | 48.3 | 2.74[a] 2.96 (1H, dd, $J$=7.65, 17.4) | 3 | 3, 4, 12 |
| 11′ | 39.9 | 2.70[a] | 3′ | 3′, 4′, 12′ |
| 12 | 205.7 | - | - | - |

| | | | | |
|---|---|---|---|---|
| 12′ | 170.9 | - | - | - |
| 13 | 30.6 | 2.12 (3H, s) | - | 11, 12 |
| 12′-OMe | 52.4 | 3.65 (3H, s) | - | 12′ |

[a] unable to integrate or calculate *J* due to signal overlay

#### Table S6. Structural information of 3 (acetonitrile-$d_3$).

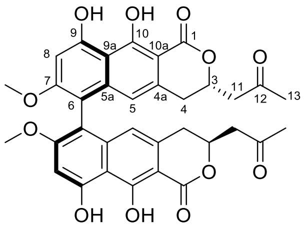

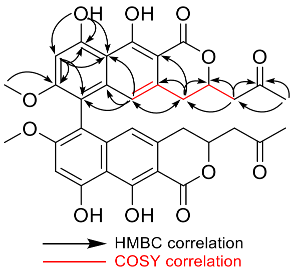


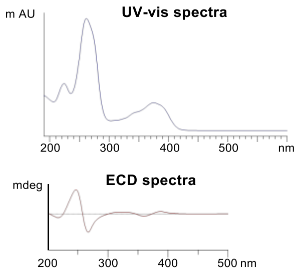


| Carbon No. | $^{13}$C NMR | $^{1}$H NMR (ppm, multi, *J*) | gCOSY | HMBC |
|---|---|---|---|---|
| 1 | 172.2 | - | - | - |
| 2 | - | - | - | - |
| 3 | 76.8 | 4.98 (1H, m) | 4, 11 | - |
| 4 | 33.0 | 2.79 (2H, m) | 3, 5 | 3, 4a, 5, 10a |
| 4a | 134.4 | - | | - |
| 5 | 114.7 | 6.27 (1H, s) | 4 | 4, 5a, 6, 9a |
| 5a | 140.2 | - | - | - |
| 6 | 110.9 | - | - | - |
| 7 | 161.8 | - | - | - |
| 7-OMe | 56.8 | 3.71 (3H, s) | - | 7 |
| 8 | 99.0 | 6.84 (1H, s) | - | 6, 7, 9, 9a |
| 9 | 159.9 | - | - | - |
| 9-OH | - | 9.73 (1H, s) | - | 8, 9, 9a |
| 9a | 108.6 | - | - | - |
| 10 | 163.8 | - | - | - |
| 10-OH | - | 13.84 (1H, s) | - | - |
| 10a | 100.1 | - | - | - |
| 11 | 48.3 | 2.76 (1H, dd, *J*=7.92, 17.5)<br>2.96 (1H, dd, *J*= 7.74, 17.5) | 4 | 3, 4, 12 |
| 12 | 205.7 | - | - | - |
| 13 | 30.6 | 2.11 (3H, s) | - | 11, 12 |

### Table S7. Structural information of 4 (acetonitrile-$d_3$).

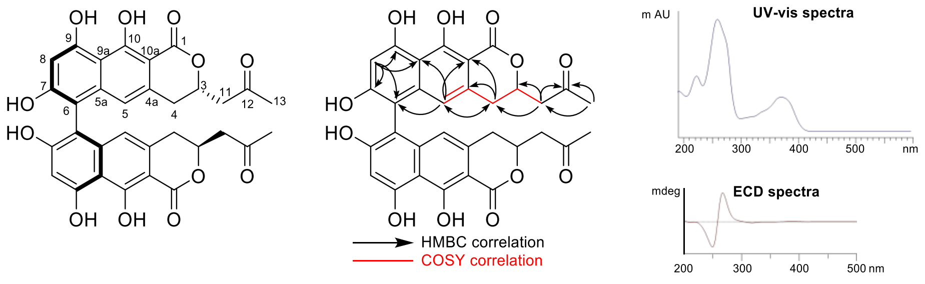


| Carbon No. | $^{13}$C NMR | $^{1}$H NMR (ppm, multi, *J*) | gCOSY | HMBC |
|---|---|---|---|---|
| 1 | 172.2 | - | - | - |
| 2 | - | - | - | - |
| 3 | 76.8 | 4.92 (1H, m) | 4, 11 | - |
| 4 | 33.1 | 2.79[a] | 3, 5 | 4a, 5, 10a |
| 4a | 135.0 | - | | - |
| 5 | 114.4 | 6.35 (1H, s) | 4 | 4, 9a, 10a |
| 5a | 141.5 | - | - | - |
| 6 | 109.2 | - | - | - |
| 7 | 160.5 | - | - | - |
| 7-OH | - | - | - | - |
| 8 | 102.7 | 6.60 (1H, s) | - | 6, 7, 9, 9a |
| 9 | 160.0 | - | - | - |
| 9-OH | - | - | - | - |
| 9a | 107.0 | - | - | - |
| 10 | 163.6 | - | - | - |
| 10-OH | - | - | - | - |
| 10a | 100.3 | - | - | - |
| 11 | 48.3 | 2.78[a]<br>2.95 (1H, dd, *J*=7.45, 17.45) | 3 | 3, 4, 12 |
| 12 | 205.9 | - | - | - |
| 13 | 30.6 | 2.11 (3H, s) | - | 11, 12 |

[a] unable to integrate or calculate *J* due to signal overlay

**Table S8. Structural information of 5 (acetonitrile-$d_3$).**

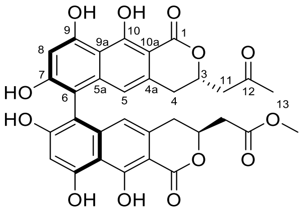


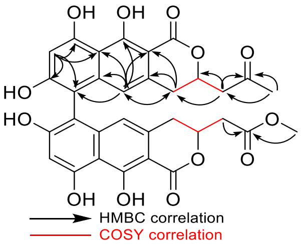


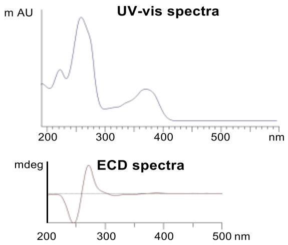


| Carbon No. | $^{13}$C NMR | $^{1}$H NMR (ppm, multi, *J*) | gCOSY | HMBC |
|---|---|---|---|---|
| 1 | .172.2 | - | - | - |
| 1′ | 172.0 | - | - | |
| 2 (2′) | - | - | - | - |
| 3 | 76.8 | 4.89 (1H, m) | 4, 11 | - |
| 3′ | 77.3 | 4.89 (1H, m) | 4′, 11′ | - |
| 4 | 33.0 | 2.81[a] | 3 | 4a, 5, 10a |
| 4′ | 32.8 | 2.81[a] | 3′ | 4a′, 5′, 10a′ |
| 4a | 135.0 | - | - | - |
| 4a′ | 134.7 | - | - | - |
| 5 | 114.5 | 6.36 (1H, s) | - | 4, 5a, 6, 9a, 10, 10a |
| 5′ | 114.4 | 6.36 (1H, s) | - | 4′, 5a′,6′, 9a′, 10′, 10a′ |
| 5a (5a′) | 141.5 | - | - | - |
| 6 (6′) | 106.9 | - | - | - |
| 7 (7′) | 160.5 | - | - | - |
| 7 (7′)-OH | - | - | - | - |
| 8 (8′) | 102.7 | 6.60 (1H, s) | - | 6 (6′), 7 (7′), 9 (9′), 9a (9a′) |
| 9 (9′) | 160.0 | - | - | - |
| 9 (9′)-OH | - | - | - | - |
| 9a (9a′) | 109.2 | - | - | - |
| 10 | 163.5 | - | - | - |
| 10′ | 163.6 | - | - | - |
| 10 (10′)-OH | - | - | - | - |
| 10a | 100.0 | - | - | - |
| 10a′ | 99.9 | - | - | - |
| 11 | 48.3 | 2.74[a] 2.96 (1H, dd, *J*=7.44, 17.52) | 3 | 3, 4, 12 |
| 11′ | 39.9 | 2.70[a] | 3′ | 3′, 4′, 12′ |
| 12 | 205.7 | - | - | - |

| | | | | |
|---|---|---|---|---|
| 12′ | 170.0 | - | - | - |
| 13 | 30.6 | 2.10 (1.5H, s) | - | 11, 12 |
| 12′-OMe | 52.5 | 3.62 (1.5H, s) | - | 12′ |

[a] unable to integrate or calculate *J* due to signal overlay

#### Table S9. Structural information of 7 (chloroform-*d*).

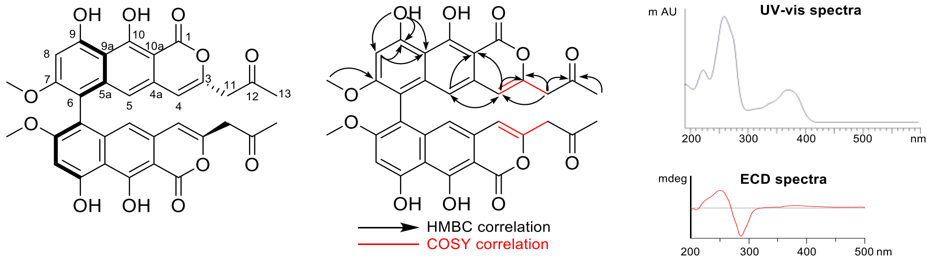


| Carbon No. | $^{13}$C NMR | $^{1}$H NMR (ppm, multi, *J*) | gCOSY | HMBC |
|---|---|---|---|---|
| 1 | 168.0[a] | - | - | - |
| 2 | - | - | - | - |
| 3 | 148.4 | - | - | - |
| 4 | 108.3 | 6.10 (1H, s)] | 11 | 3, 5, 10a |
| 4a | -[b] | - | | - |
| 5 | 111.7 | 6.36 (1H, s) | - | 4, 10a |
| 5a | 140.2[a] | - | - | - |
| 6 | 109.9[a] | - | - | - |
| 7 | 161.3 | - | - | - |
| 7-OMe | 56.4 | 3.77 (3H, s) | - | 7 |
| 8 | 98.4 | 6.84 (1H, s) | - | 9, 9a |
| 9 | 159.3 | - | - | - |
| 9-OH | - | 9.75(1H, s) | - | 8, 9, 9a |
| 9a | -[b] | - | - | - |
| 10 | 163.0[a] | - | - | - |
| 10-OH | - | 13.6 (1H, s) | - | - |
| 10a | 97.5 | - | - | - |
| 11 | 47.6 | 3.47 (2H, s) | 4 | 3, 4, 12 |
| 12 | 202.2 | - | - | - |
| 13 | 30.1 | 2.23 (3H, s) | - | 12 |

[a] Lacking HMBC correlations. Chemical shifts assigned based on other compounds isolated.
[b] $^{13}$C signals not detected.

#### Table S10. Structural information of 8 (chloroform-*d*).

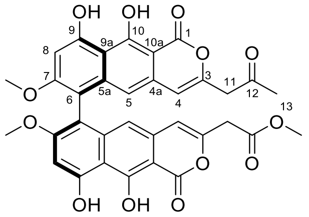

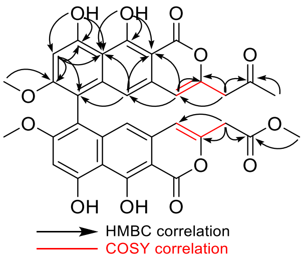


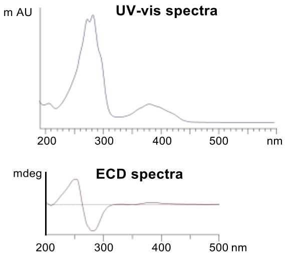


| Carbon No. | $^{13}$C NMR | $^{1}$H NMR (ppm, multi, *J*) | gCOSY | HMBC |
|---|---|---|---|---|
| 1 | .168.0 | - | - | - |
| 1′ | 167.9 | - | - | |
| 2 (2′) | - | - | - | - |
| 3 | 148.4 | - | - | - |
| 3′ | 147.9 | - | - | - |
| 4 | 108.3 | 6.10 (1H, s) | 11 | 3, 5, 10a |
| 4′ | 108.0 | 6.15 (1H, s) | 11′ | 3′, 5′, 10a′ |
| 4a | 130.2 | - | - | - |
| 4a′ | 130.5 | - | - | - |
| 5 | 111.7 | 6.36 (1 H, s) | - | 6, 9a, 10a |
| 5′ | 111.8 | 6.37 (1 H, s) | - | 6′, 9a′, 10a′ |
| 5a (5a′) | 140.2 | - | - | - |
| 6 (6′) | 109.9 | - | - | - |
| 7 (7′) | 161.3 | - | - | - |
| 7 (7′)-OMe | 56.4 | 3.77 (6H, s) | - | 7 (7′) |
| 8 (8′) | 98.4 | 6.84 (2H, s) | - | 6 (6′), 7 (7′), 9 (9′), 9a (9a′) |
| 9 (9′) | 159.3 | - | - | - |
| 9 (9′)-OH | - | 9.75 (2H, s) | - | 8 (8′), 9 (9′), 9a (9a′) |
| 9a (9a′) | 108.2 | - | - | - |
| 10 (10′) | 163.0 | - | - | - |
| 10 (10′)-OH | - | 13.6 (2H, s) | - | 9a (9a′), 10 (10a′), 9a (9a′) |
| 10a | 97.5 | - | - | - |
| 10a′ | 97.6 | - | - | - |
| 11 | 47.6 | 3.47 (2H, s) | 4 | 3, 4, 12 |
| 11′ | 38.8 | 3.43 (2H, s) | 4′ | 3′, 4′, 12′ |
| 12 | 202.2 | - | - | - |
| 12′ | 168.7 | - | - | - |

| | | | | |
|---|---|---|---|---|
| 13 | 30.1 | 2.23 (3H, s) | - | 12 |
| 12′-OMe | 52.7 | 3.71 (3H, s) | - | 12′ |

#### Table S11. Structural information of 9 (chloroform-*d*).

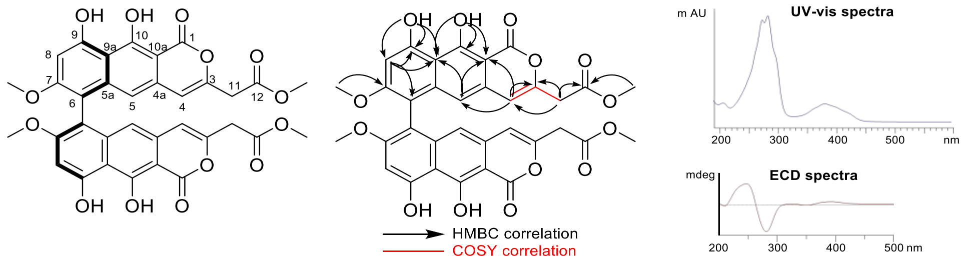


| Carbon No. | 13C NMR | 1H NMR (ppm, multi, *J*) | gCOSY | HMBC |
|---|---|---|---|---|
| 1 | .167.9[a] | - | - | - |
| 2 | - | - | - | - |
| 3 | 147.9 | - | - | - |
| 4 | 108.0 | 6.15 (1H, s) | 11 | 3, 5, 10a |
| 4a | 130.8[a] | - | | - |
| 5 | 111.8 | 6.37 (1H, s) | - | 9a, 10a |
| 5a | 140.2[a] | - | - | - |
| 6 | 110.0[a] | - | - | - |
| 7 | 161.2 | - | - | - |
| 7-OMe | 56.4 | 3.77 (3H, s) | - | 7 |
| 8 | 98.5 | 6.84 (1H, s) | - | 6, 9, 9a |
| 9 | 159.3 | - | - | - |
| 9-OH | - | 9.69 | - | 8, 9, 9a |
| 9a | 108.1 | - | - | - |
| 10 | 163.0 | - | - | - |
| 10-OH | - | 13.75 | - | 9a, 10, 10a |
| 10a | 97.6 | - | - | - |
| 11 | 38.8 | 3.43 (2H, s) | 4 | 3, 4, 12 |
| 12 | 168.7 | - | - | - |
| 12-OMe | 52.7 | 3.71(3H, s) | - | 12 |

[a] 13C signals have no HMBC correlations. The chemical shifts were assigned based on data of other compounds.

#### Table S12. Oligonucleotide primers designed in this study

| Name | Sequence (5′ to 3′) | Notes |
|---|---|---|
| *vdtB5'F* | AACAGCTATGACATGATTACGAATCGACGTTGTTTCTTGG | Primers used to amplify 5′ flank of *vdtB* |
| *vdtB5'R* | GTGGGCAGAATTCGCTCCATGGCAGCAATGTCC | |
| *vdtC5'F* | AACAGCTATGACATGATTACGTGCTTCTTTATACTGGGTCG | Primers used to amplify 5′ flank of *vdtC* |
| *vdtC5'R* | TTGAAGAGAATTCGAAGCACTGATTGCCTGTGC | |
| *vdtD5'F* | AACAGCTATGACATGATTACGCCTTCTTCAAGTATCTTTCC | Primers used to amplify 5′ flank of *vdtD* |
| *vdtD5'R* | ACATACGGAATTCTGGTAACGGAAGGAAAAGACC | |
| *vdtE5'F* | AACAGCTATGACATGATTACGGGTTTGTAGGGATCGAATGC | Primers used to amplify 5′ flank of *vdtE* |
| *vdtE5'R* | CCGAAGGGAATTCCAACTCTGAAATACATCTCC | |
| *vdtF5'F* | AACAGCTATGACATGATTACGCGTCAATCGAGTTGAAATGG | Primers used to amplify 5′ flank of *vdtF* |
| *vdtF5'R* | CAGTCGAGAATTCGTCTGTTCTGTTGTTGGTGC | |
| *vdtG5'F* | AACAGCTATGACATGATTACGGGTCTGTTCTGTTGTTGGTGC | Primers used to amplify 5′ flank of *vdtG* |
| *vdtG5'R* | CCTCAGGGAATTCTGATCGAGCGCCAACCTACC | |
| *vdtA5'F* | AACAGCTATGACATGATTACGTTCTTATATACTTATAATTCG | Primers used to amplify 5′ flank of *vdtA* |
| *vdtA5'R* | GAGAACGGGATCCATAAACGACGCAAGCTGG | |
| *vdtR5'F* | AACAGCTATGACATGATTACGGCGTGCGTAGTAGCTGCTGC | Primers used to amplify 5′ flank of *vdtR* |
| *vdtR5'R* | AGATTGGAATTCTTGAGATAGGGTTTGACTGG | |
| *vdtB3'F* | ATGGAGCGAATTCTGCCCACGGAGACGATATGC | Primers used to amplify 3′ flank of *vdtB* |
| *vdtB3'R* | GTAAAACGACGGCCAGTGCCAGGGCACACAAAGAATCTAGG | |
| *vdtC3'F* | GTGCTTCGAATTCTCTTCAAGTATCTTTCCAGC | Primers used to amplify 3′ flank of *vdtC* |
| *vdtC3'R* | GTAAAACGACGGCCAGTGCCAGGTAACGGAAGGAAAAGACC | |
| *vdtD3'F* | GTTACCAGAATTCCGTATGTCTTTAATGAGTTGG | Primers used to amplify 3′ flank of *vdtD* |
| *vdtD3'R* | GTAAAACGACGGCCAGTGCCAATAATAATGTATCCGCTAGG | |
| *vdtE3'F* | AGAGTTGGAATTCCCTTCGGAGGAAAACGTACC | Primers used to amplify 3′ flank of *vdtE* |
| *vdtE3'R* | GTAAAACGACGGCCAGTGCCAATGCTAATATTCTGCCCTGC | |
| *vdtF3'F* | AACAGACGAATTCTCGACTGCCTGCTTCACTGG | Primers used to amplify 3′ flank of *vdtF* |
| *vdtF3'R* | GTAAAACGACGGCCAGTGCCAGGCTTCGACACAAACACTGG | |
| *vdtG3'F* | TCGATCAGAATTCCCTGAGGGGAGGAAACAAGG | Primers used to amplify 3′ flank of *vdtG* |
| *vdtG3'R* | GTAAAACGACGGCCAGTGCCAGCCCAGCCCATTACTGCAGC | |
| *vdtA3'F* | CGTTTATGGATCCCGTTCTCATTCGGTACAGC | Primers used to amplify 3′ flank of *vdtA* |
| *vdtA3'R* | GTAAAACGACGGCCAGTGCCAAAAAGGGGTGGATAATAA | |

| | | |
|---|---|---|
| | GC | |
| *vdtR3'F* | TATCTCAAGAATTCCAATCTGCCCTCGAGTACG | Primers used to amplify 3′ flank of *vdtR* |
| *vdtR3'R* | GTAAAACGACGGCCAGTGCCAGTGCATTTCTTATACTCAAGG | |
| *vdtB*HYGF | GGACATTGCTGCCATGGAGCGCTGGGATTGCCCCTCGATGC | Primers used to amplify HYG cassette for introduction into the *vdtB* construct |
| *vdtB*HYGR | GCATATCGTCTCCGTGGGCAGCCTACTGAACGTTATGAC | |
| *vdtC*HYGF | CACAGGCAATCAGTGCTTCGCTGGGATTGCCCCTCGATGC | Primers used to amplify HYG cassette for introduction into the *vdtC* construct |
| *vdtC*HYGR | CTGGAAAGATACTTGAAGAGCCTACTGAACGTTATGAC | |
| *vdtD*HYGF | TCTTTTCCTTCCGTTACCAGCTGGGATTGCCCCTCGATGC | Primers used to amplify HYG cassette for introduction into the *vdtD* construct |
| *vdtD*HYGR | AACTCATTAAAGACATACGGCCTACTGAACGTTATGAC | |
| *vdtE*HYGF | GAGATGTATTTCAGAGTTGGCTGGGATTGCCCCTCGATGC | Primers used to amplify HYG cassette for introduction into the *vdtE* construct |
| *vdtE*HYGR | GTACGTTTTCCTCCGAAGGGCCTACTGAACGTTATGAC | |
| *vdtF*HYGF | CACCAACAACAGAACAGACGCTGGGATTGCCCCTCGATGC | Primers used to amplify HYG cassette for introduction into the *vdtF* construct |
| *vdtF*HYGR | CAGTGAAGCAGGCAGTCGAGCCTACTGAACGTTATGAC | |
| *vdtG*HYGF | GTAGGTTGGCGCTCGATCAGCTGGGATTGCCCCTCGATGC | Primers used to amplify HYG cassette for introduction into the *vdtG* construct |
| *vdtG*HYGR | CTTGTTTCCTCCCCTCAGGGCCTACTGAACGTTATGAC | |
| *vdtA*HYGF | CCAGCTTGCGTCGTTTATGCTGGGATTGCCCCTCGATGC | Primers used to amplify HYG cassette for introduction into the *vdtA* construct |
| *vdtA*HYGR | GCTGTACCGAATGAGAACGGCCTACTGAACGTTATGAC | |
| *vdtR*HYGF | CCAGTCAAACCCTATCTCAAGCTGGGATTGCCCCTCGATGC | Primers used to amplify HYG cassette for introduction into the *vdtR* construct |
| *vdtR*HYGR | CGTACTCGAGGGCAGATTGCCTACTGAACGTTATGAC | |
| | **qPCR primers** | |
| **Name** | **Sequence (5′ to 3′)** | **Notes** |
| *vdtBqPCRF* | CTATCACAGACAAAGGGTGC | qPCR primers for *vdtB* |
| *vdtBqPCRR* | AGGGAACACAGCCTTCCTGG | |
| *vdtCqPCRF* | GCGATCGTGGTTGATGTTGG | qPCR primers for *vdtC* |
| *vdtCqPCRR* | TCACCATGGGCAGGTCTTGC | |
| *vdtDqPCRF* | ACGGGTTCCGCGCATTTATGC | qPCR primers for *vdtD* |

| | | |
|---|---|---|
| *vdtDqPCRR* | AGGATAGATATATTGGCTGG | |
| *vdtEqPCRF* | GAGCCGTTGAATTGGTTTGG | qPCR primers for *vdtE* |
| *vdtEqPCRR* | AAGACCGAAGAGAATATTCC | |
| *vdtFqPCRF* | CATCTACGGTGCGAGTAAGG | qPCR primers for *vdtF* |
| *vdtFqPCRR* | ATGTCGGTGTTGAAATATCC | |
| *vdtGqPCRF* | AGGATGCGGAATGAGCATGC | qPCR primers for *vdtG* |
| *vdtGqPCRR* | AAGAAGATGAGCGAGATACC | |
| *vdtX1qPCRF* | AGAGAGTAAACGTCTGTTGG | qPCR primers for *vdtX* |
| *vdtXqPCRR* | GCATTGTCGGATATCCGTCG | |
| *vdtAqPCRF* | AGCGACGGACAGATTTCTCG | qPCR primers for *vdtA* |
| *vdtAqPCRR* | TATAGGGTTCCTCCTGCAGC | |
| *β-tubulinqPCRF* | GGGCGAGGAGGAGTACAACG | qPCR primers for β-tubulin |
| *β-tubulinqPCRR* | AATGGGGTATTACTGAGAGC | |
| ID423248*qPCRF* | AGAGGATGGATTCAGAGAGC | qPCR primers for gene ID423248 |
| ID423248*qPCRR* | GCAACGCGCGACGTTCTTCC | |

**Fluorescent tagging primers**

| Name | Sequence (5′ to 3′) |
|---|---|
| 24GFP5F | AACAGCTATGACATGATTACGTATTCTGCTCTTCTTGAACC |
| 24GFP5R | GCTCCTCGCCCTTGCTCACTACCTTGTTTCCTCCCCTCAGG |
| 24GFPGFPF | GTGAGCAAGGGCGAGGAGC |
| 24GFPGFPR | TAAAACGACGGCCAGTGCCAGAATTCGGTTGTTGGTGCTGGTGG |
| 24GFPHYGF | CACCAGCACCAACAACCGCTGGGATTGCCCCTCGATGC |
| 24GFPHYGR | GCTTACTATACATGCATATTTCCTACTGAACGTTATGAC |
| 24GFP3F | AAATATGCATGTATAGTAAGC |
| 24GFP3R | AAAACGACGGCCAGTGCCAGGCCCAGCCCATTACTGCAGC |
| 69GFP5F | AACAGCTATGACATGATTACGAACAAGACGGCGACAAATCC |
| 69GFP5R | TCTAGCCGGATCCTAACTGCAAAACTCGAGAC |
| 69GFP3F | GCAGTTAGGATCCGGCTAGAGGAGCAGAGGTTGG |
| 69GFP3R | GTAAAACGACGGCCAGTGCCAGCAGATGGAGGAGCCATAGC |
| 69GFPHYGF | GTCTCGAGTTTTGCAGTTAGTGAGCAAGGGCGAGGAGCTG |
| 69GFPHYGR | AACCTCTGCTCCTCTAGCCGCCTACTGAACGTTATGAC |
| Erg11F | TCGAAACCTAATCAATCAACATGGGGTTGCTCTCCGCTGTGC |
| Erg11R | GCTCACCATTGCTTTTTCGGACAGTTGACG |
| McherryErg11F | CCGAAAAAGCAATGGTGAGCAAGGGCGAGG |
| McherryErg11R | TGCTCATAGTCACATCCCTCACTTGTACAGCTCGTCCATGC |
| AP57 | TCGAAACCTAATCAATCAACATGGTGAGCAAGGGCGAGG |
| AP58 | TGCTCATAGTCACATCCCTCAAAGCTTAGACTTGTACAGCTCGTCCATGC |

**Primers used for RIP mutation of *vdtX***

| Oligo nucleotide | Sequence | Notes |
|---|---|---|
| RIPF1 | CCTCTGCAGGTCGACTCTAGACGATGCTAGATCACAAGTCC | Amplification and cloning of 2258 bp of *vdtX* |
| RIPR2 | GGCCAATTCTTAATTAAGATATCGGCGAGAAATGATGTTTTCC | |

| | | |
|---|---|---|
| RIPseqF | AATTTCCAACGATGCTCAGC | Amplification and sequencing of the native *vdtX* allele in the region targeted for RIP |
| RIPseqR | TTGGAACGAATCAGACAAGG | |

### Figure S1 UV-vis-spectra and mass spectra for 6, 10, 11, and 12

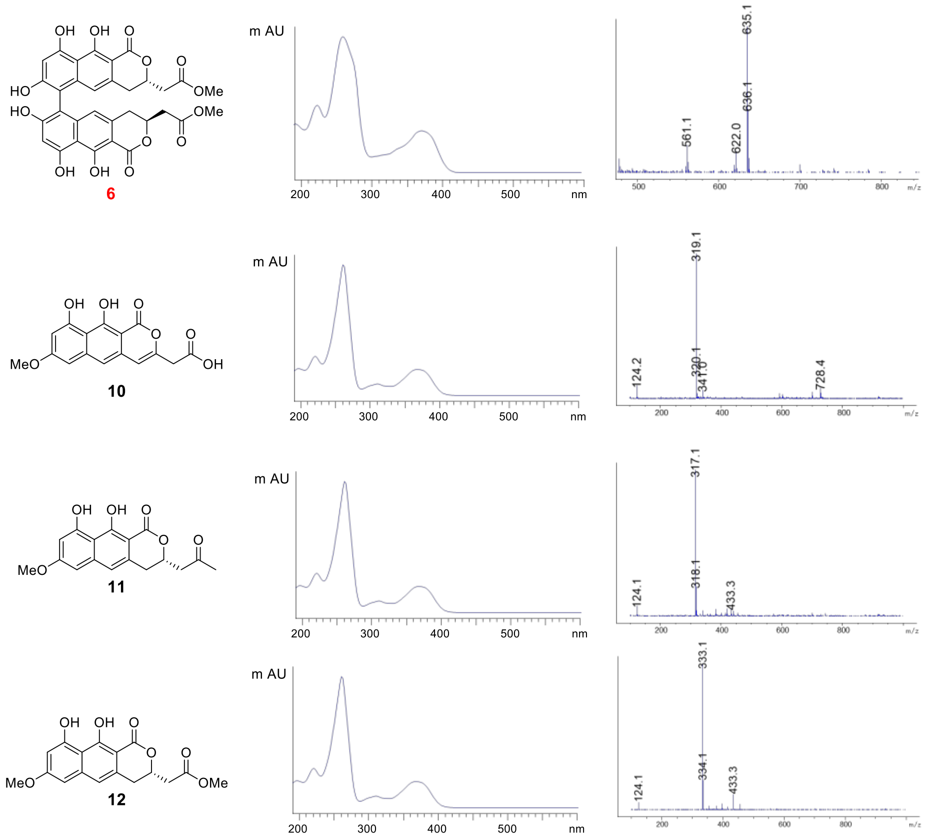

**Figure S2. Mutation in *vdtX* via repeat induced point mutation.** The top sequence is from wild type and the bottom from the RIP mutant.

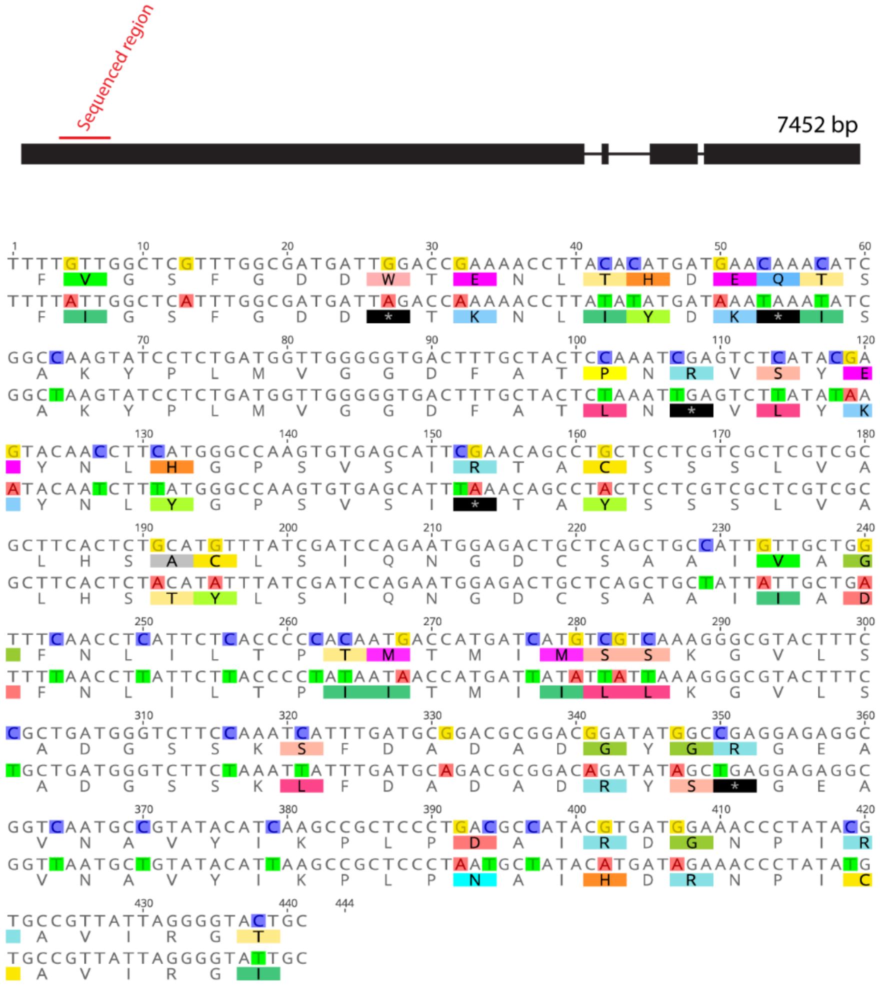

**Figure S3. $^{1}H$ NMR spectrum (500 MHz) of 1 in chloroform-*d***

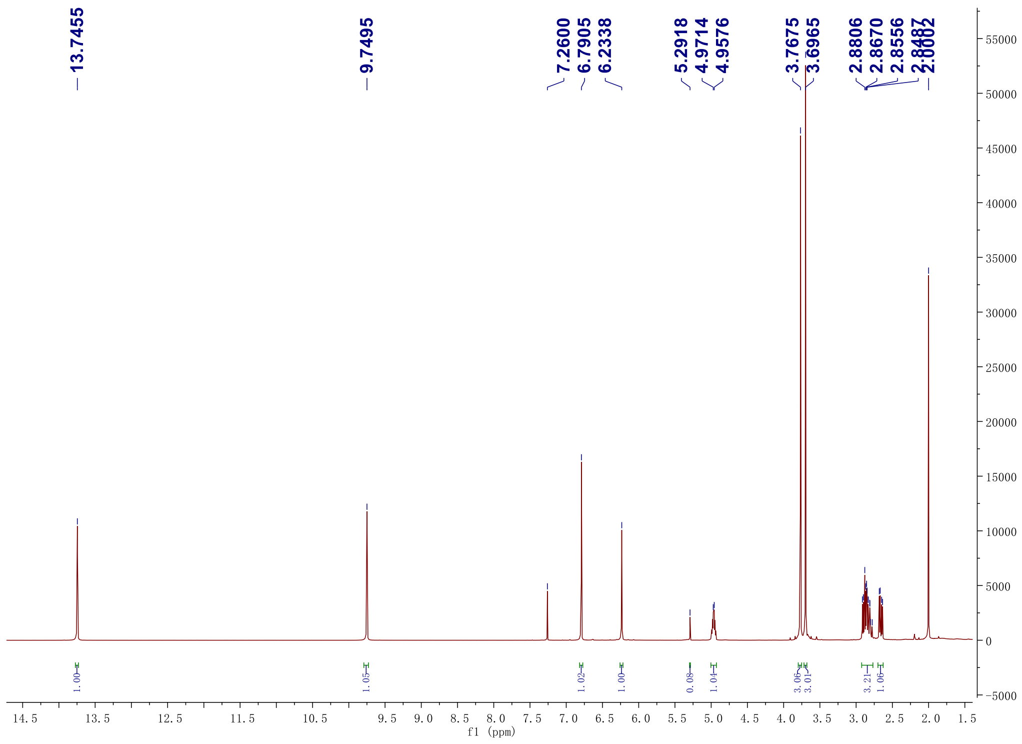

#### Figure S4. $^{13}C$ NMR spectrum (125 MHz) of 1 in chloroform-*d*

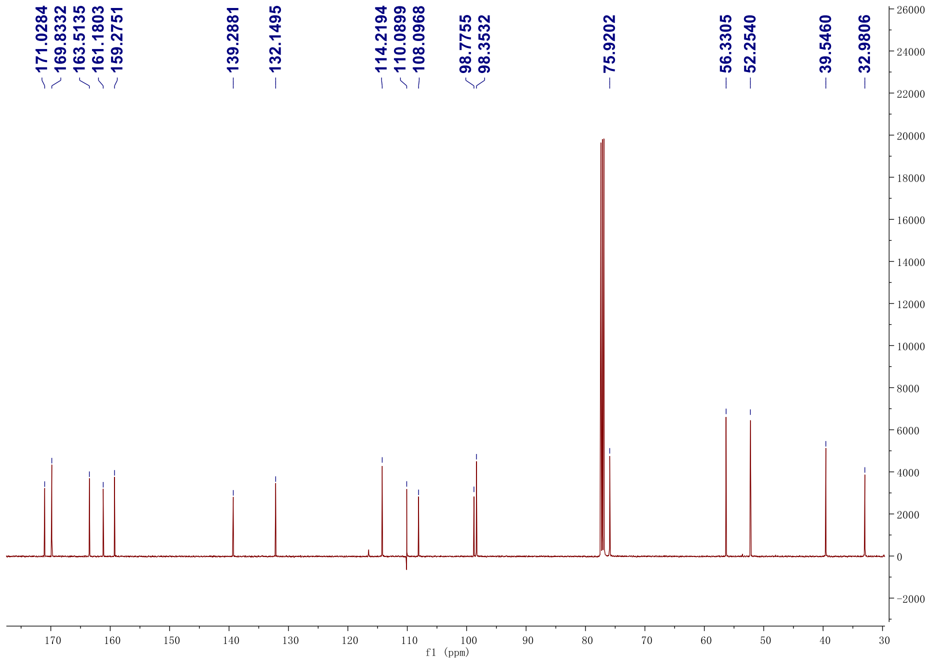

#### Figure S5. DEPT-135 $^{13}C$ NMR spectrum (125 MHz) of 1 in chloroform-*d*

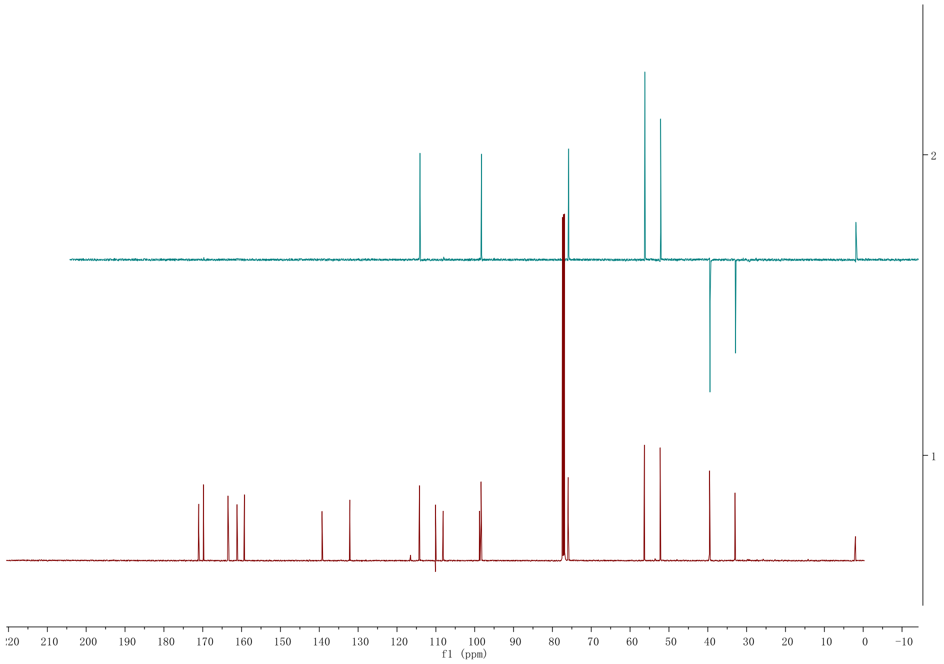

**Figure S6. $^1H$-$^1H$ gCOSY NMR spectrum (500 MHz) of 1 in chloroform-*d***

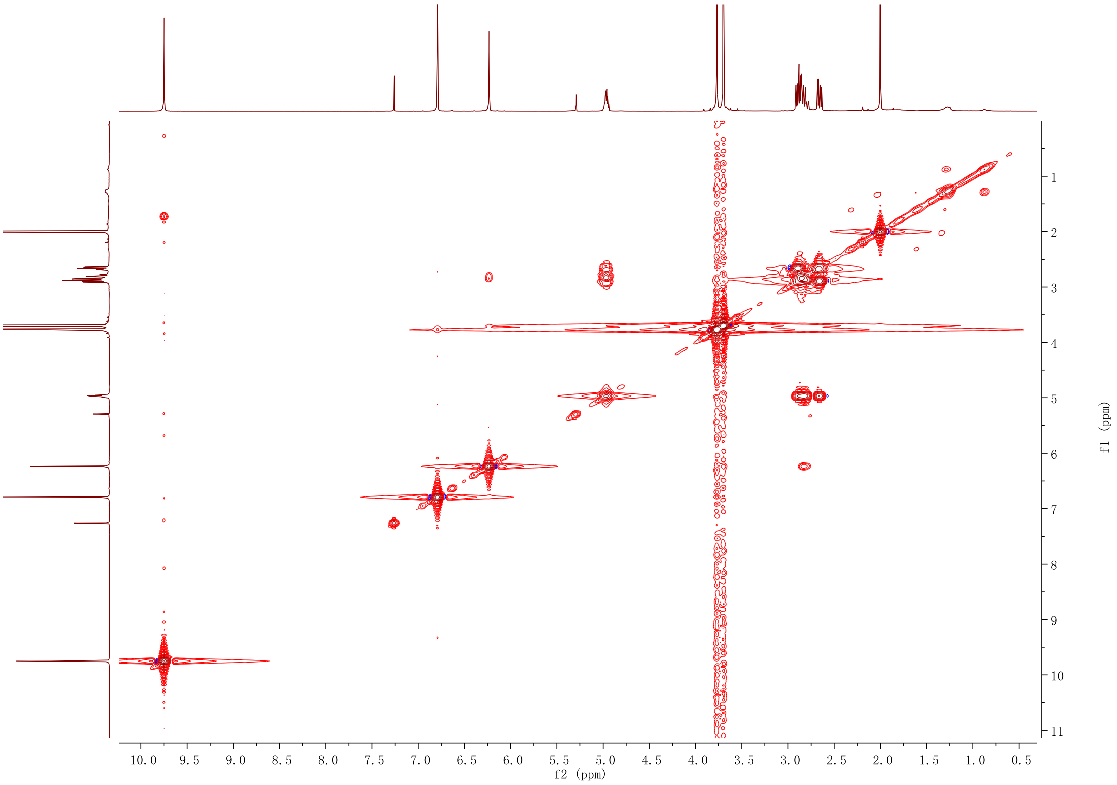

**Figure S7. HSQC NMR spectrum (500 MHz) of 1 in chloroform-*d***

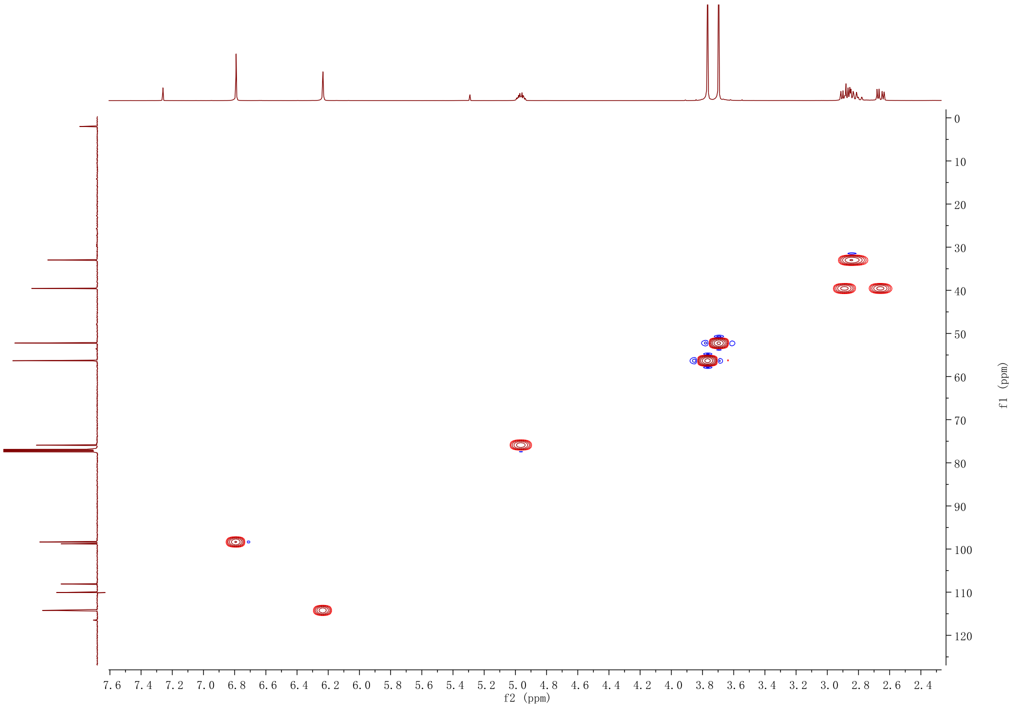

**Figure S8. HMBC NMR spectrum (500 MHz) of 1 in chloroform-*d***

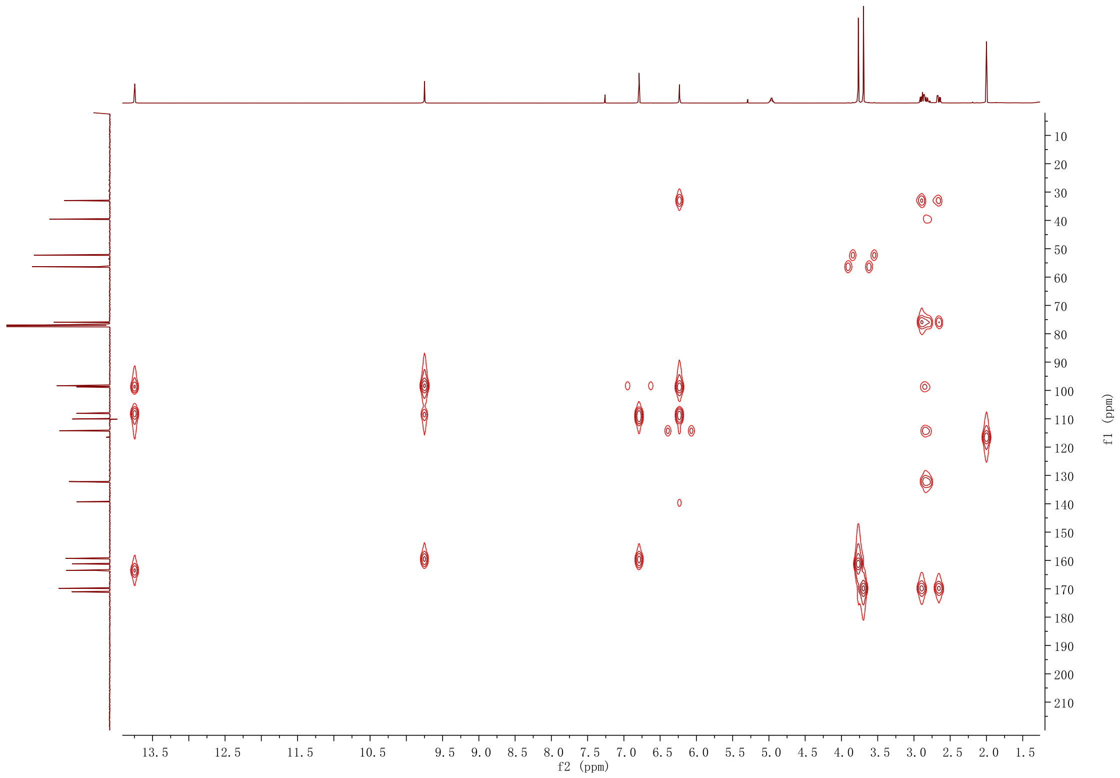

**Figure S9. $^{1}H$ NMR spectrum (500 MHz) of 1′ in chloroform-*d***

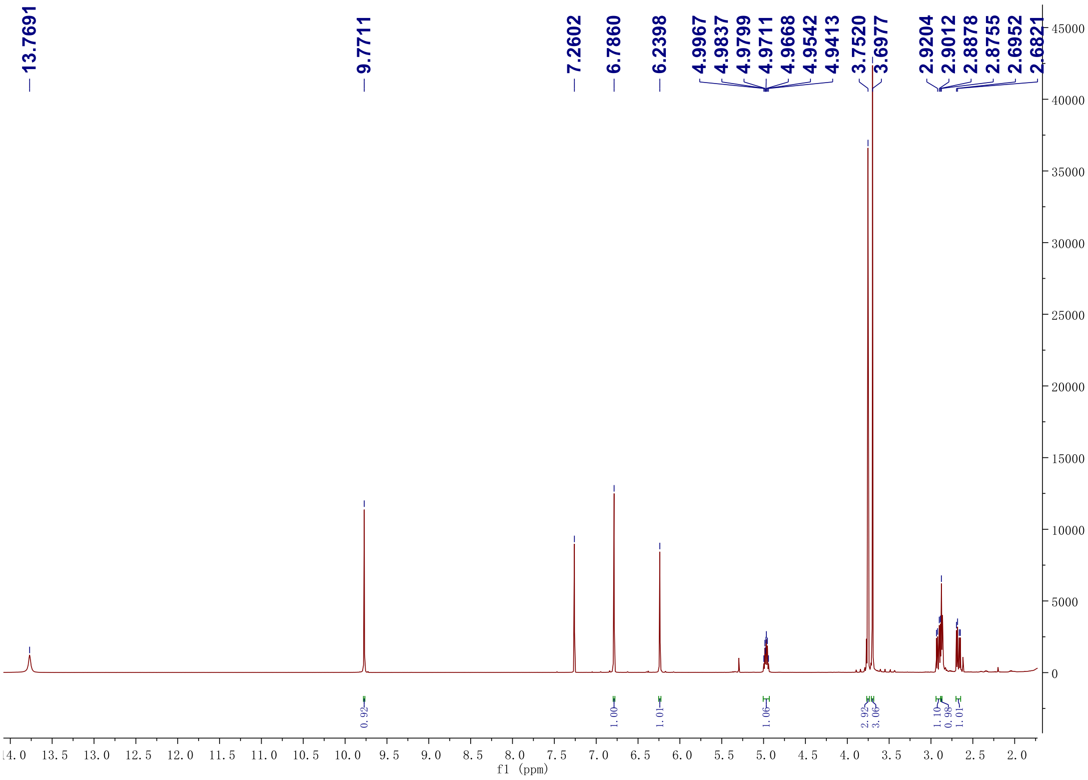

**Figure S10. $^{13}C$ NMR spectrum (125 MHz) of 1′ in chloroform-*d***

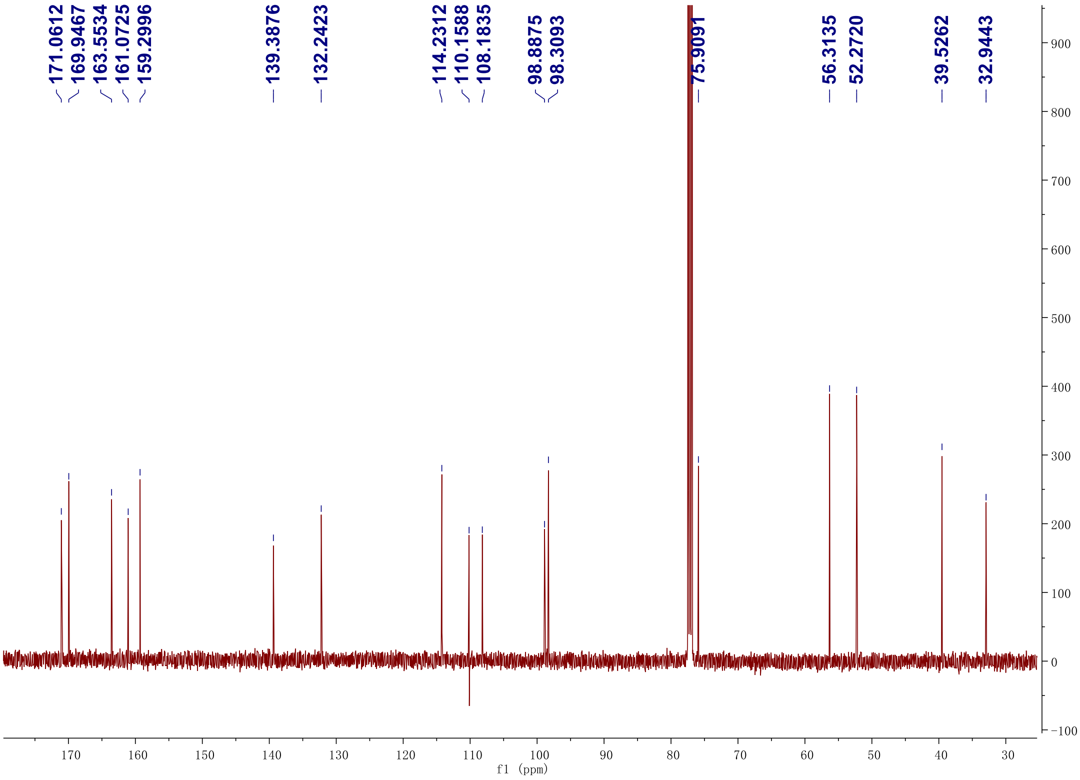

**Figure S11. DEPT-135 $^{13}C$ NMR spectrum (125 MHz) of 1′ in chloroform-*d***

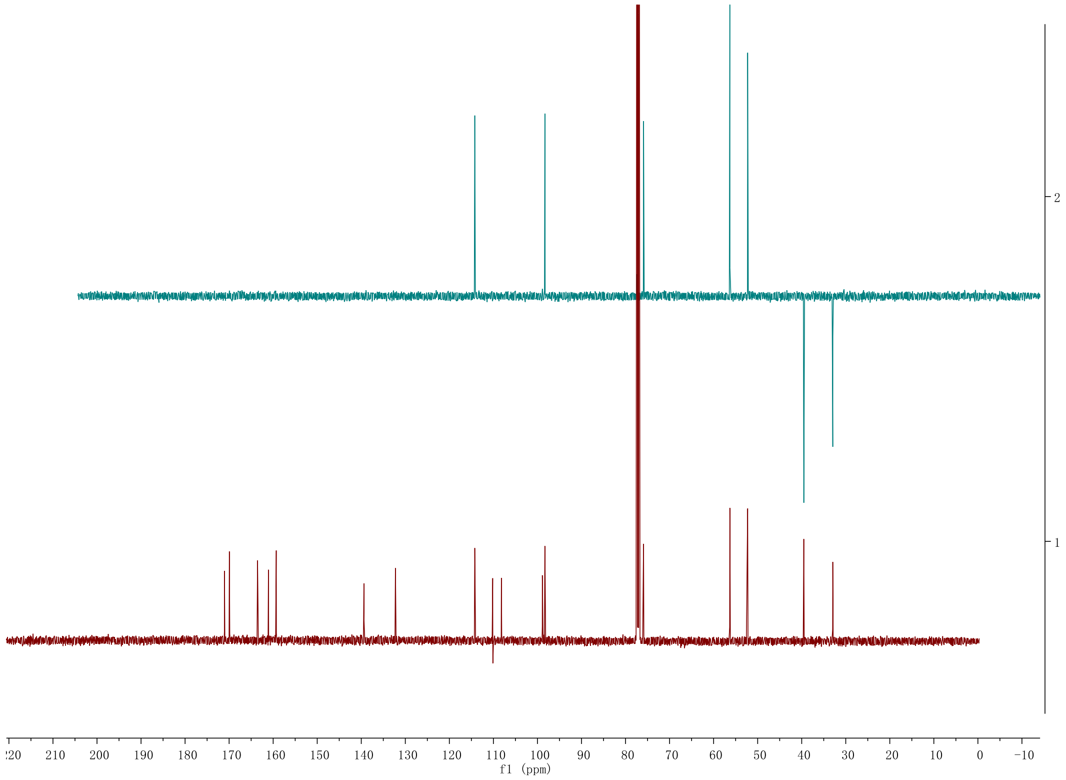

**Figure S12. $^{1}H$-$^{1}H$ gCOSY NMR spectrum (500 MHz) of 1′ in chloroform-*d***

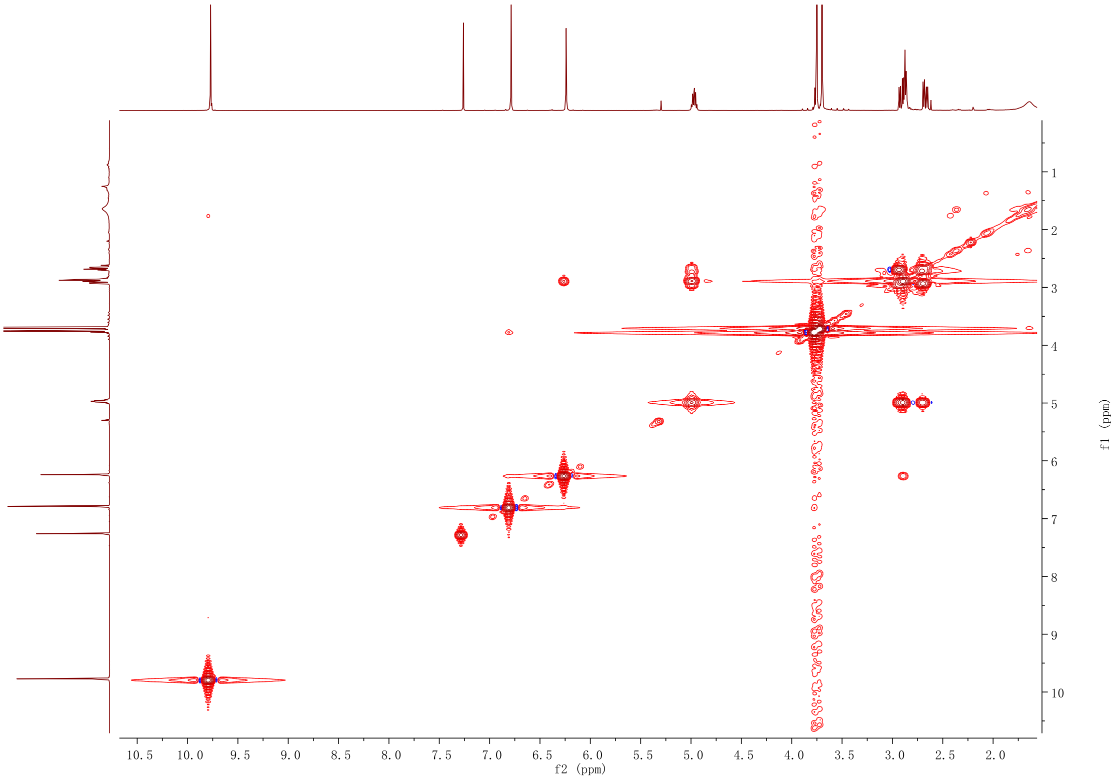

Figure S13. HSQC NMR spectrum (500 MHz) of 1′ in chloroform-*d*

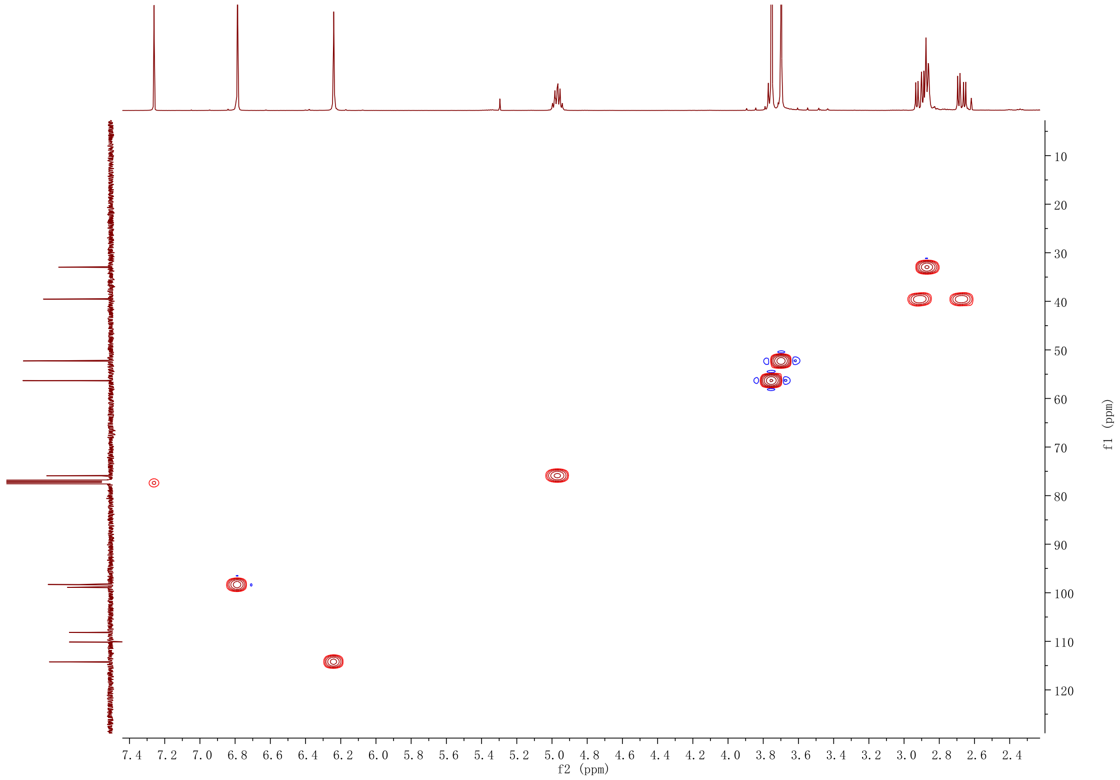

**Figure S14. HMBC NMR spectrum (500 MHz) of 1′ in chloroform-*d***

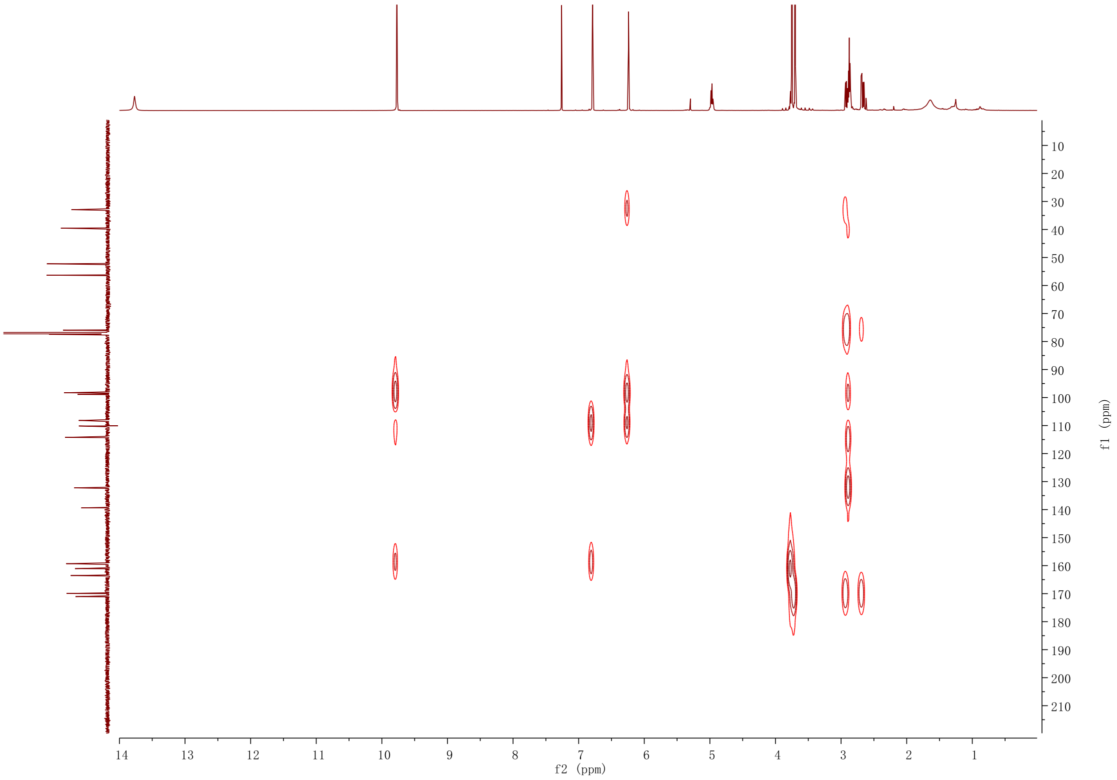

**Figure S15. [1]H NMR spectrum (500 MHz) of 2 in acetonitrile-$d_3$**

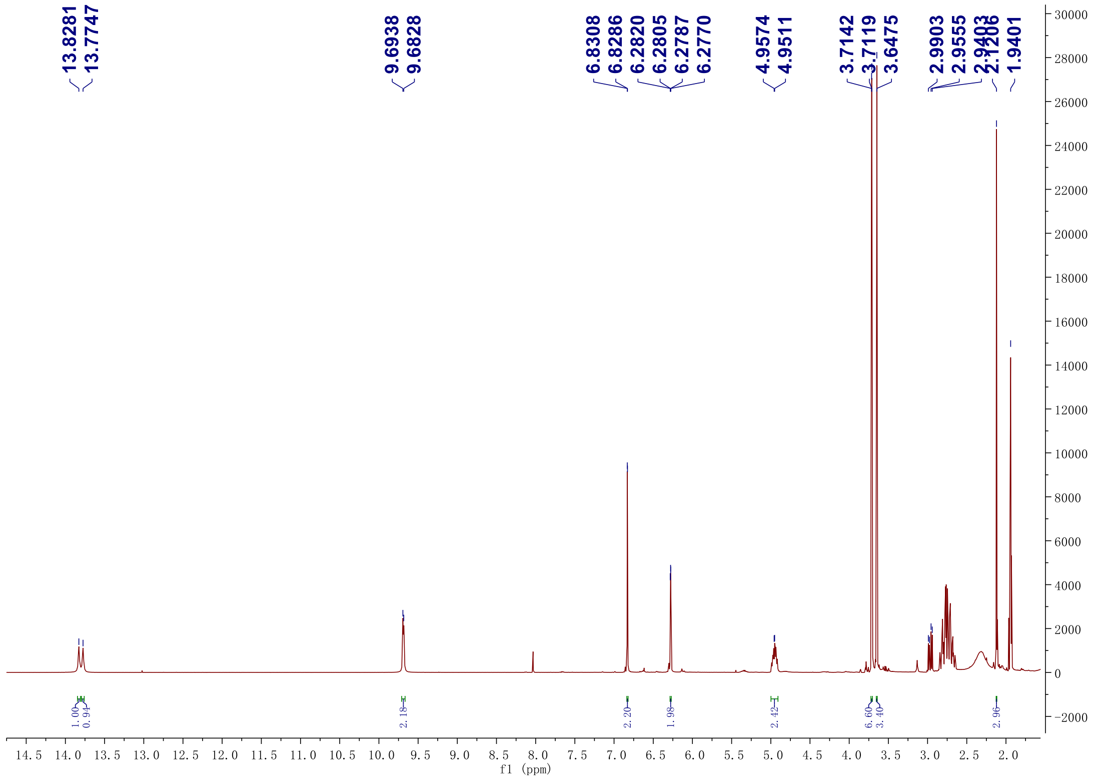

**Figure S16. [13]C NMR spectrum (125 MHz) of 2 in acetonitrile-$d_3$**

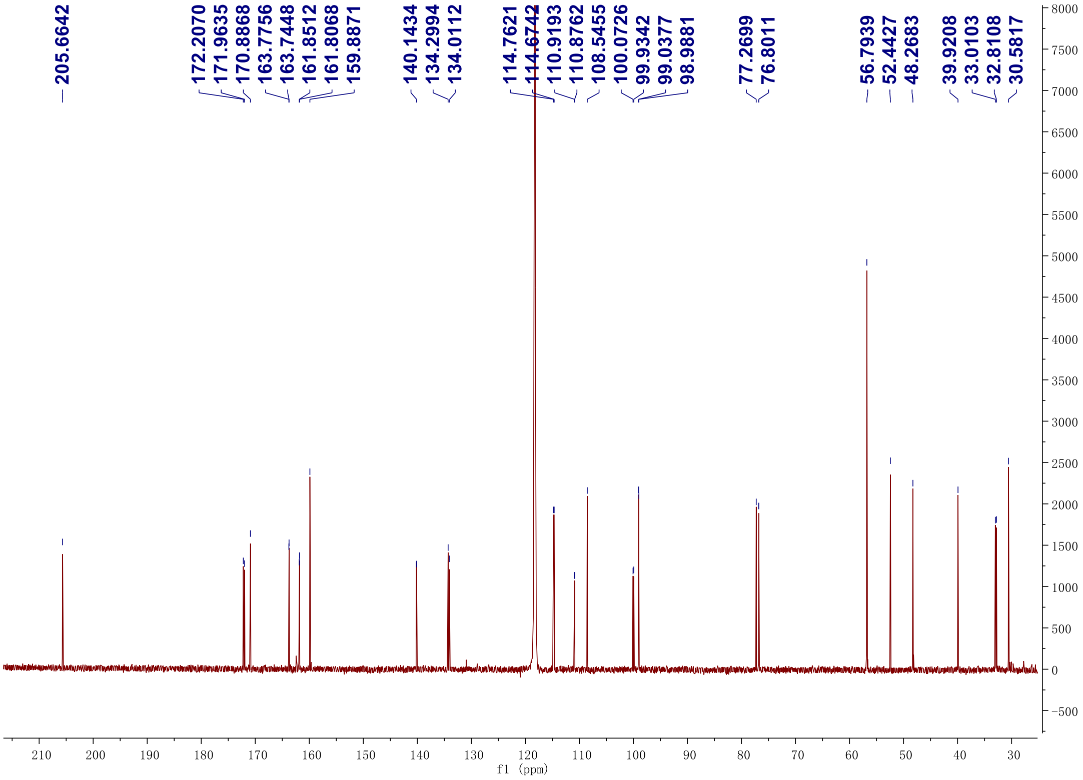

**Figure S17. DEPT-135 and DEPT-90 $^{13}C$ NMR spectrum (125 MHz) of 2 in acetonitrile-$d_3$**

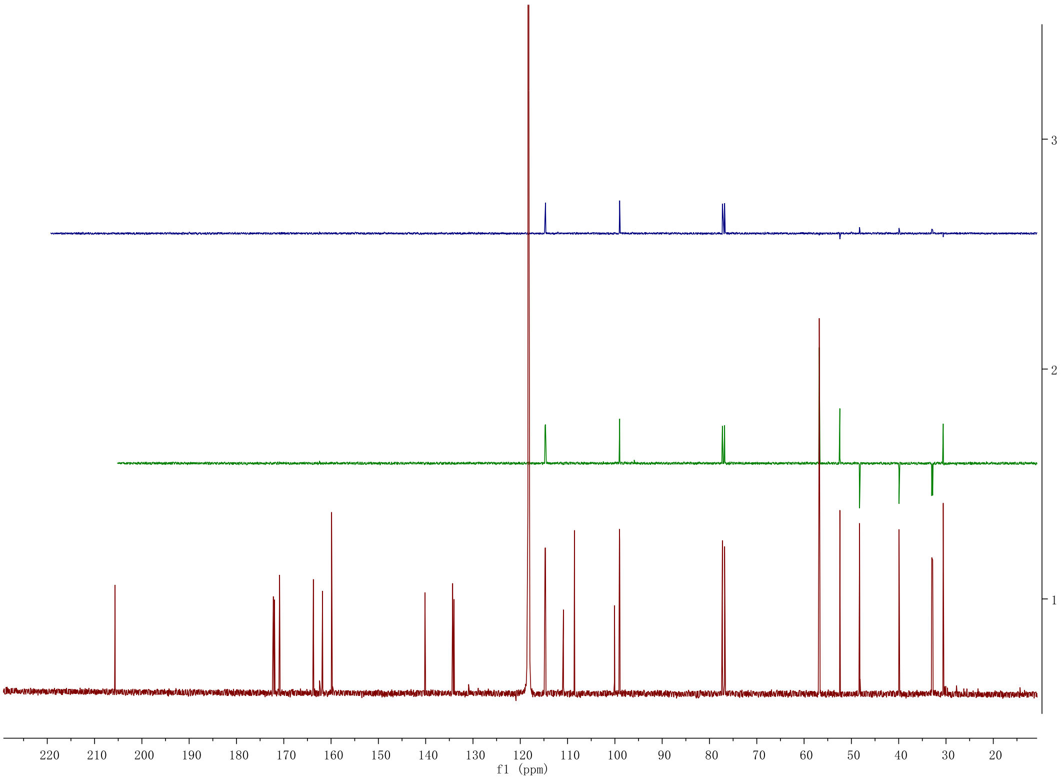

**Figure S18. [1]H-[1]H gCOSY NMR spectrum (500 MHz) of 2 in acetonitrile-$d_3$**

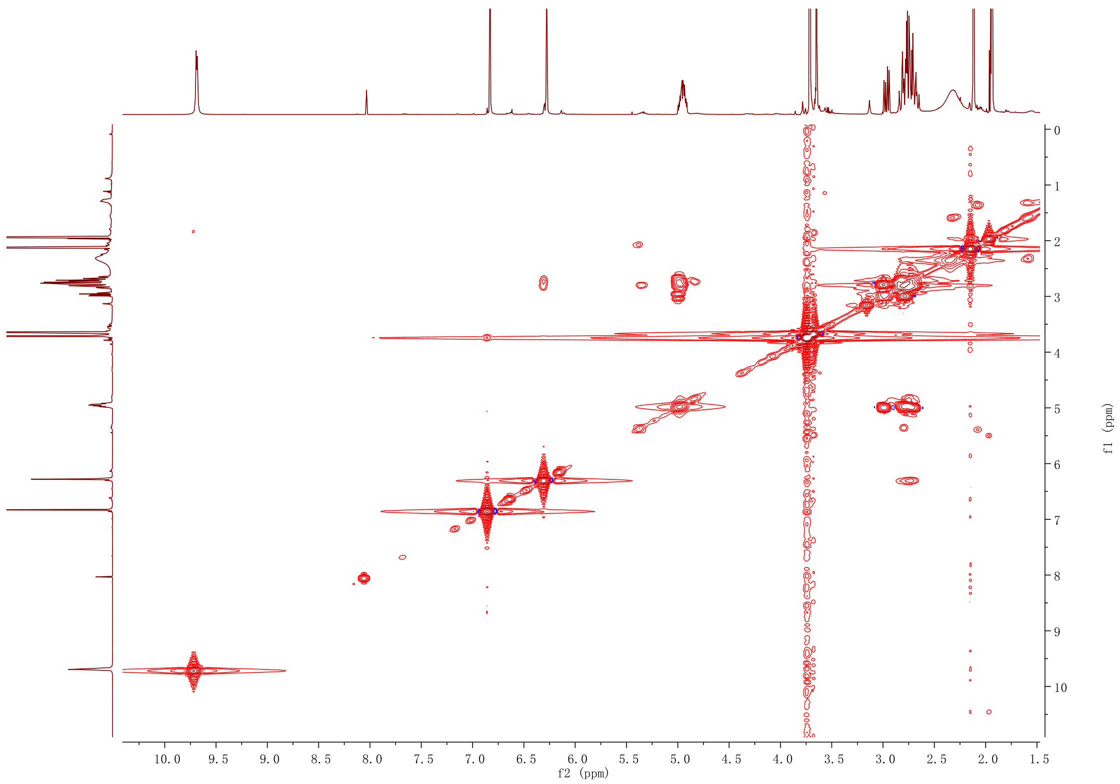

**Figure S19. HSQC NMR spectrum (500 MHz) of 2 in acetonitrile-$d_3$**

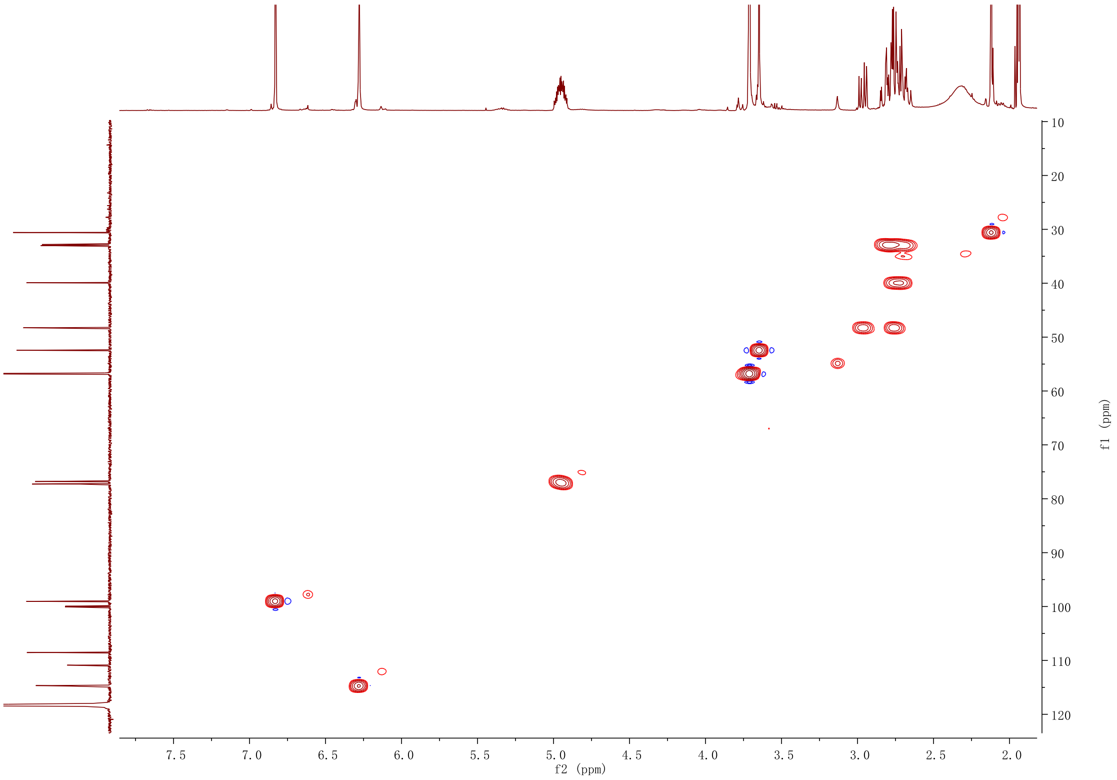

**Figure S20. HMBC NMR spectrum (500 MHz) of 2 in acetonitrile-$d_3$**

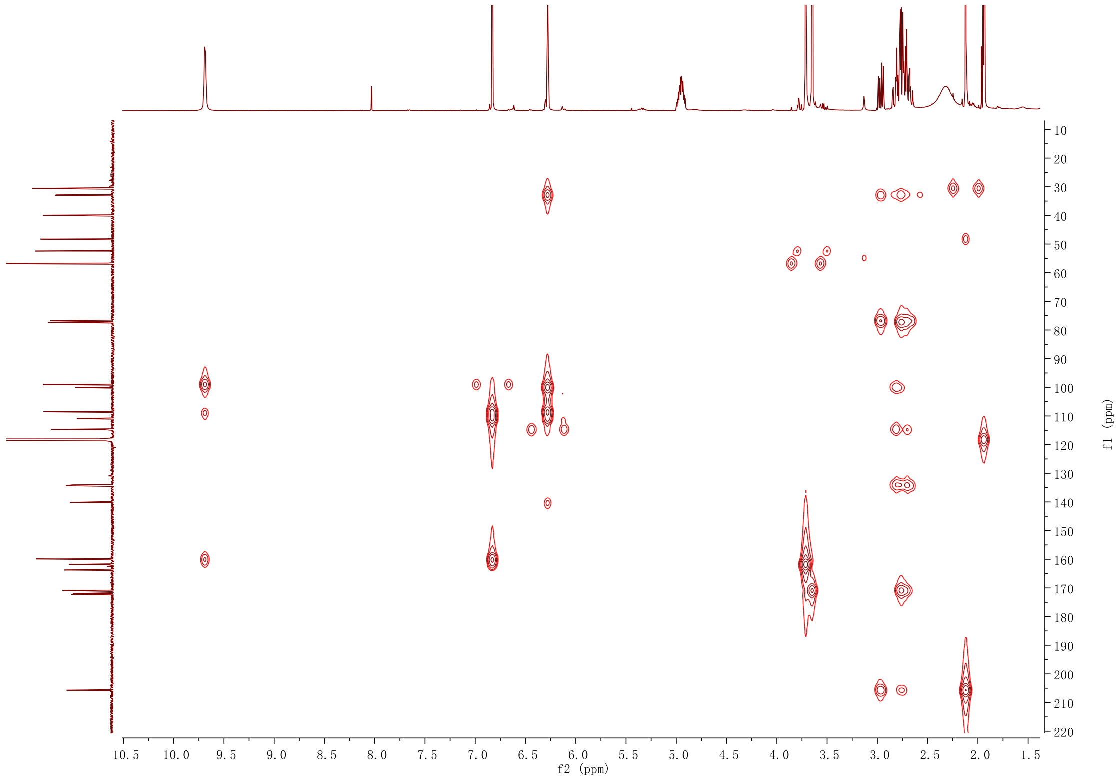

**Figure S21. $^1H$ NMR spectrum (600 MHz) of 3 in acetonitrile-$d_3$**

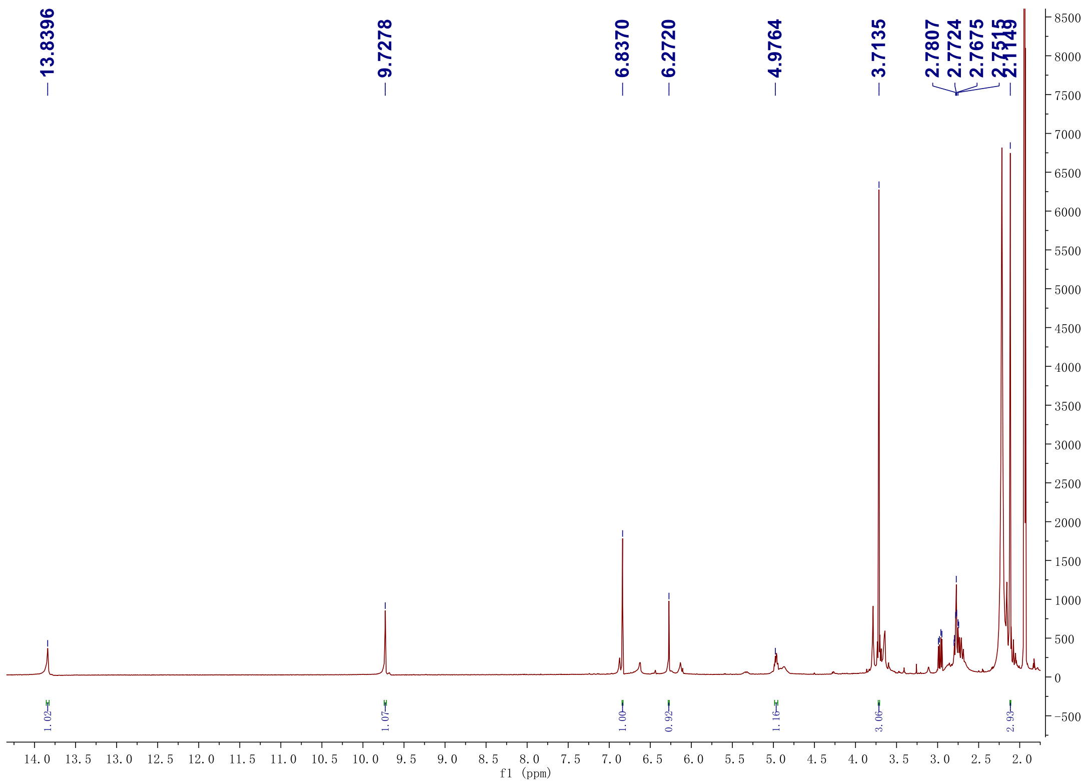

**Figure S22. $^{13}C$ NMR spectrum (150 MHz) of 3 in acetonitrile-$d_3$**

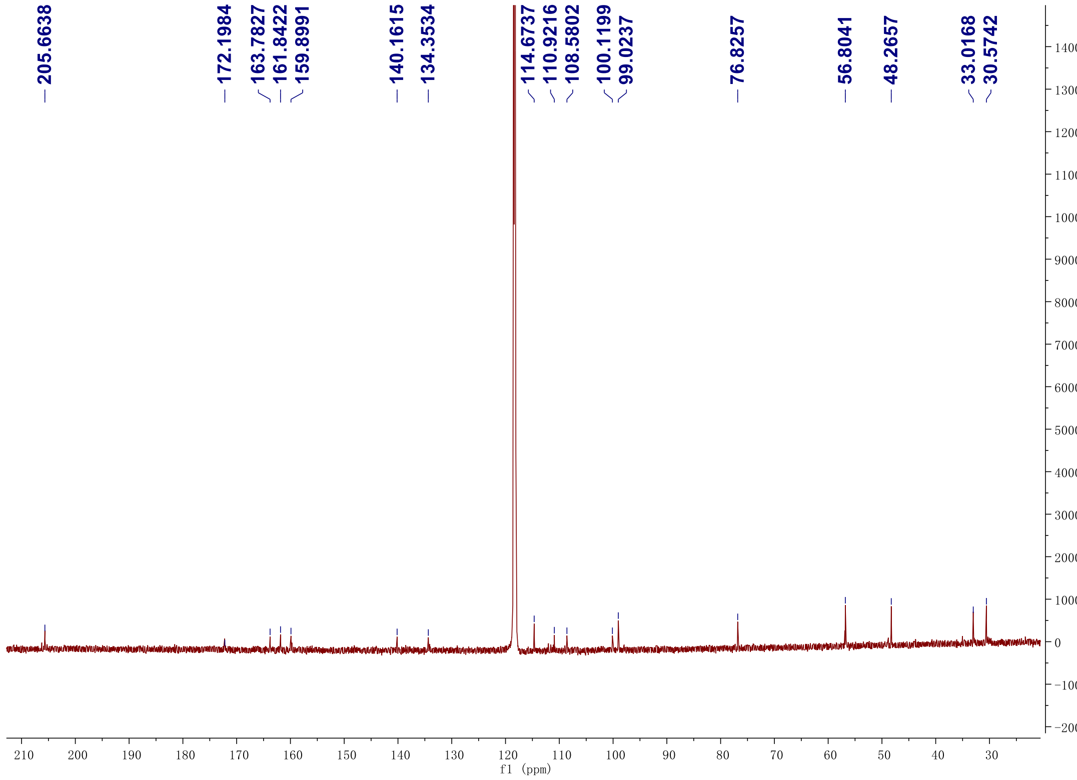

**Figure S23. DEPT-135 $^{13}C$ NMR spectrum (150 MHz) of 3 in acetonitrile-$d_3$**

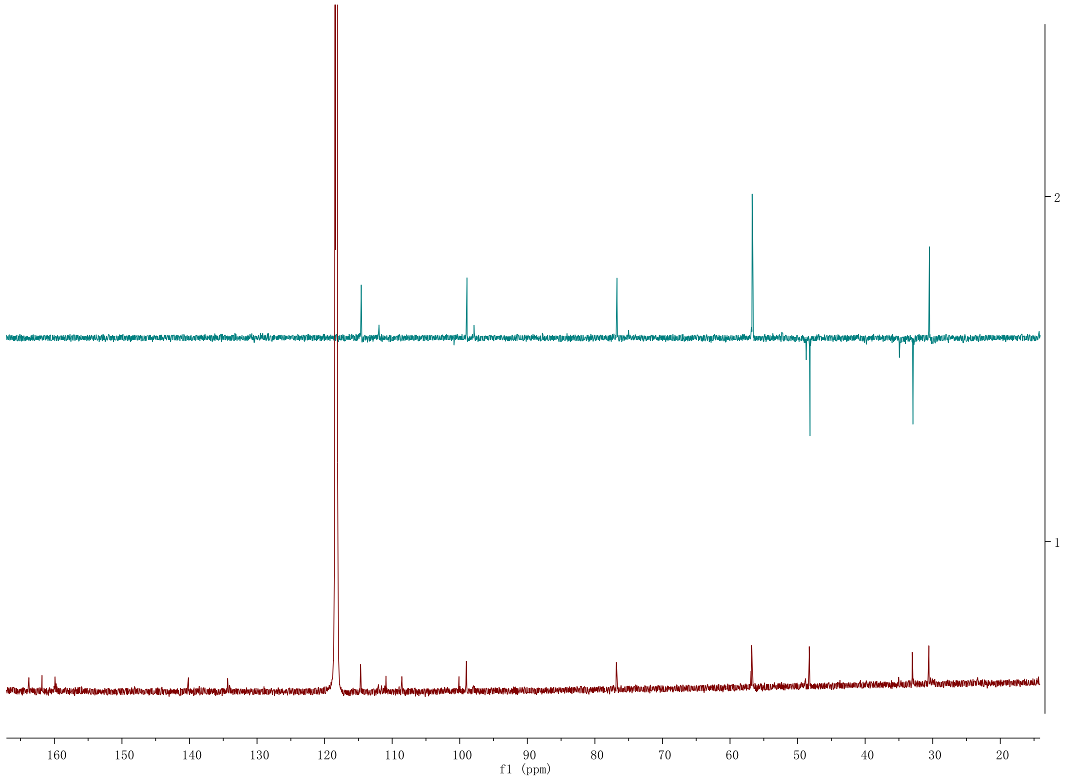

**Figure S24. $^{1}H$-$^{1}H$ gCOSY NMR spectrum (600 MHz) of 3 in acetonitrile-$d_3$**

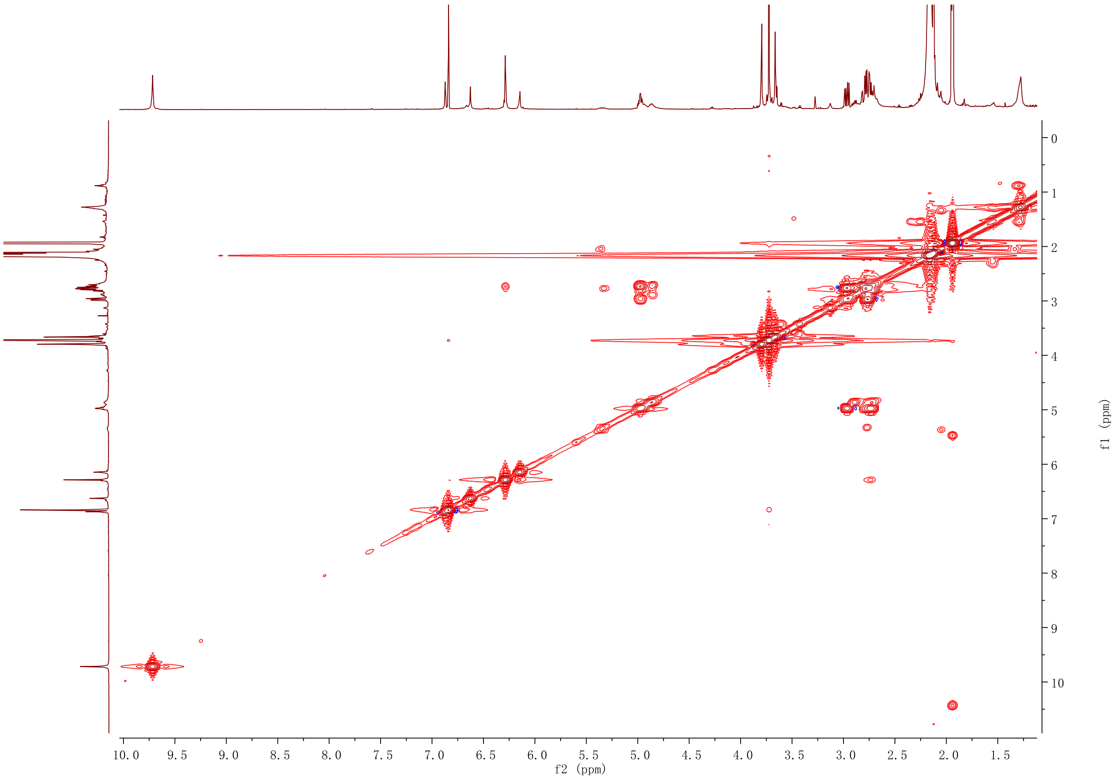

**Figure S25. HSQC NMR spectrum (600 MHz) of 3 in acetonitrile-$d_3$**

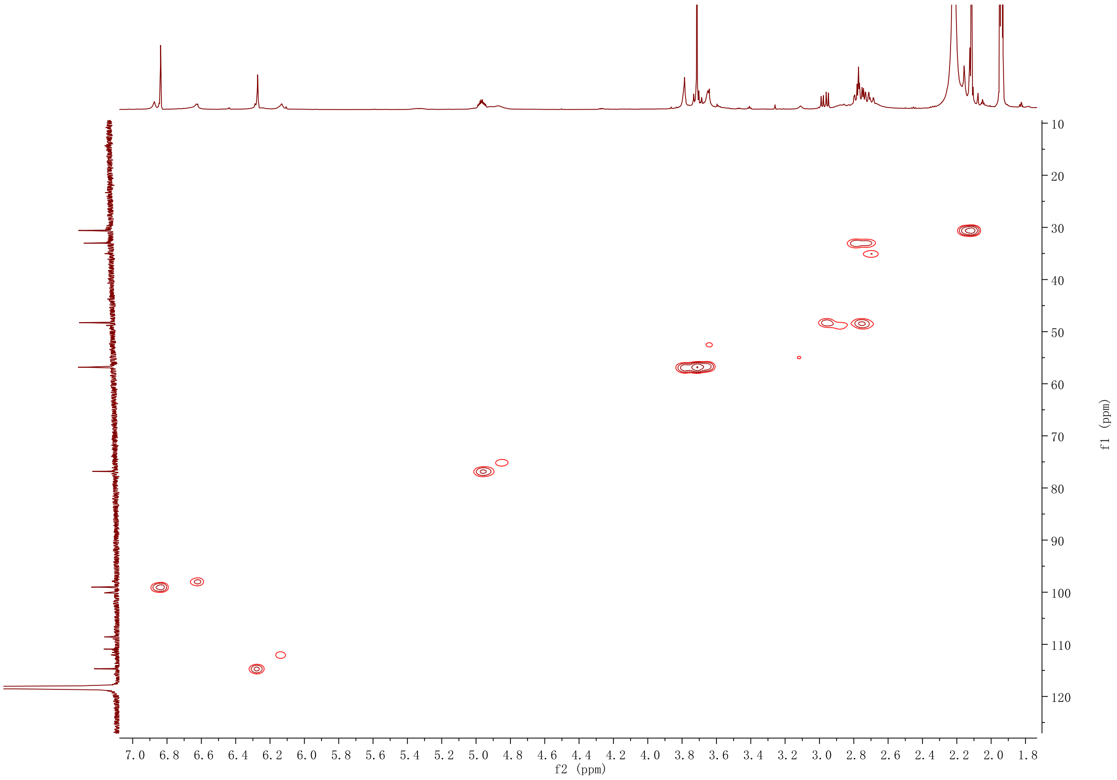

**Figure S26. HMBC NMR spectrum (600 MHz) of 3 in acetonitrile-$d_3$**

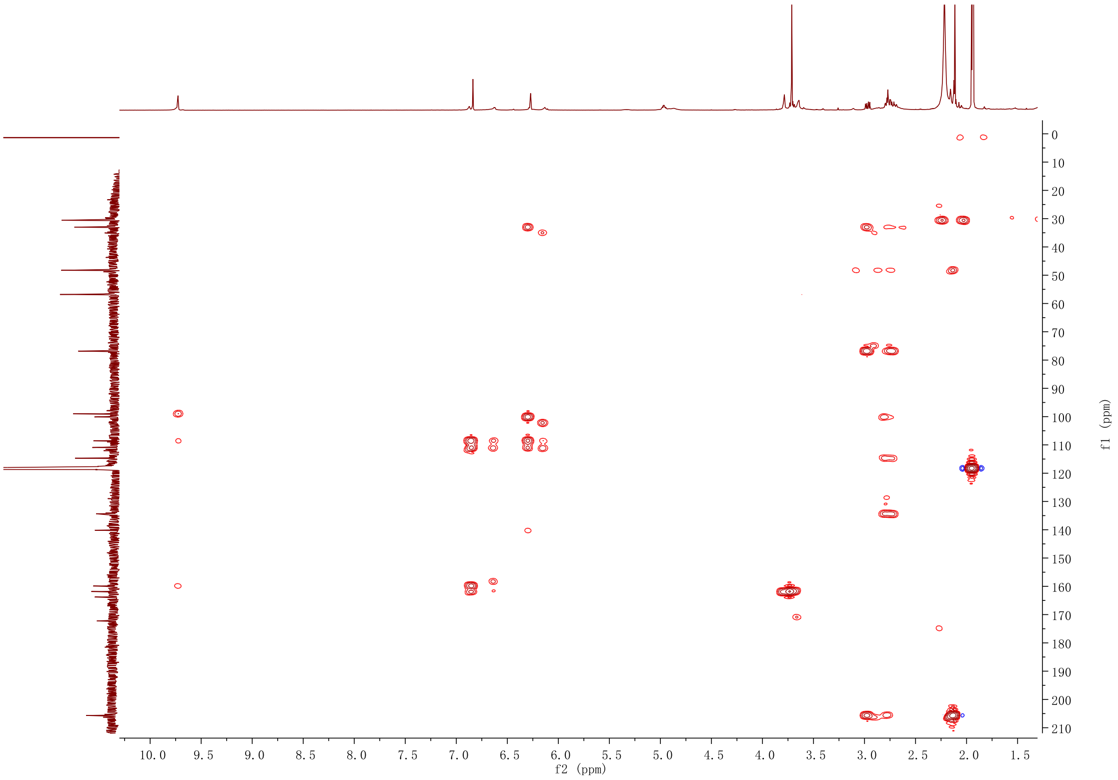

**Figure S27. $^{1}$H NMR spectrum (500 MHz) of 4 in acetonitrile-$d_3$**

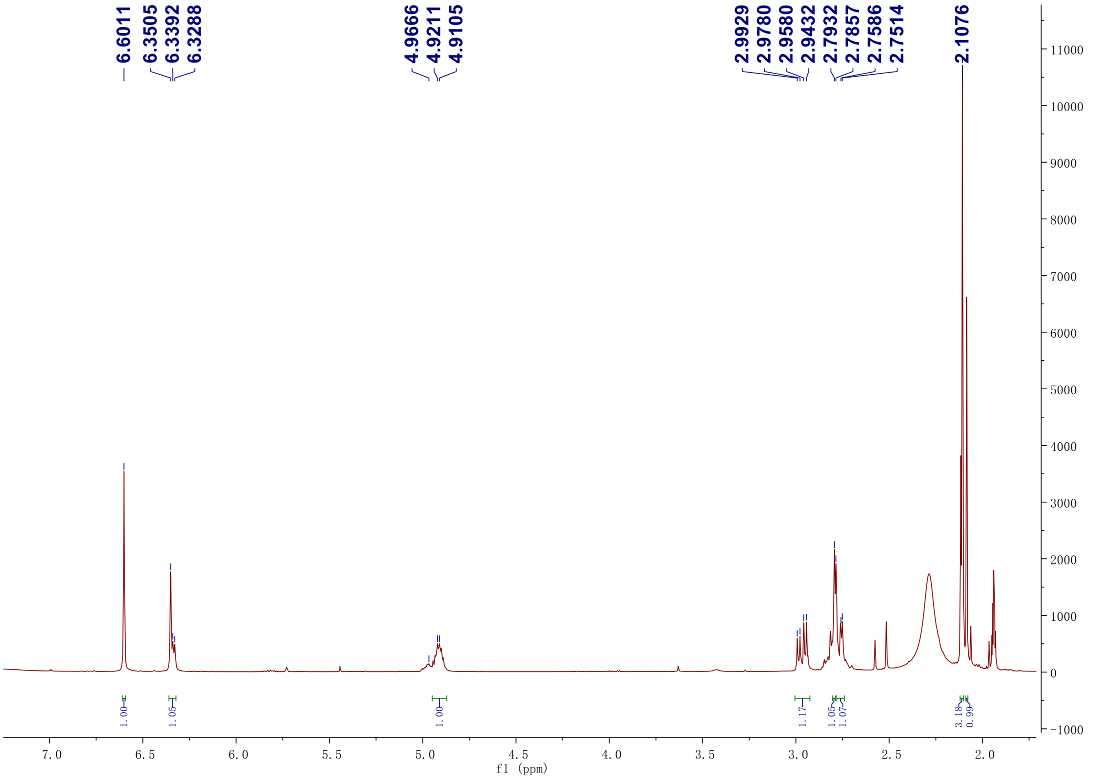

#### Figure S28. $^{13}C$ NMR spectrum (125 MHz) of 4 in acetonitrile-$d_3$

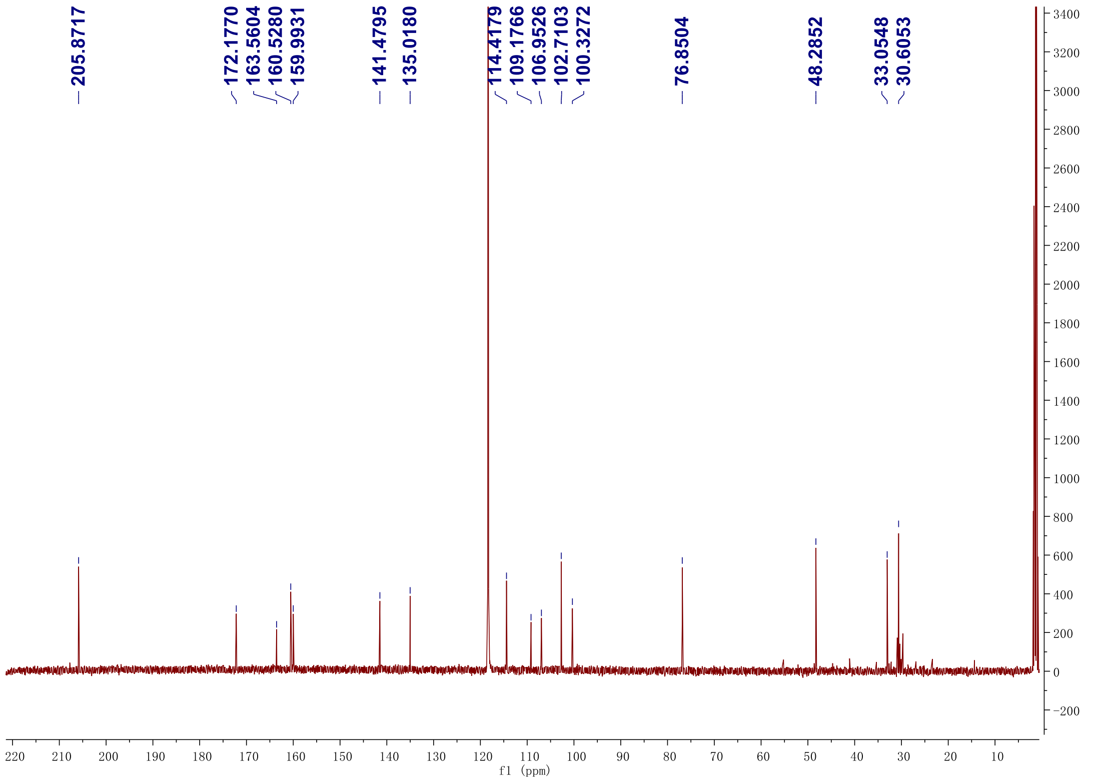

**Figure S29. DEPT-135 $^{13}$C NMR spectrum (125 MHz) of 4 in acetonitrile-*$d_3$***

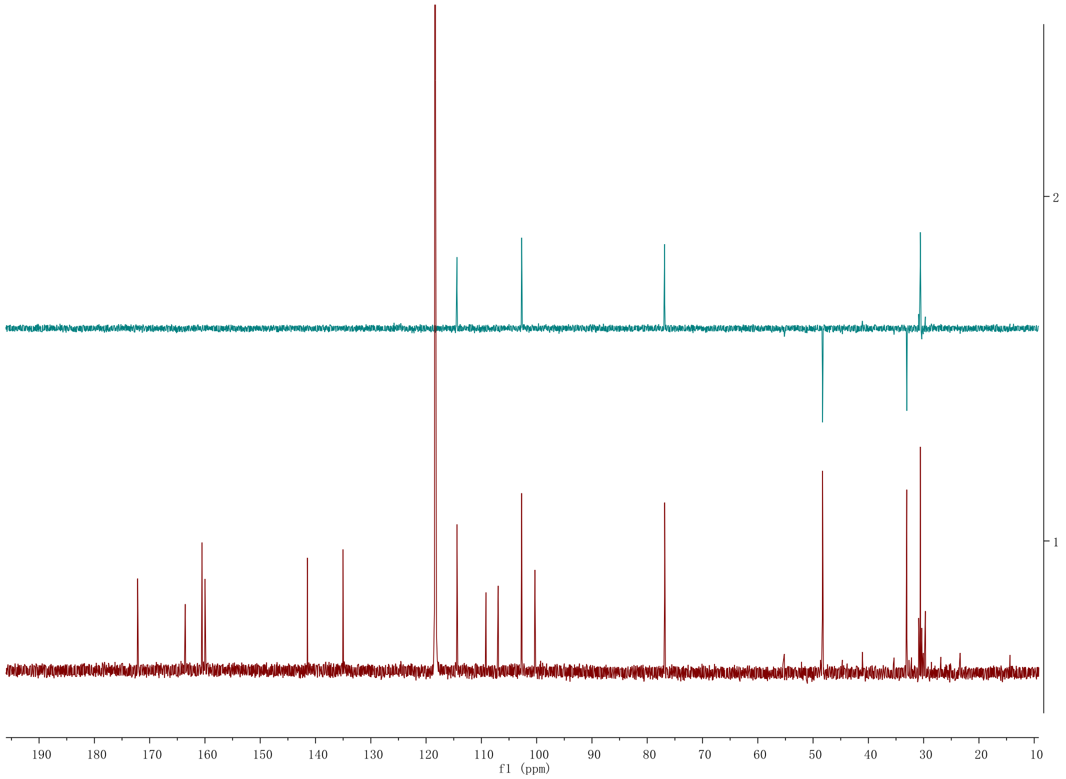

**Figure S30. $^1$H-$^1$H gCOSY NMR spectrum (500 MHz) of 4 in acetonitrile-$d_3$**

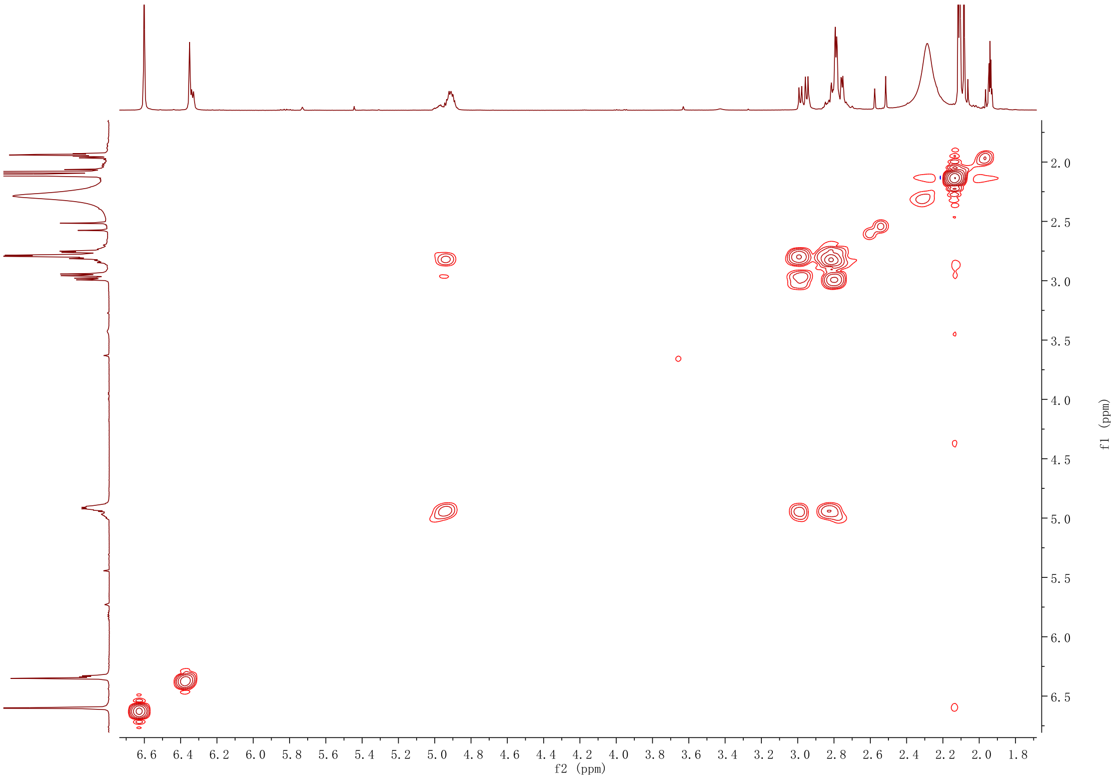

**Figure S31. HSQC NMR spectrum (500 MHz) of 4 in acetonitrile-$d_3$**

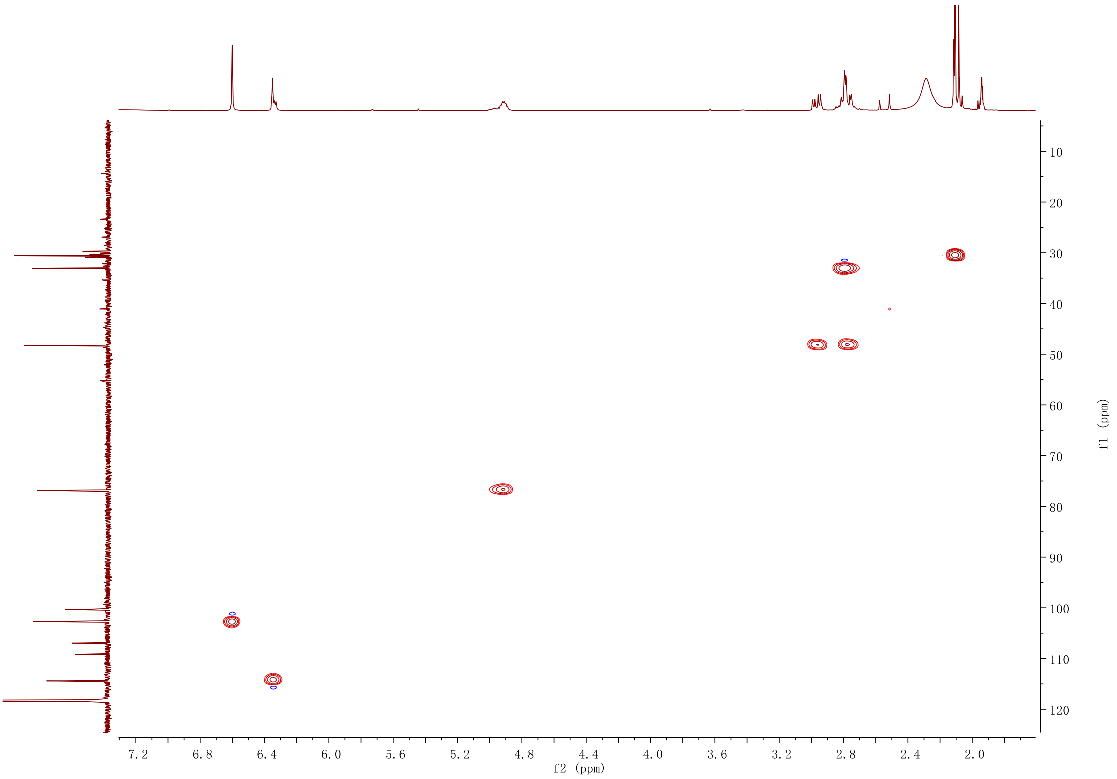

**Figure S32. HMBC NMR spectrum (500 MHz) of 4 in acetonitrile-$d_3$**

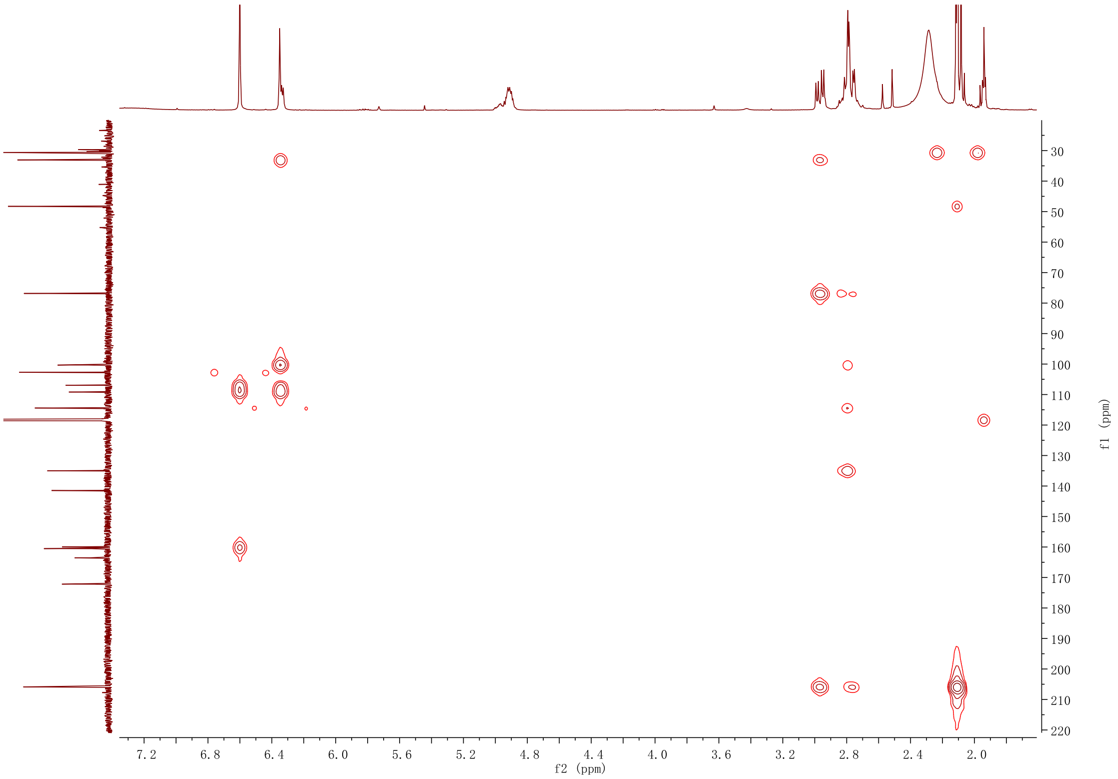

#### Figure S33. $^1$H NMR spectrum (600 MHz) of 5 in acetonitrile-$d_3$

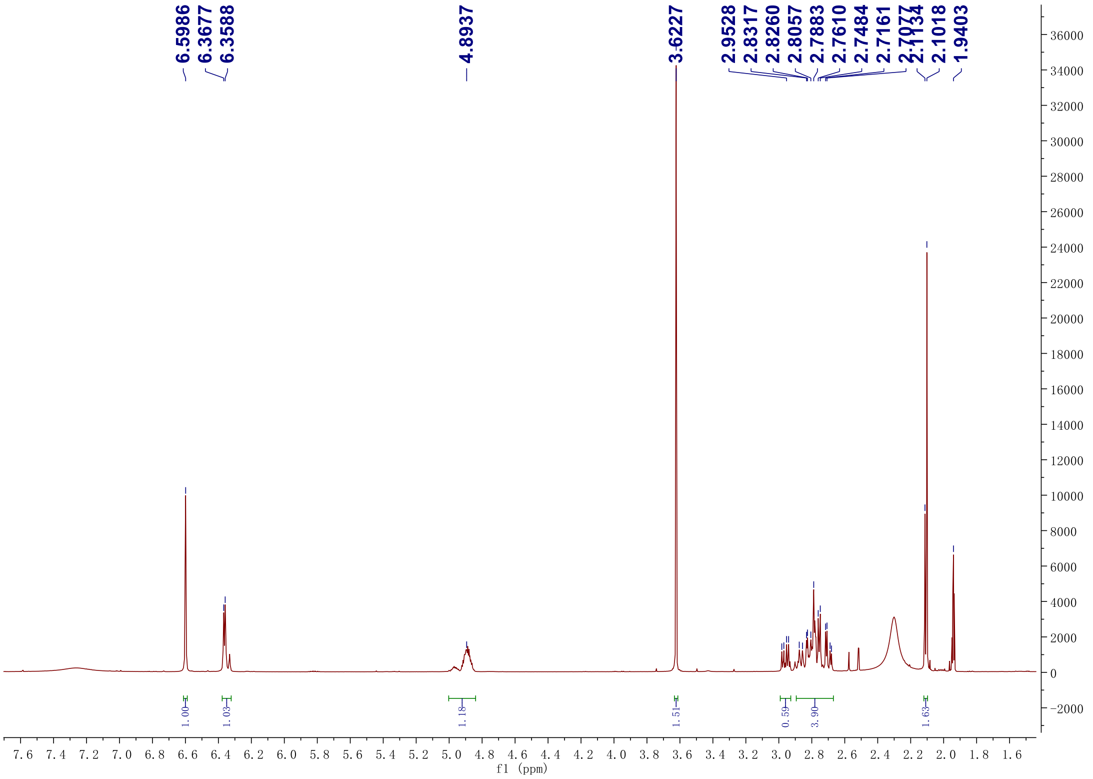

**Figure S34.** $^{13}C$ **NMR spectrum (150 MHz) of 5 in acetonitrile-*d*$_3$**

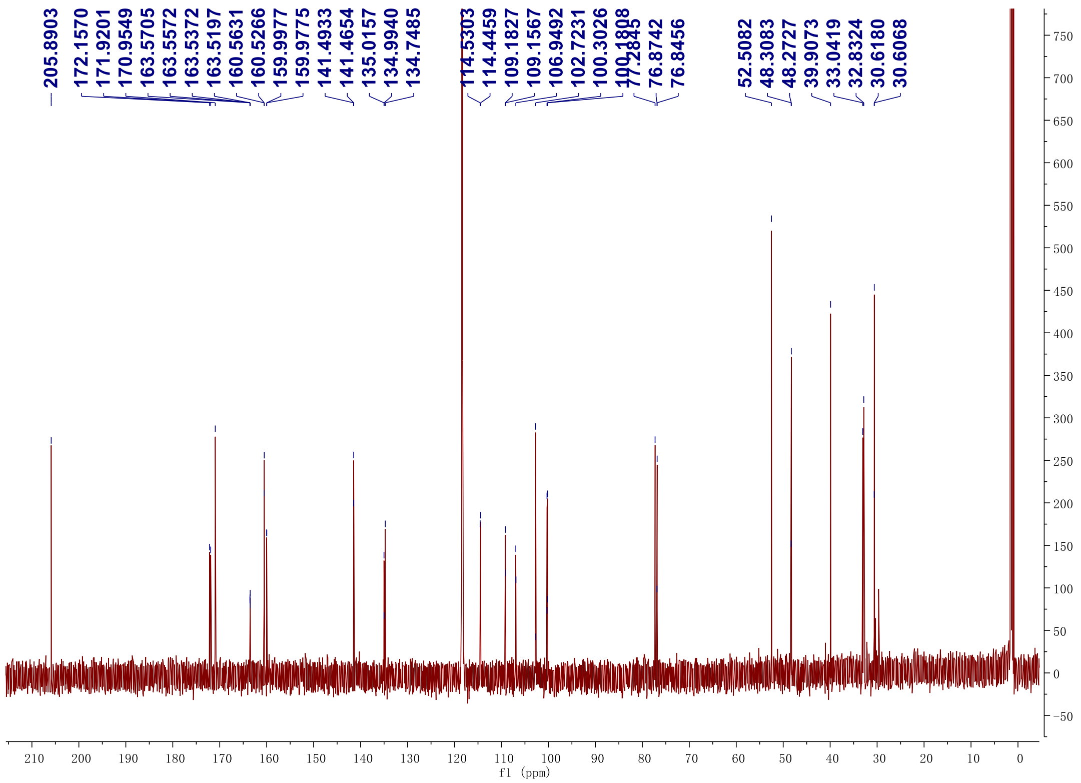

**Figure S35. DEPT-135 $^{13}$C NMR spectrum (150 MHz) of 5 in acetonitrile-$d_3$**

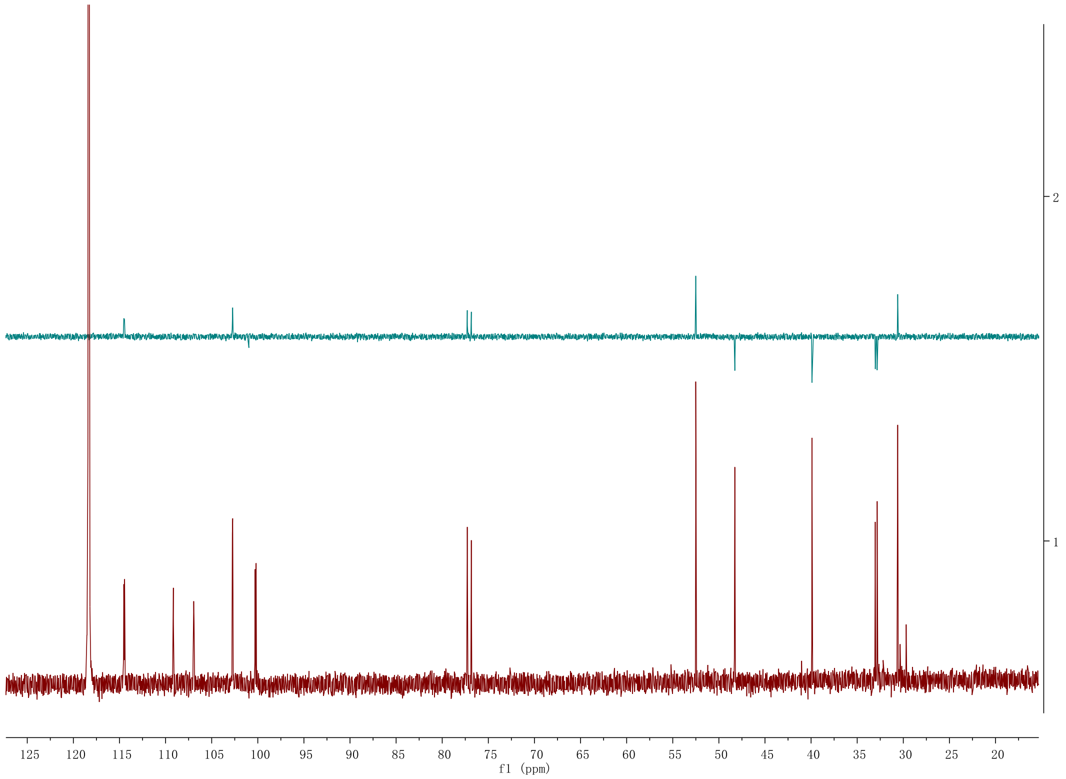

**Figure S36. $^1H$-$^1H$ gCOSY NMR spectrum (500 MHz) of 5 in acetonitrile-$d_3$**

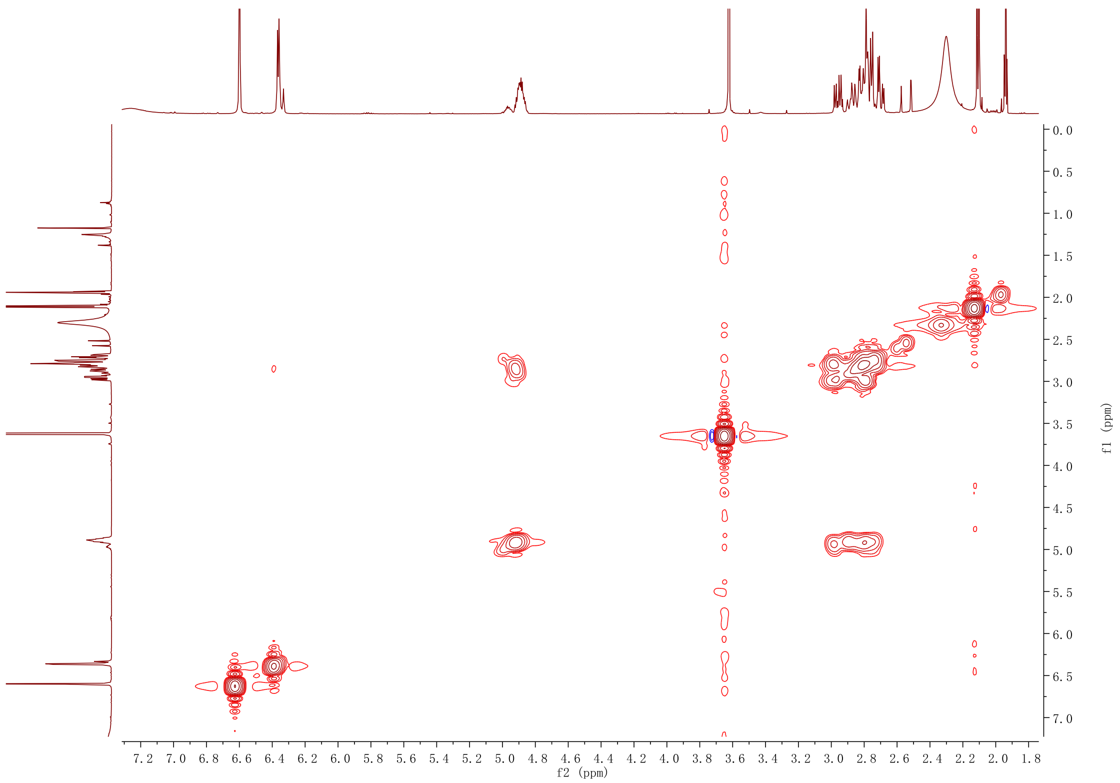

**Figure S37. HSQC NMR spectrum (600 MHz) of 5 in acetonitrile-$d_3$**

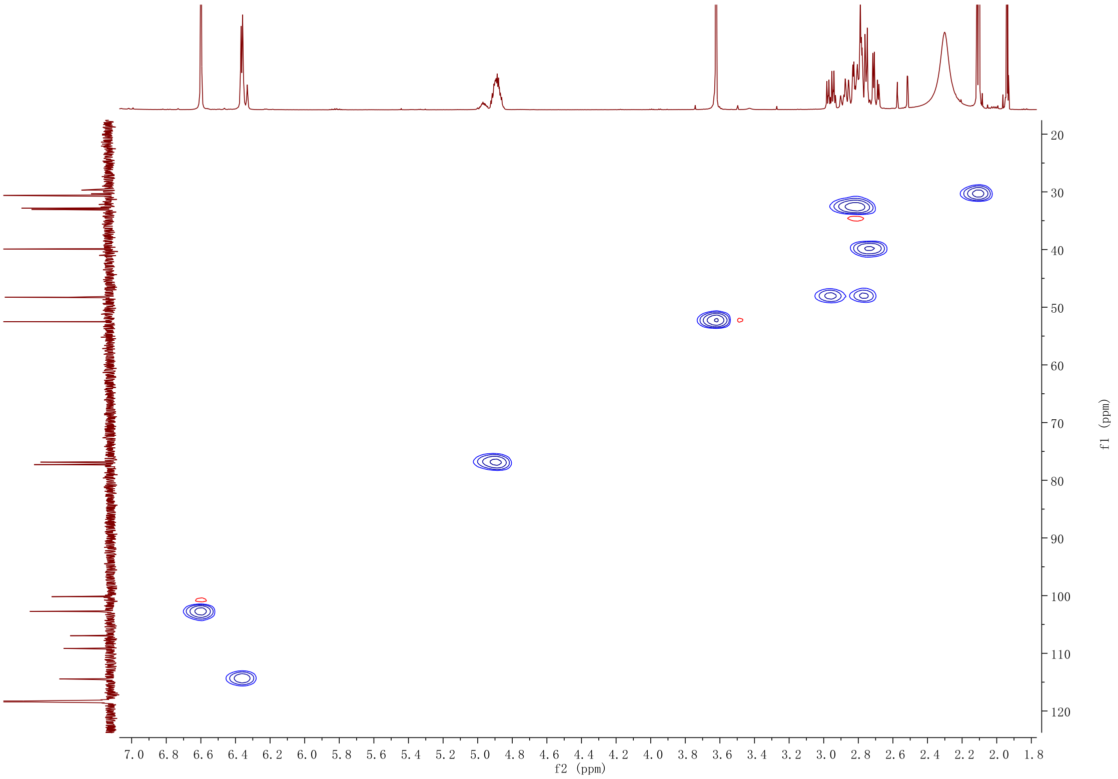

**Figure S38. HMBC NMR spectrum (600 MHz) of 5 in acetonitrile-$d_3$**

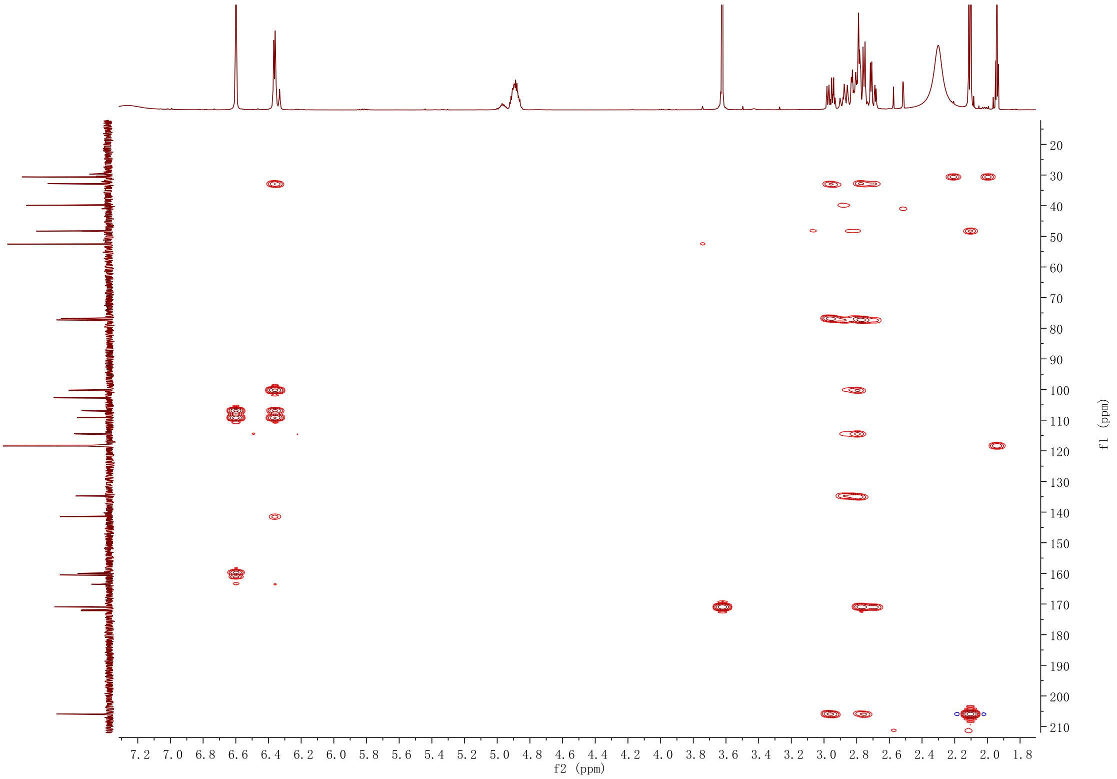

**Figure S39. [1]H NMR spectrum (500 MHz) of 7 in chloroform-*d***

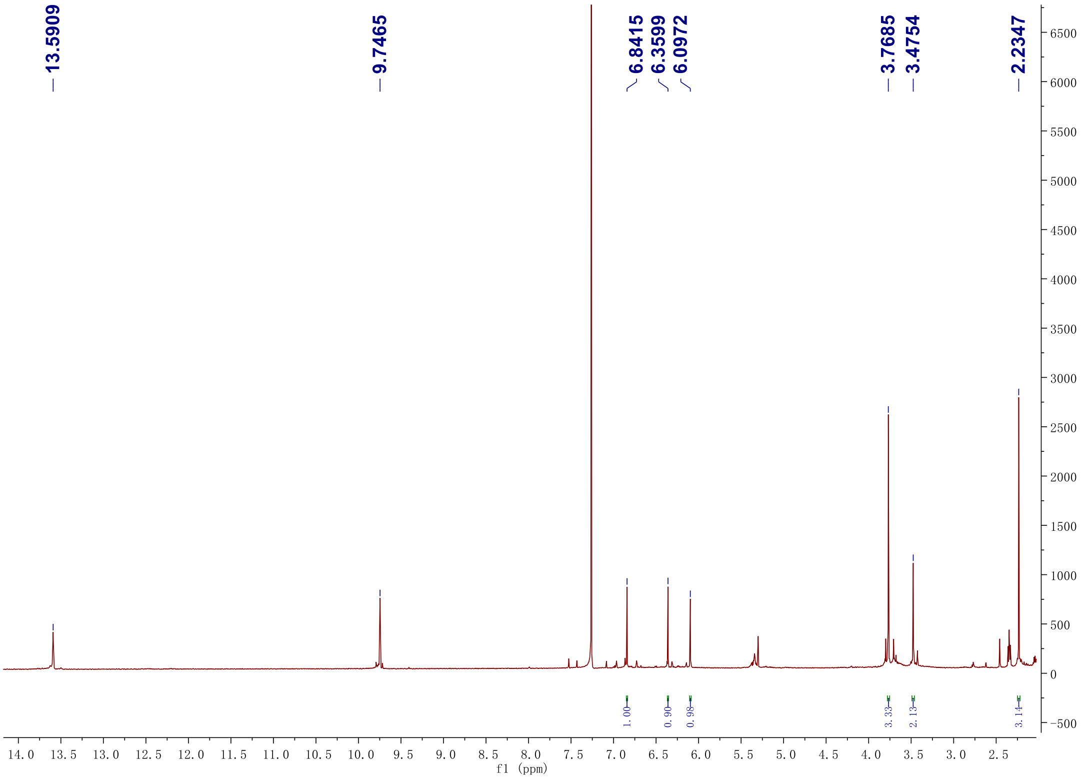

**Figure S40. $^{13}$C NMR spectrum (125 MHz) of 7 in chloroform-*d***

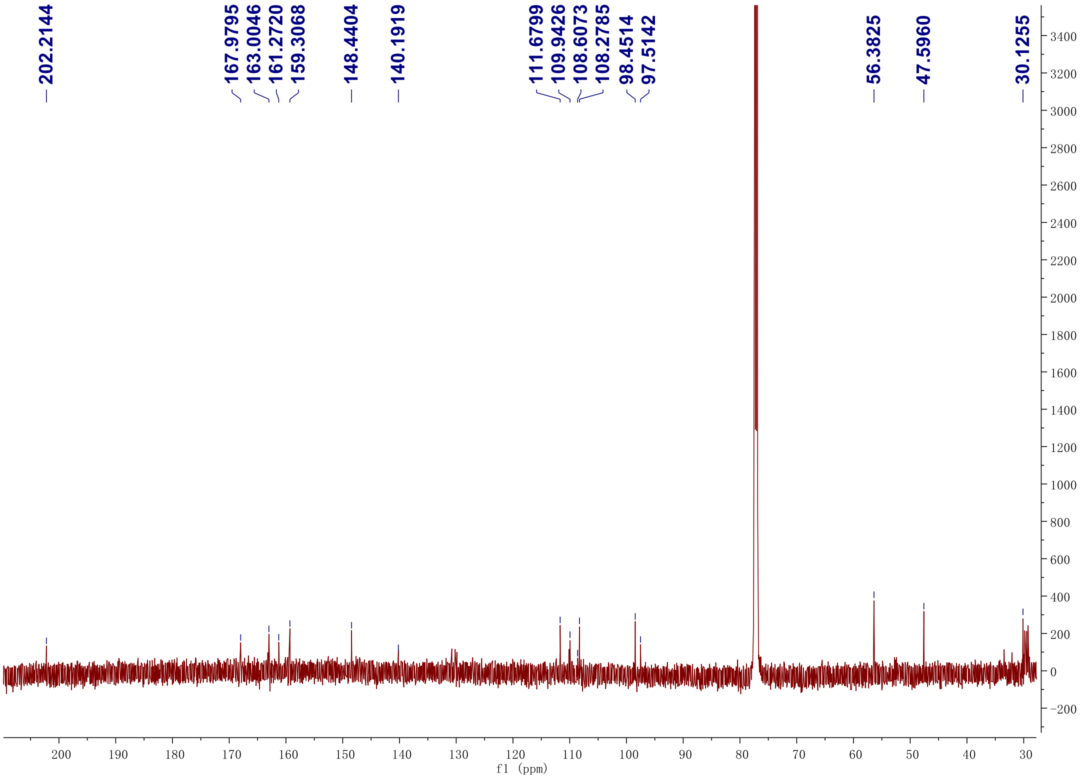

**Figure S41. DEPT-135 $^{13}$C NMR spectrum (125 MHz) of 7 in chloroform-*d***

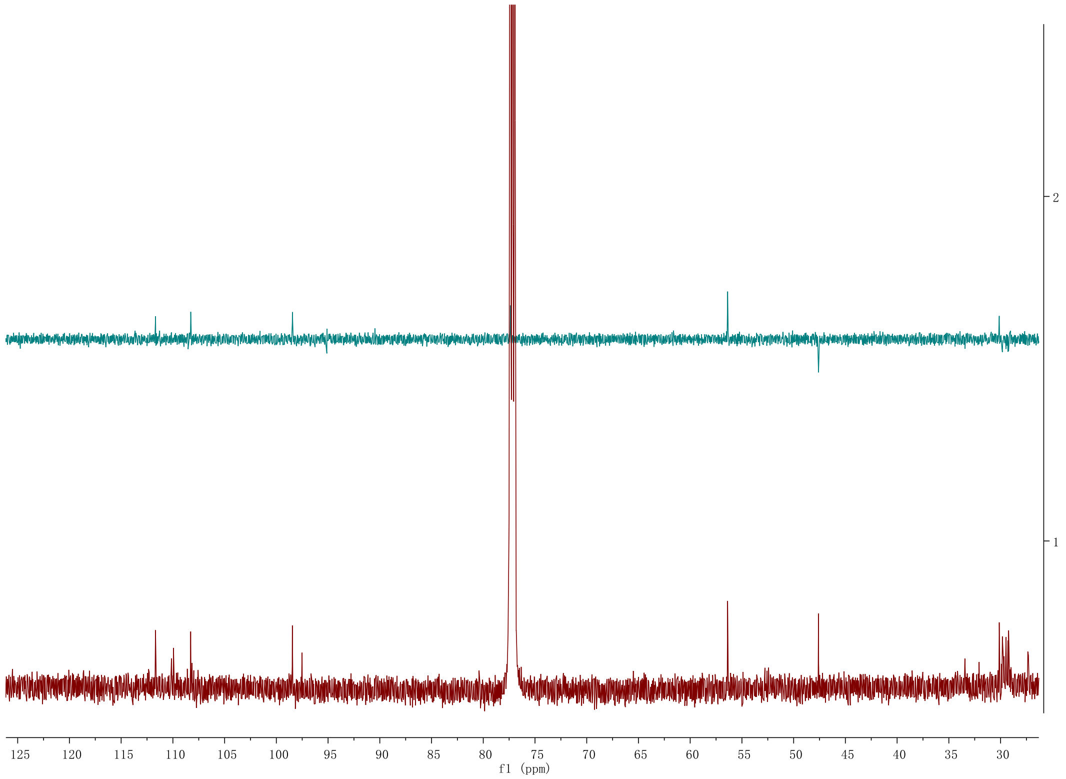

**Figure S42.** **$^1H$-$^1H$ gCOSY NMR spectrum (500 MHz) of 7 in chloroform-*d***

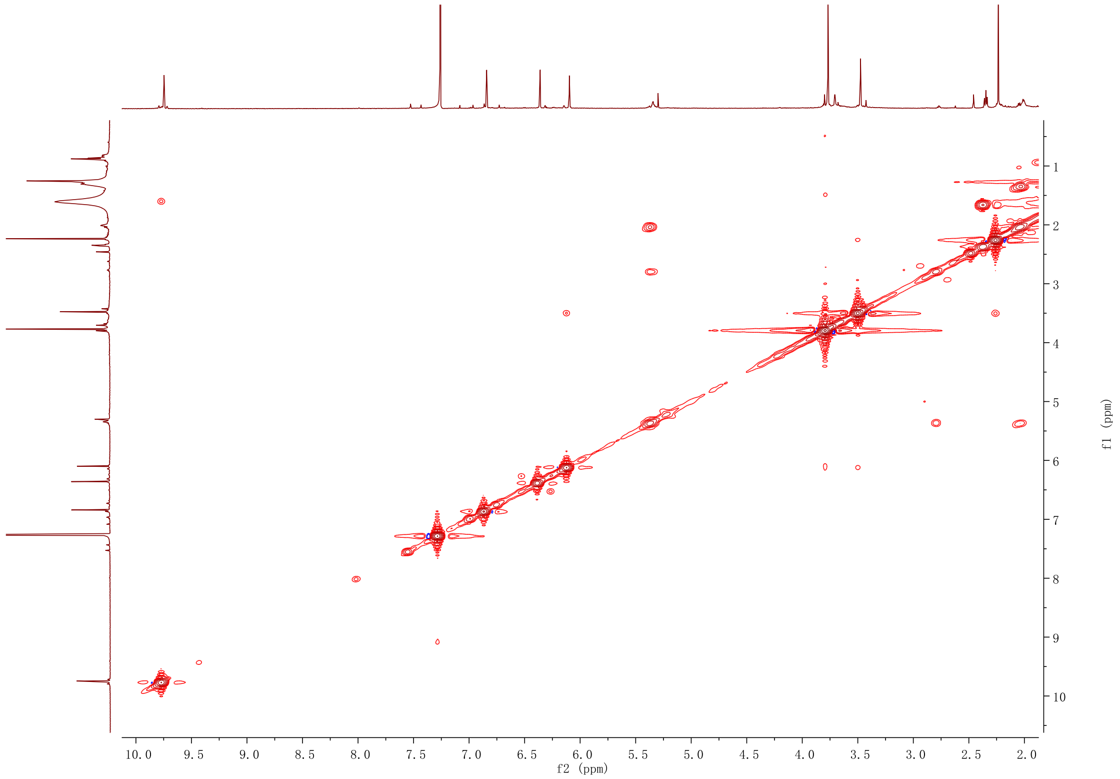

**Figure S43. HSQC NMR spectrum (500 MHz) of 7 in chloroform-*d***

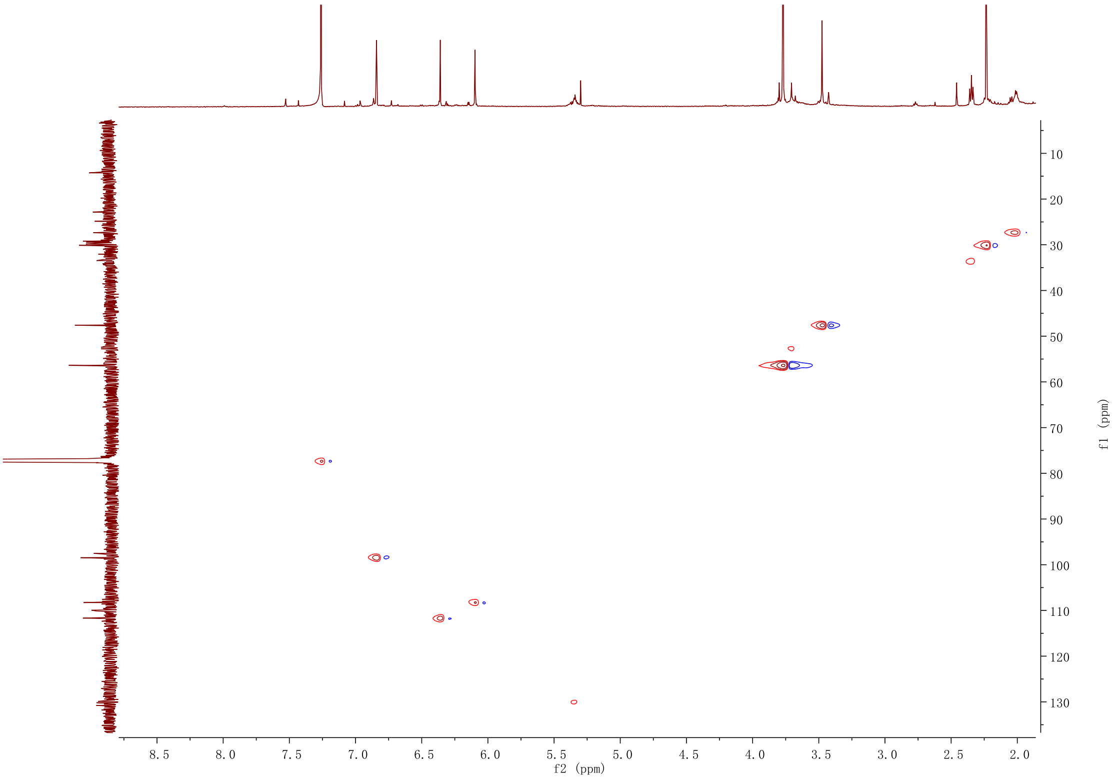

**Figure S44. HMBC NMR spectrum (500 MHz) of 7 in chloroform-*d***

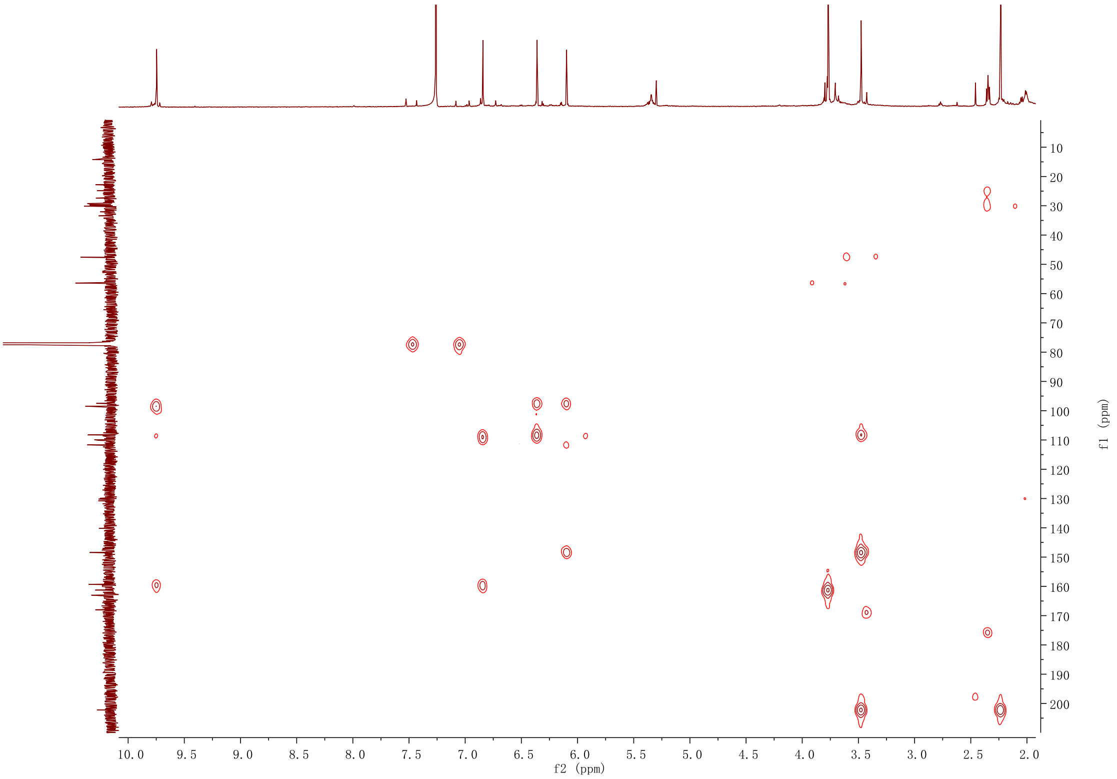

**Figure S45. [1]H NMR spectrum (600 MHz) of 8 in chloroform-*d***

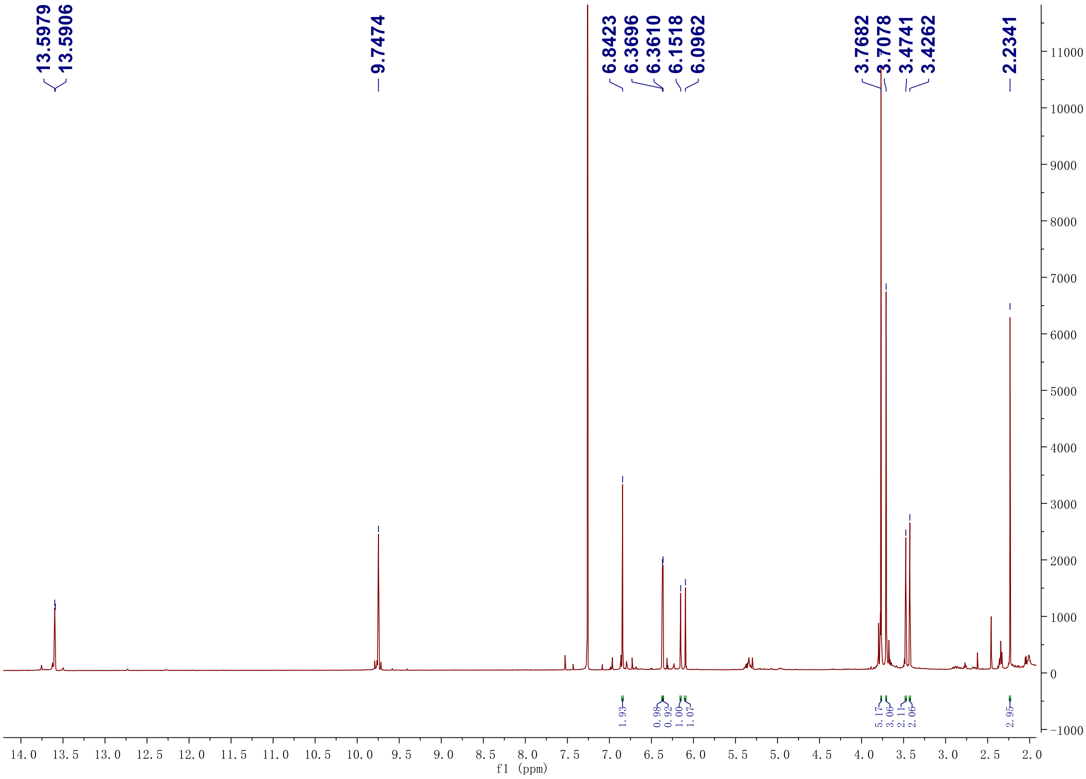

#### Figure S46. $^{13}$C NMR spectrum (150 MHz) of 8 in chloroform-*d*

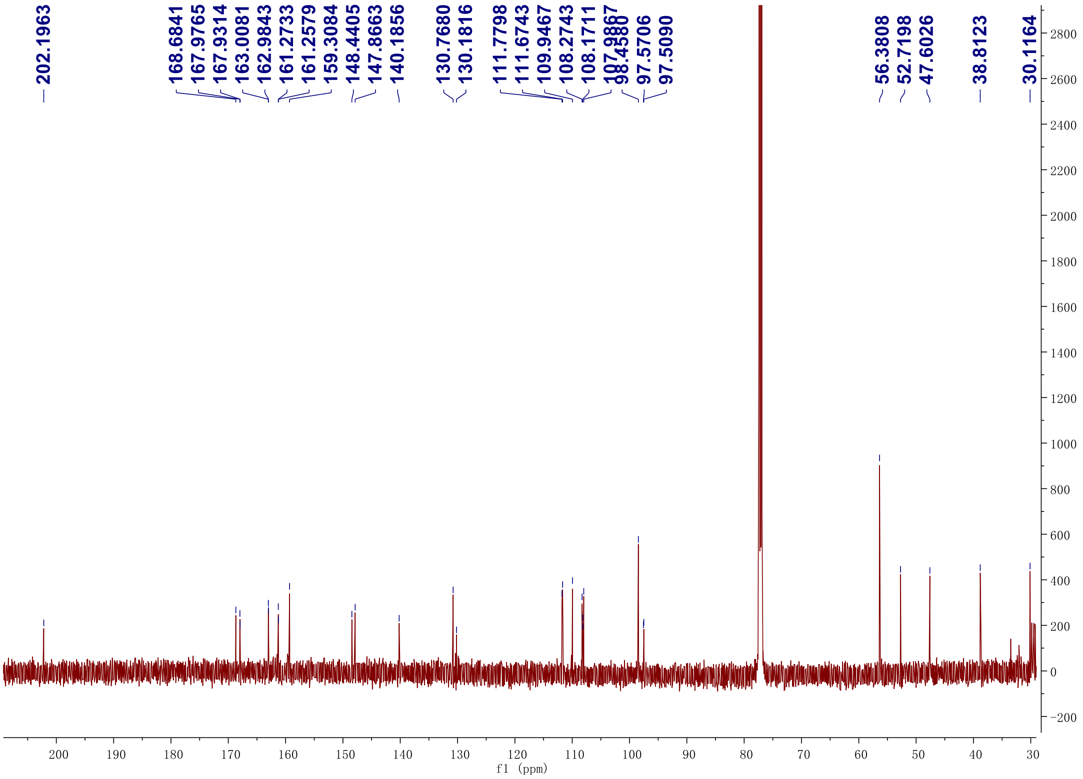

**Figure S47. DEPT-135 $^{13}$C NMR spectrum (150 MHz) of 8 in chloroform-*d***

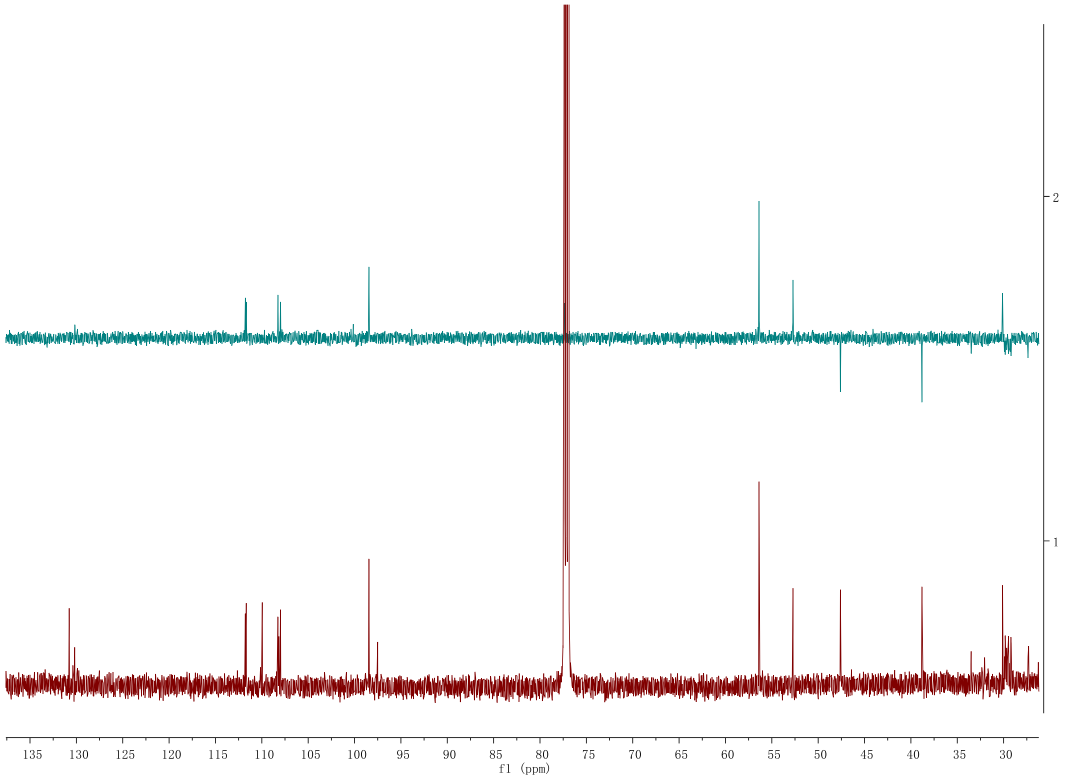

### Figure S48. $^1H$-$^1H$ gCOSY NMR spectrum (500 MHz) of 8 in chloroform-*d*

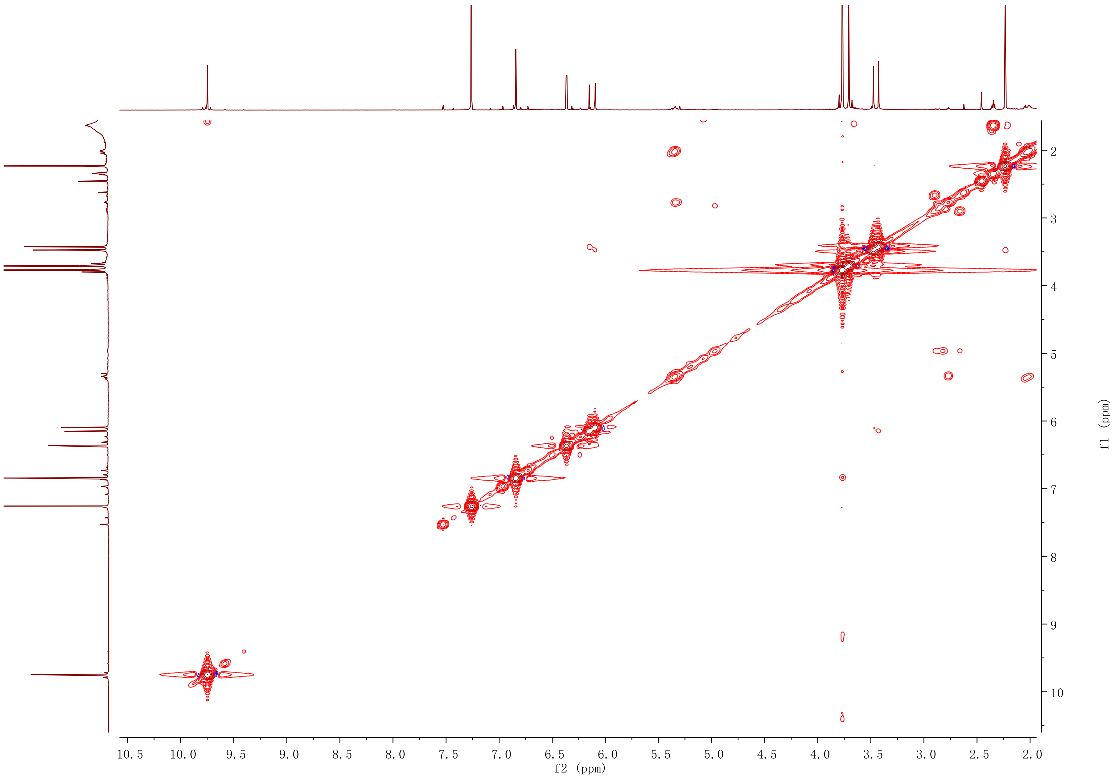

**Figure S49. HSQC NMR spectrum (600 MHz) of 8 in chloroform-*d***

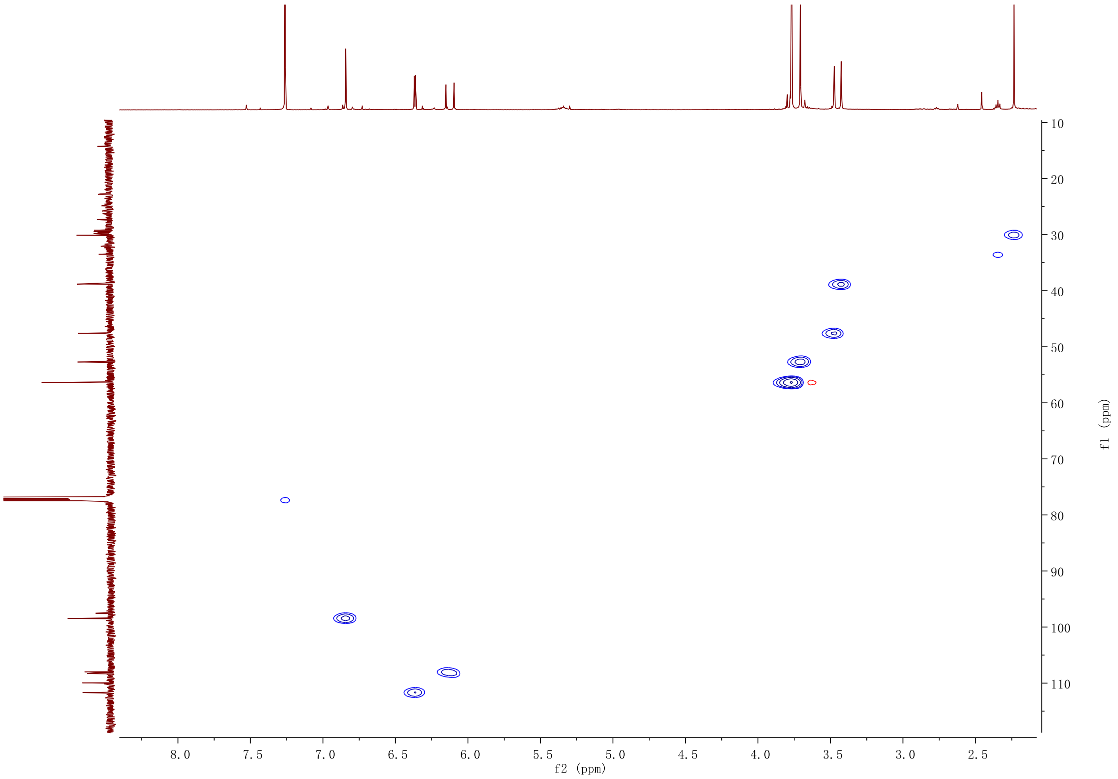

**Figure S50 HMBC NMR spectrum (600 MHz) of 8 in chloroform-*d***

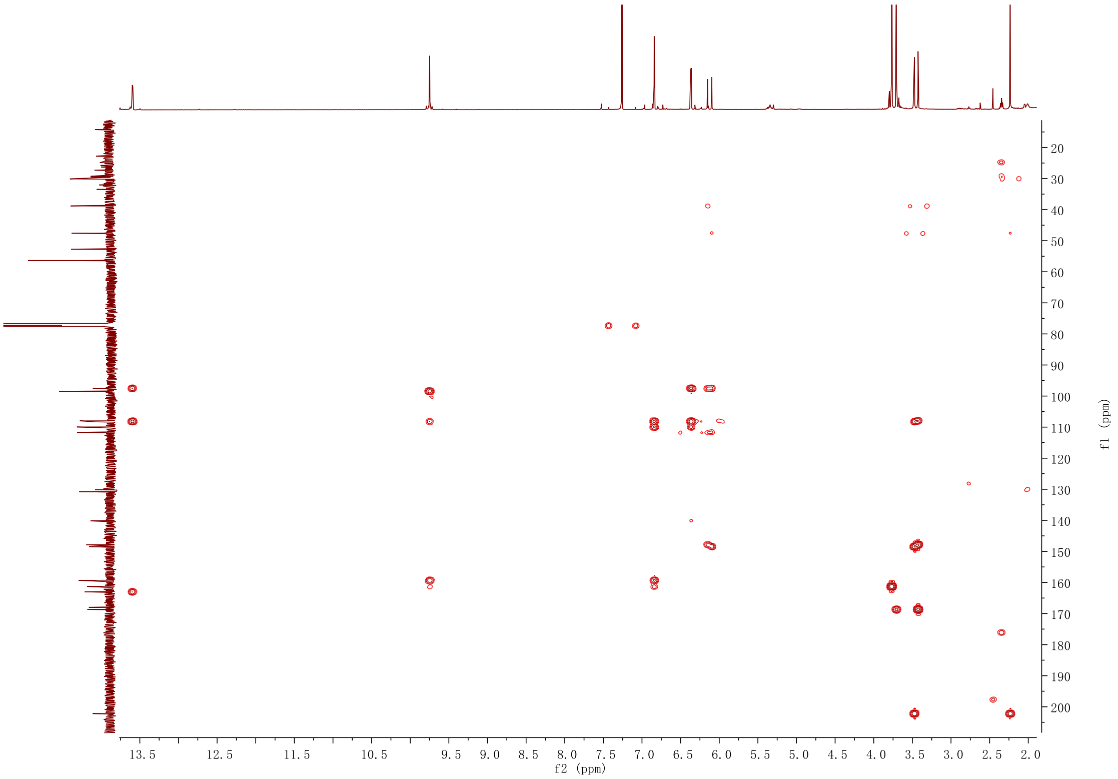

**Figure S51.** $^{1}H$ **NMR spectrum (500 MHz) of 9 in chloroform-*d***

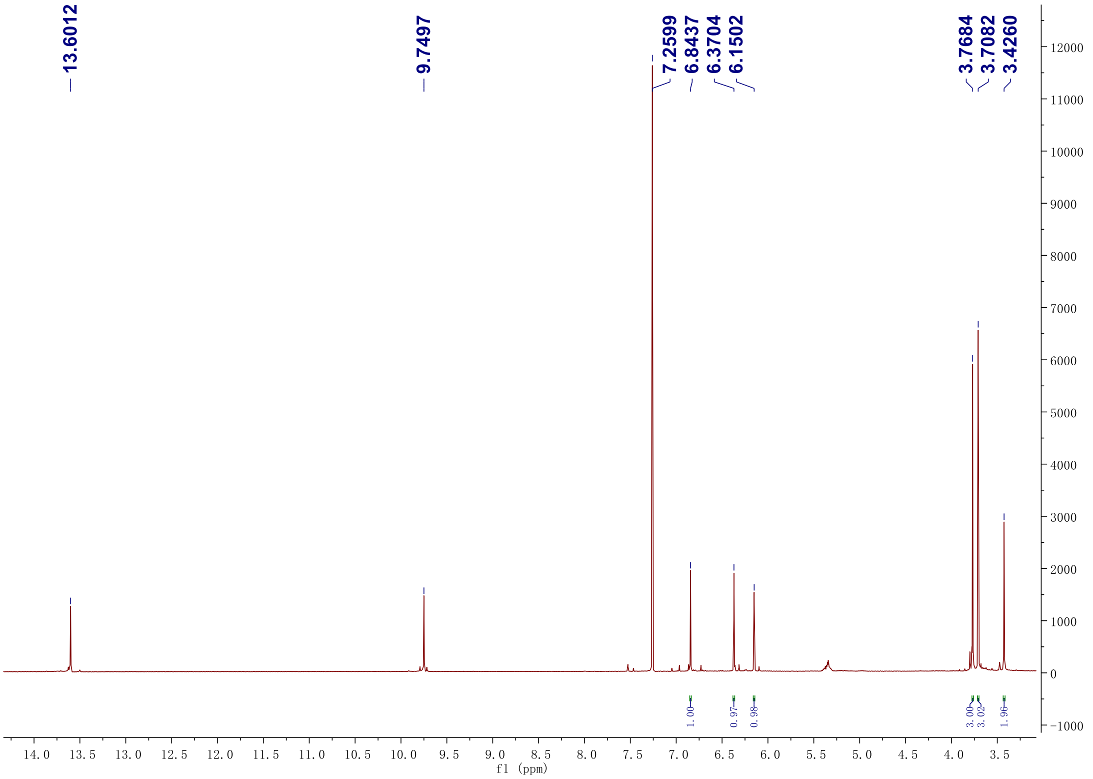

**Figure S52. $^{13}C$ NMR spectrum (125 MHz) of 9 in chloroform-*d***

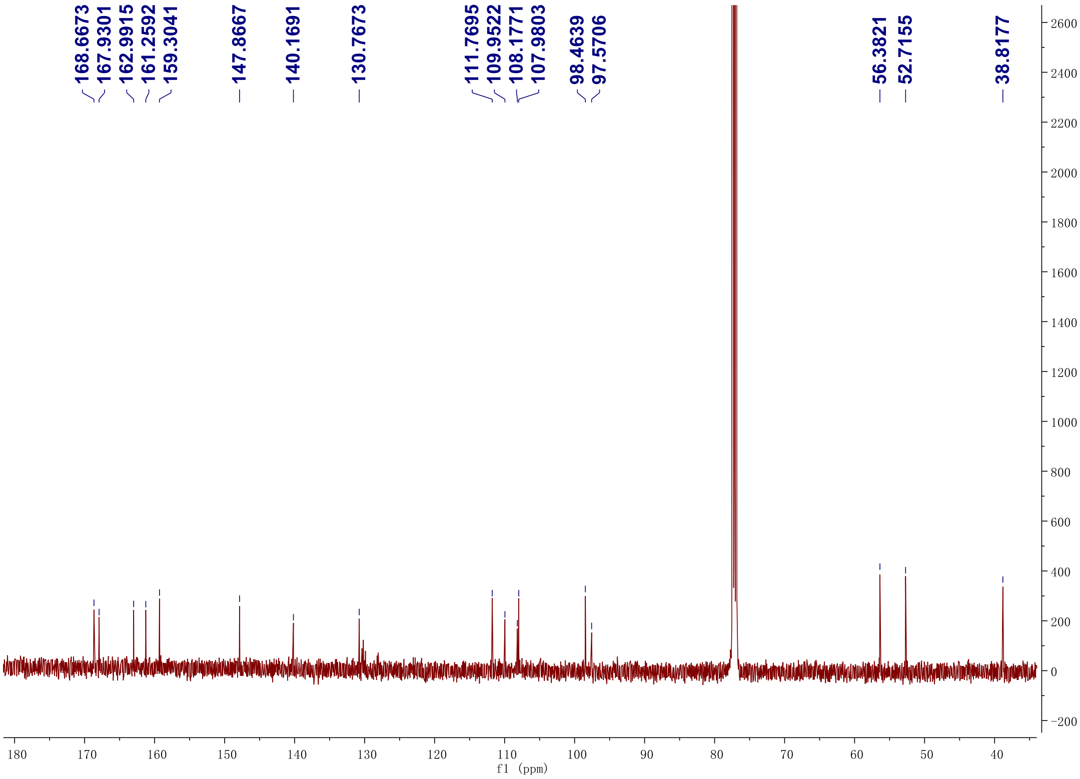

**Figure S53. DEPT-135 $^{13}$C NMR spectrum (125 MHz) of 9 in chloroform-*d***

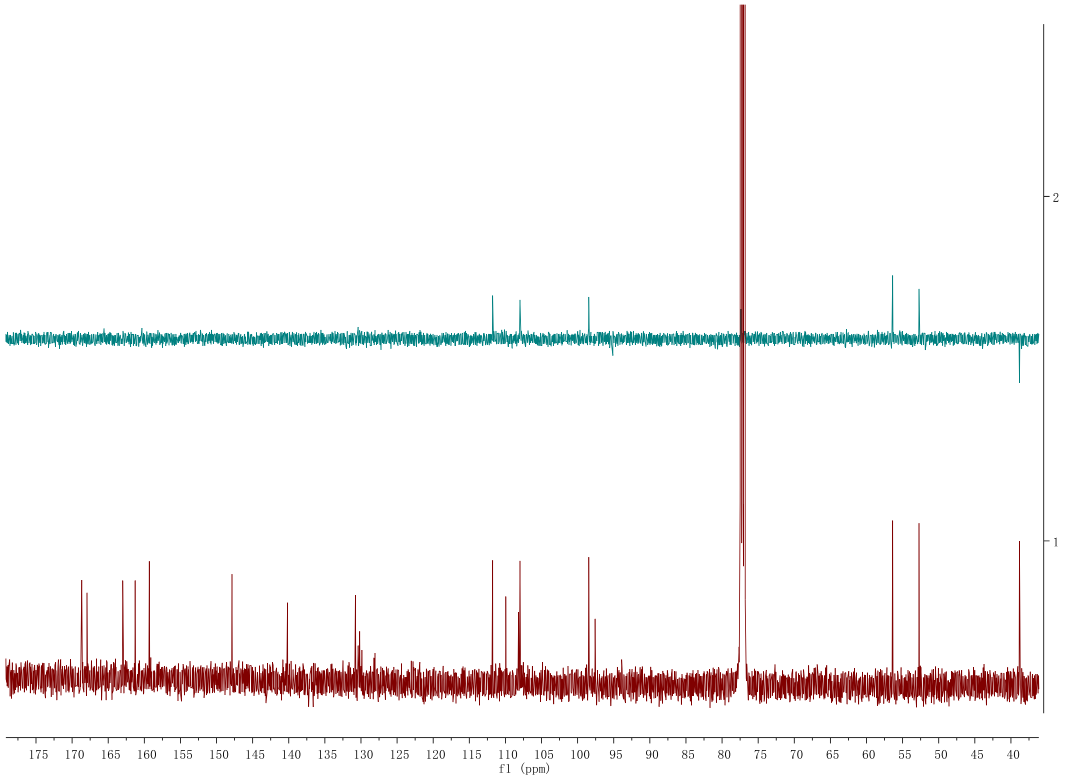

**Figure S54. $^1H$-$^1H$ gCOSY NMR spectrum (500 MHz) of 9 in chloroform-*d***

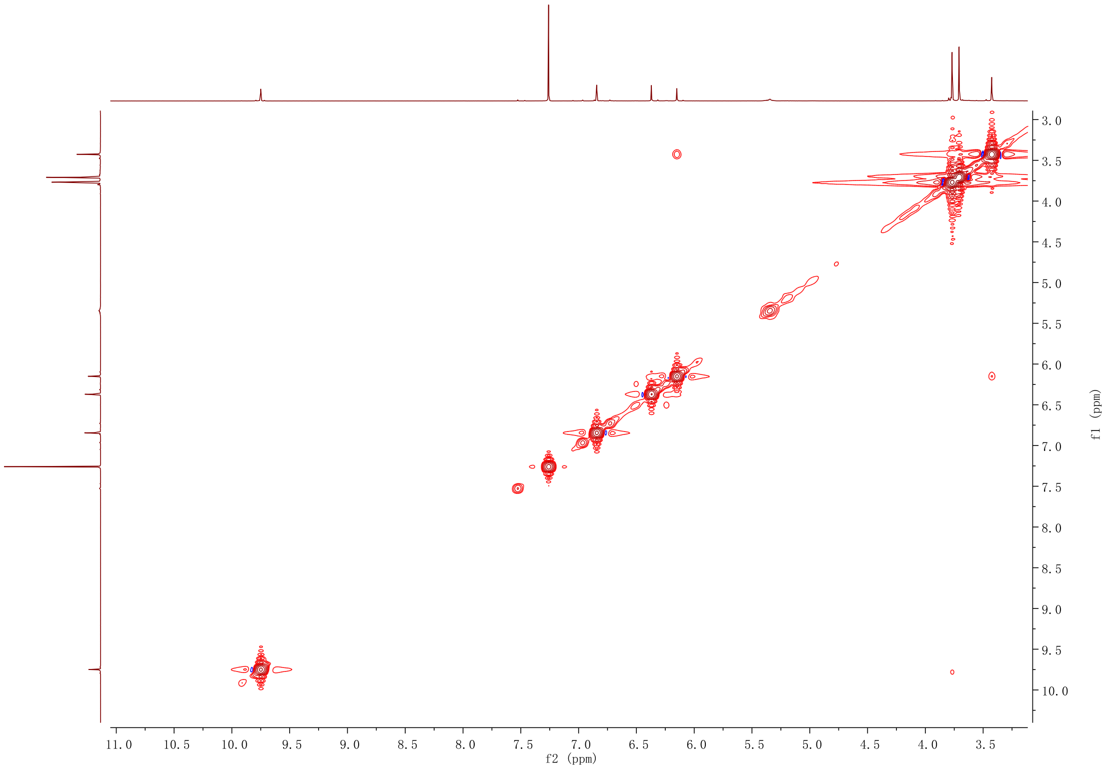

**Figure S55. HSQC NMR spectrum (500 MHz) of 9 in chloroform-*d***

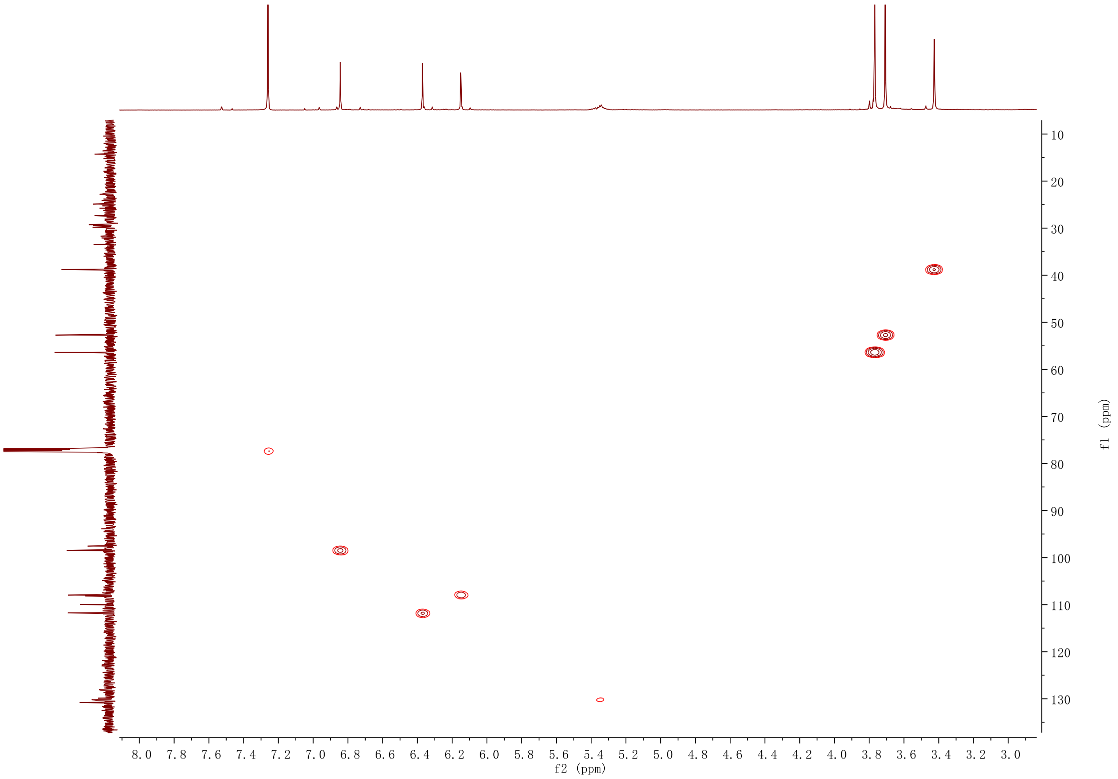

**Figure S56. HMBC NMR spectrum (500 MHz) of 9 in chloroform-*d***

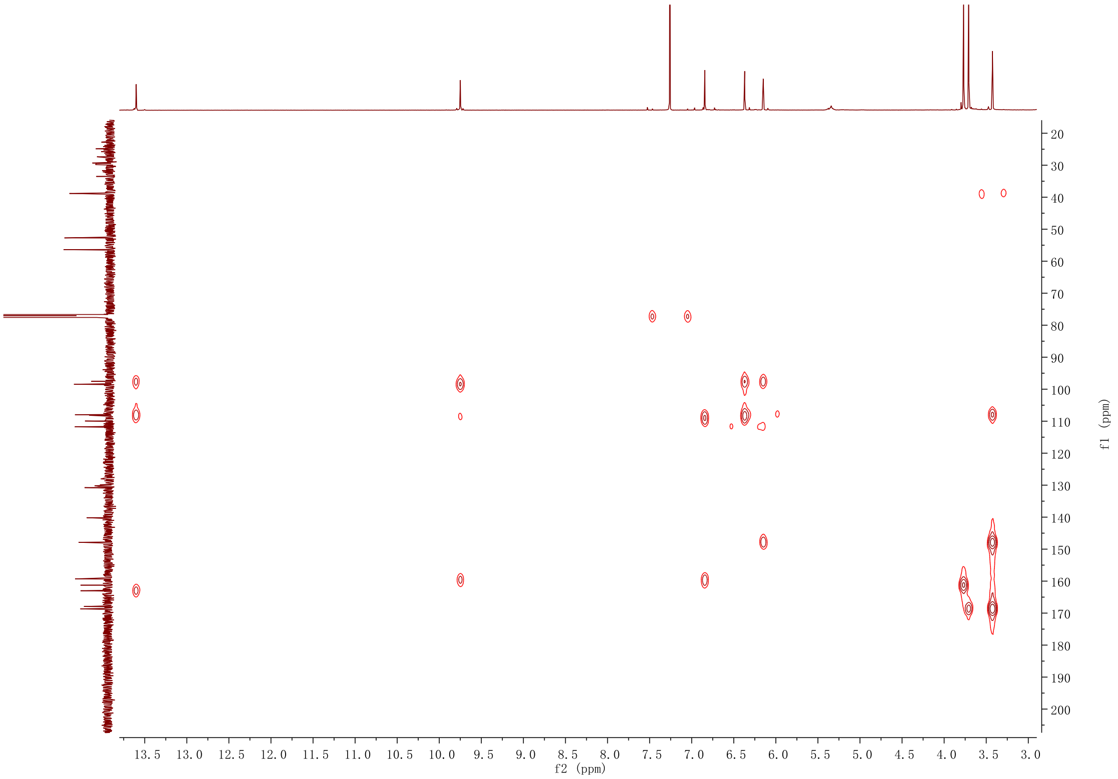
